# Robust inference and widespread genetic correlates from a large-scale genetic association study of human personality

**DOI:** 10.1101/2025.05.16.648988

**Authors:** Ted Schwaba, Margaret L. Clapp Sullivan, Wonuola A. Akingbuwa, Kerli Ilves, Peter T. Tanksley, Camille M. Williams, Yavor Dragostinov, Wangjingyi Liao, Lindsay S. Ackerman, Josephine C. M. Fealy, Gibran Hemani, Javier de la Fuente, Priya Gupta, Murray B. Stein, Joel Gelernter, Daniel F. Levey, Urmo Võsa, Liisi Ausmees, Anu Realo, Estonian Biobank Research Team, Mariliis Vaht, Jüri Allik, Tõnu Esko, René Mõttus, Uku Vainik, Gudrun A. Jonsdottir, Gudmar Thorleifsson, Árni Freyr Gunnarsson, Gyda Bjornsdottir, Thorgeir E. Thorgeirsson, Hreinn Stefansson, Kari Stefansson, Rosa Cheesman, Qi Qin, Elizabeth C. Corfield, Helga Ask, Fartein Ask Torvik, Eivind Ystrom, Martin Tesli, Dorret I. Boomsma, Eco J. C. de Geus, Jouke-Jan Hottenga, Dener Cardoso Melo, Harold Snieder, Catharina A. Hartman, Charley Xia, Archie Campbell, Michelle Luciano, Ian J. Deary, W. David Hill, Seon-Kyeong Jang, Scott I. Vrieze, Gonçalo Abecasis, Michelle K. Lupton, Brittany L Mitchell, Petra V. Viher, Lucía Colodro-Conde, Nicholas G. Martin, Sarah E. Medland, Eske M. Derks, Briar Wormington, Jaakko Kaprio, Karri Silventoinen, Teemu Palviainen, Agnieszka Gidziela, Kaili Rimfeld, Robert Plomin, Margherita Malanchini, Danielle M. Dick, Fazil Aliev, COGA Collaborators, The Spit for Science Working Group, Laura W. Wesseldijk, Fredrik Ullén, Miriam A. Mosing, Henry R. Kranzler, Yaira Nunez, Sarah Beck, Renato Polimanti, Tobias Edwards, Alexandros Giannelis, Emily A. Willoughby, James J. Lee, Matt McGue, Antonio Terracciano, Michele Marongiu, Edoardo Fiorillo, Francesco Cucca, Angelina R. Sutin, Peter J. van der Most, Albertine J. Oldehinkel, Tina Kretschmer, Andrey A. Shabalin, Anna R. Docherty, Robert F. Krueger, Colin D. Freilich, Binisha H. Mishra, Terho Lehtimäki, Olli T. Raitakari, Mika Kähönen, Aino Saarinen, Henrik Dobewall, Liisa Keltikangas-Järvinen, Klaus Berger, Marisol Herrera-Rivero, Fabian Streit, Swapnil Awasthi, Stephanie H. Witt, Johanna Tuhkanen, Katri Räikkönen, Johan G. Eriksson, Jari Lahti, Gail Davies, Paul Redmond, Adele Taylor, Janie Corley, Tom C. Russ, Marina Ciullo, Teresa Nutile, Jun Ding, Yong Qian, Toshiko Tanaka, Luigi Ferrucci, Lea Zillich, Lea Sirignano, K. Paige Harden, Erhan Genç, Patrick D. Gajewski, Stephan Getzmann, Christoph Fraenz, Javier E. Schneider Peñate, Stefanie Lis, Alisha S. M. Hall, Christian Schmahl, Sabine C. Herpertz, Abdel Abdellaoui, Michel G. Nivard, Elliot M. Tucker-Drob

## Abstract

Personality traits describe stable differences in how individuals think, feel, and behave and how they interact with and experience their social and physical environments. We assemble data from 46 cohorts including 611K-1.14M participants with European-like and African-like genomes for genome-wide association studies (GWAS) of the Big Five personality traits (extraversion, agreeableness, conscientiousness, neuroticism, and openness to experience), and data from 51K participants for within-family GWAS. We identify 1,257 lead genetic variants associated with personality, including 823 novel variants. Common genetic variants explain 4.8%-9.3% of the variance in each trait, and 10.5%-16.2% accounting for measurement unreliability. Genetic effects on personality are highly consistent across geography, reporter (self vs. close other), age group, and measurement instrument, and we find minimal spousal assortment for personality in recent history. In stark contrast to many other social and behavioral traits, within-family GWAS and polygenic index analyses indicate little to no shared environmental confounding in genetic associations with personality. Polygenic prediction, genetic correlation, and Mendelian randomization analyses indicate that personality genetics have widespread, potentially causal associations with a wide range of consequential behaviors and life outcomes. The genetic architecture of personality is robust and fundamental to being a human.

## Introduction

Personality is defined by relatively stable patterns of thinking, feeling, and behaving that vary across individuals (Roberts and Yoon, 2022). Decades of research have demonstrated that human personality can be meaningfully and efficiently summarized by five broad trait factors known as the *Big Five*: Extraversion (which encompasses traits such as sociability, assertiveness, and energy level), agreeableness (compassion, respectfulness, and trust), conscientiousness (organization, productiveness, and responsibility), neuroticism (anxiety, depression, and volatility) and openness to experience (imagination, curiosity, and aesthetic sensitivity) (John et al., 2008; Matthews et al., 2009; Soto & John, 2017). The Big Five predict academic performance and educational attainment, labor market outcomes, and career choices, at comparable levels to cognitive ability and socioeconomic status (Anni et al., 2024; Borghans et al., 2016; Bucher et al., 2019; Deary et al., 2010; Hampson, 2012; Roberts et al., 2007; Soto, 2019). They relate to a wide range of health behaviors, such as eating habits, drug use, and physical activity (Bogg & Roberts, 2004; De Moor et al., 2006; Hampson et al., 2007; Willroth et al., 2023) and to disease burden (Yoneda et al., 2023) and longevity (Beck & Jackson, 2022; Graham et al., 2017). The Big Five are so closely linked to mental health that modern taxonomies of psychopathology are foundationally informed by patterns of personality trait variation (Kotov et al., 2010; Widiger et al., 2019). And they predict relationships and social behavior (Antonoplis & John, 2017; Landis, 2016), residential mobility (Jokela, 2020, Jokela et al., 2015), and political preferences (Duckitt & Sibley, 2010). Given the fundamental relevance of personality to core phenomena studied across scientific disciplines, understanding the molecular genetics of personality is crucially important for developing biopsychosocial models of both human behavior and life outcomes.

Here we report results from the Revived Genomics of Personality Consortium (ReGPC), a collaboration across 46 cohorts incorporating between 611K and 1.14M participants with European-like (EUR) and African-like (AFR) genomes (National Academies of Sciences, Engineering, and Medicine, 2023) per Big Five trait. This effort represents a substantial boost in statistical power for genetic discovery over past personality GWAS efforts, increasing the number of significant loci relative to most recent work (Gupta et al., 2024) from 3 to 131 for conscientiousness, 4 to 39 for agreeableness, 8 to 126 for openness to experience, 14 to 258 for extraversion, and 224 to 703 for neuroticism. Though personality traits often vary on average across groupings of people (Bleidorn et al., 2022; Ebert et al., 2022), we find that the genetic etiology of each trait is highly consistent across geography, reporter perspective (self vs. close other), age, and measurement instrument. We empirically confirm the strong relationship between a measurement’s internal consistency and its heritability; after accounting for measurement error, heritability estimates for the Big Five are substantially greater. We report results of within-family GWAS of the Big Five in over 50K participants, which allow us to parse genetic effects from environmental confounds. We find that the genetic architecture of the Big Five is only minimally confounded by population stratification, dynastic, and assortative mating effects that commonly bias GWAS of other behavioral traits (Howe et al., 2022; Tan et al., 2024; Young et al., 2022). Polygenic indices (PGIs) constructed from each primary GWAS robustly predict personality in five independent cohorts, with no identifiable reduction in predictive accuracy when compared within families. Across the Big Five, common variant heritability estimates are largely indistinguishable when estimated using within-family vs. population GWAS, and population-level and within-family genetic effects are highly genetically correlated (mean *r*_*g*_= .86). We characterize the functional genomics of each of the Big Five and provide evidence of negative selection on the genetics of personality. Finally, we extensively characterize personality’s wide-ranging genetic correlates and consequences across social relevant behaviors and life outcomes ranging from psychopathology and physical health to research participation and residential mobility, highlighting a central role of personality genetics in understanding the human experience.

## Results

### Population-level Association Meta-Analysis

We performed population-level GWAS meta-analysis of each Big Five trait in genetic-similarity-stratified and genetic-similarity-combined cohorts of EUR (k = 46) and AFR (k = 10) participants. In each cohort analysts conducted GWAS of each available Big Five measurement instrument using a standard protocol across the 22 autosomes and, when available, the X-chromosome (**Supplementary Tables S1-S3)**. We jointly analyzed results using inverse-variance weighted meta-analysis, resulting in pooled associations for ∼ 10 million Single Nucleotide Polymorphisms (SNPs) for each Big Five trait.

We identified 1,257 approximately independent (*r*^2^ < .10 within a 250kb range) genome-wide significant (GWS; *p <* 5×10^−8^) SNPs associated with the Big Five traits, an increase of 3 to 43 times over the most recent findings for each trait (**Table 1; Supplementary Tables S4-S11**). Of these lead SNPs, 823 (65%) were found in novel loci not previously associated with variation in the respective trait. Manhattan plots of these results are displayed in **Figure 1 A-E**. Demonstrating the independence between the Big Five, 82% of genomic loci containing a GWS SNP were associated with only a single trait (**Supplementary Table S12**), and the average absolute genetic correlation among Big Five traits among EUR participants was .19 (**Figure 1 F)**, which is similar to estimates of phenotypic correlations (Soto & John, 2017).

**Table 1.**
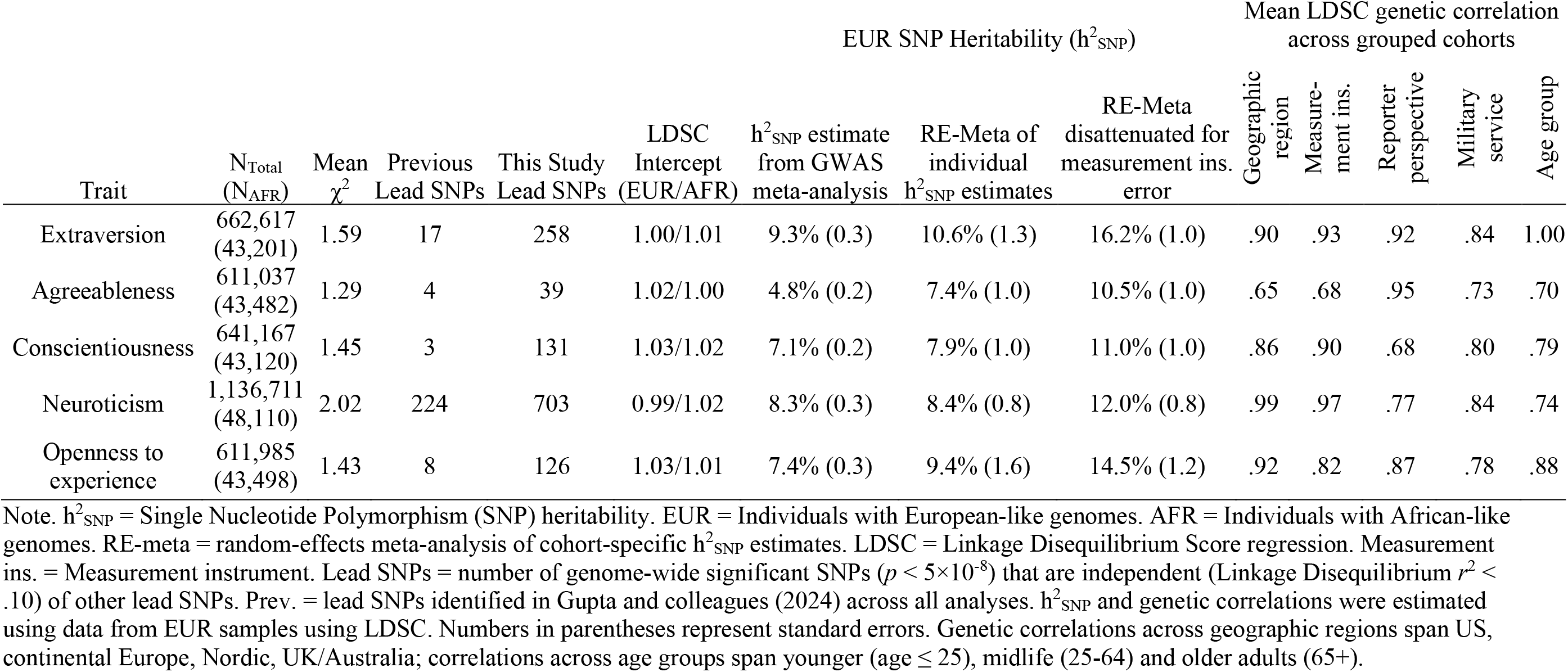
Summary of Population-Level Genome-Wide Association Study Results.

**Figure 1.**
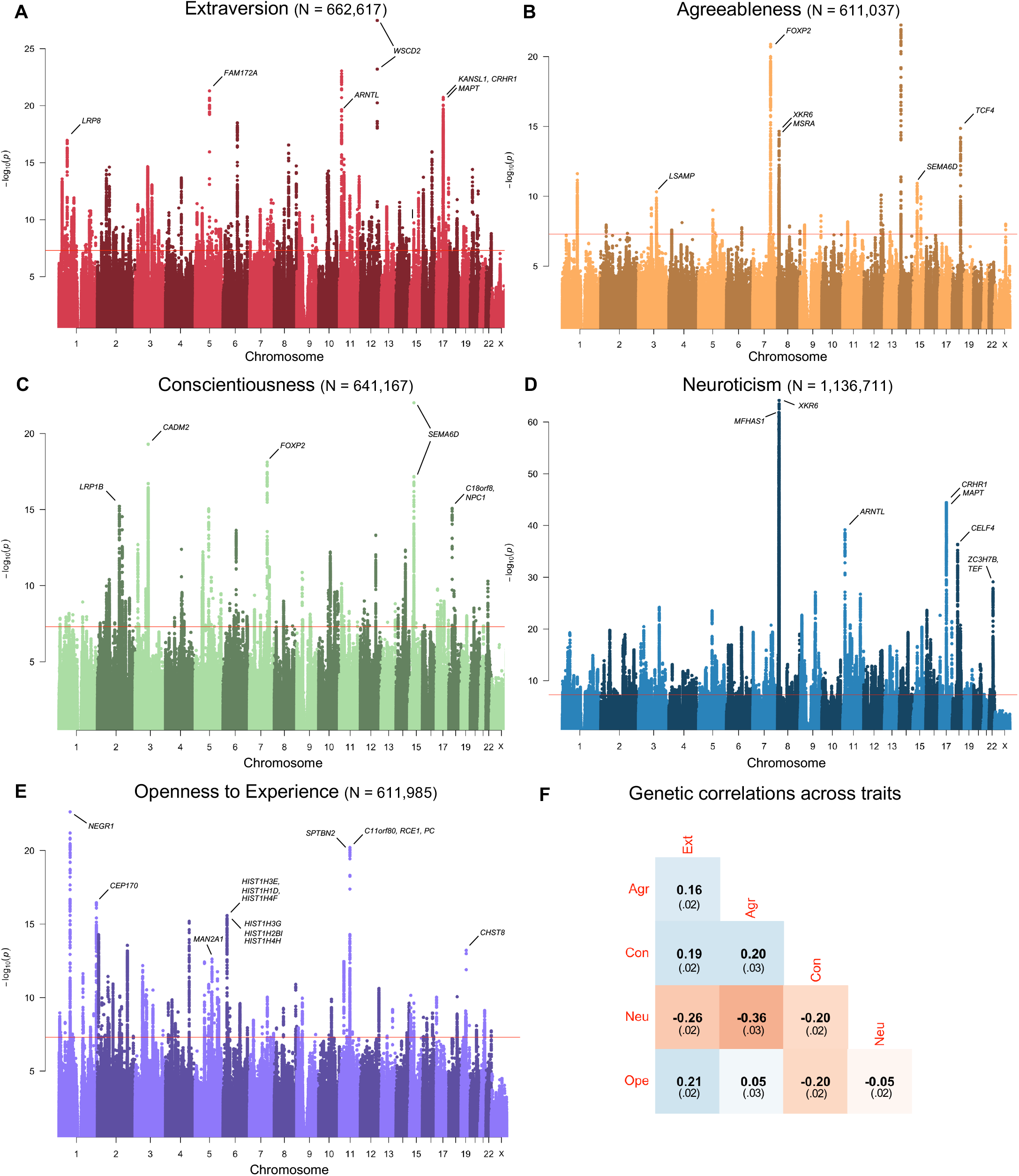
Manhattan plots of Genome-Wide Associations for each Big Five personality trait (A-E) and genetic correlations across traits (F). Genome wide significance is denoted with a red line at *p* = 5×10^−8^. A selection of genes containing or nearby the most significant lead SNPs are annotated in each panel (**Supplementary Tables S23-S27**). Genetic correlations in panel F were estimated among EUR participants using LDSC; standard errors are in parentheses.

Genetic variants with previously reported significant Big Five associations evinced a high replication rate: 207 of 246 (84%) lead SNPs identified in the most recent Big Five GWAS, which partially overlapped with the present GWAS sample, were found in the same locus as a significant variant in this study (Gupta et al., 2024; **Supplementary Table S13**). Individual SNP effects were extremely small, reflecting personality’s highly distributed genetic architecture. For example, the median association estimate among extraversion lead SNPs, corrected for the winner’s curse (Forde et al., 2023), is .009 SD per effect allele (corresponding to scoring at the 50.35^th^ vs 50^th^ percentile in extraversion). Genetic effects on personality were especially strongly dispersed, as indicated by Linkage Disequilibrium (LD) Fourth Moments Regression (O’Connor et al., 2019). On average, the estimated effective number of independently associated common SNPs for each trait was 16,180 (**Supplementary Table S14**); Extraversion, agreeableness, conscientiousness, and openness to experience were each estimated to be as or more polygenic than 31 of the 32 biobehavioral phenotypes compared in O’Connor and colleagues (2019), including schizophrenia liability and smoking status. We present additional GWAS results, including genetic ancestry-stratified analyses, **Supplementary Figures S1-S10** and **Supplementary Tables S15-S16.**

### Characterizing Common-Variant Heritability

Among EUR participants, SNP heritability (h^2^_SNP_) estimated for the Big Five traits using LD Score Regression (LDSC; Bulik-Sullivan et al., 2015) ranged from 4.8% (SE = 0.2%) for agreeableness to 9.3% (SE = 0.3%) for extraversion (**Table 1; Supplementary Table S4**). Importantly, these SNP heritability estimates from GWAS meta-analysis incorporate genetic effects that are consistent across contributing cohorts. To allow for variability in genetic effects across cohorts, we conducted a random-effects meta-analysis of cohort-specific h^2^_SNP_estimates, which indicated an average h^2^_SNP_of 8.6% (SE = 0.6%) across traits (ranging from 7.4% for agreeableness to 10.6% for extraversion; **Table 1**; **Supplementary Table S17**), with significant variability across cohorts (mean τ = 3.6%).

h^2^_SNP_is expected to vary as an inverse function of the measurement error in the measurement instrument used (Spearman, 1910; Tucker-Drob, 2017). We confirmed this to be the case using a weighted meta-regression in which we regressed cohort-level h^2^_SNP_on Cronbach’s alpha (which indexes a measure’s internal consistency on a scale of 0-1), allowing for random intercepts to account for nesting of trait estimates within cohorts. Across all traits, we found that personality measures with greater internal consistency tended to be much more heritable (*b* = 6.7%, SE = .7%; **Supplementary Figure S11**). In this analysis, the expected h^2^_SNP_for an error-free personality measure ranged from 10.5% for agreeableness (SE = 1.0%) to 16.2% for extraversion (SE = 1.0%; **Table 1**).

Biological follow-up of GWAS signal using MAGMA (de Leeuw et al., 2015) indicated that enriched gene-sets intersected across the Big Five (mean enrichment rank-order ρ = .72; **Supplementary Figures S12-S17**), providing evidence for trait-overlapping molecular and cellular systems in personality neurobiology despite only modest genetic correlations (**Figure 1 panel F**). Consistent with theories of personality development that emphasize the prefrontal cortex (Casey & Caudle, 2013; Romer et al., 2017), genetic associations for each Big Five trait except agreeableness were enriched in genes expressed in the prefrontal cortex (among these, our top lead SNPs implicate *RCE1, FOXP2*, and *SEMA6D*, indicated in **Figure 1**; **Supplementary Tables S18-S27**). Each of the Big Five demonstrated strong enrichment in protein truncating variant intolerant gene-sets specifically expressed in neurons (e.g., *ARNTL, TCF4, NEGR1*; **Figure 1**). This pattern, which has also been found for psychiatric disorders and cognitive function (Grotzinger et al., 2022a; Grotzinger et al., 2025), suggests personality-relevant variants are under negative selection, which appears inconsistent with evolutionary theories that posit balancing selection mechanisms as responsible for maintaining genetic variation in personality (Penke & Jokela, 2015). Neurobiological theories of the Big Five posit dopaminergic etiology to variation in openness to experience and extraversion, and serotonergic etiology to variation in conscientiousness, neuroticism, and agreeableness (Carver & Miller, 2006; DeYoung et al., 2021; Depue & Collins, 1999). Based on sets of genes differentially expressed in specific brain cell types, both in mouse and human post-mortem tissues, we found little support for these hypotheses across trait-stratified tests: genes differentially expressed in several types of dopaminergic and serotonergic neurons were in some cases (nominally) significantly enriched for specific outcomes but neither the effect size or p-value placed them among the most enriched neuron types (**Supplementary Tables S18**-**S22**), suggesting their prominence in the literature relative to other types of neurons is not warranted.

To further characterize the generalizability of genetic effects on personality, we examined the concordance of genetic signal across geography, age, veteran status, measurement instrument, and reporter perspective. We clustered cohorts by these grouping variables, re-estimated GWAS meta-analyses within balanced subgroups, and estimated genetic correlations between subgroups using LDSC (**Table 1; Supplementary Figures S18-S22**). We found that genetic effects were highly similar across four western country clusters (United States, Continental Europe, Nordic, and United Kingdom/Australia, mean *r*_*g*_= .86), three age groups (young [≤25], middle [25-64] and older [65+], mean *r*_*g*_= .80), between Million Veteran Program participants and other cohorts (mean *r*_g_ = .80), and across four personality measurement instruments (mean *r*_g_ = .89). Additional characterization of genetic architecture across measurement instruments using Genomic Structural Equation Modeling (Grotzinger et al., 2019) confirmed that genetic effects plausibly operate at the level of instrument-general latent factors, with only one locus (associated with both higher extraversion and lower neuroticism) demonstrating significantly heterogenous effects across measurement instruments (**Supplementary Tables S28-S30, Supplementary Figures S23-S29**). Notably, correlations across geography and measurement instrument were lower for agreeableness than the other Big Five (**Table 1**). This greater heterogeneity explains in part why agreeableness exhibited lower heritability than other traits in the combined meta-analytic GWAS: its genetic effects are less consistent across cohorts. In the Estonian Biobank, where participants’ personality was assessed both by their self-report (N = 73,983) and reports of close others (N = 20,269), we found a high degree of genetic overlap between rater perspectives (mean *r*_g_ = .84), providing strong evidence that the genetic architecture of personality is not a simple epiphenomenon of self-perception.

### Polygenic Prediction of Personality

Past efforts to predict personality from polygenic indices (PGIs) have been hampered by low power of the discovery GWAS used to estimate prediction weights (Becker et al., 2021). Here, we capitalized on the very large sample sizes of our EUR discovery GWAS to estimate weights for PGIs using SBayesR (Lloyd-Jones et al., 2019). PGIs were then tested in five independent cohorts (holding the cohort out from discovery GWAS when there was sample overlap). In all five cohorts, PGIs predicted significant additive variance in their respective phenotypic personality trait score among EUR participants, after controlling for sex, age, age^2^, and 10 ancestral principal components (extraversion M_β_ = .19, range = [.16, .23]; agreeableness M_β_ = .10, range = [.07, .14]; conscientiousness M_β_ = .16, range = [.12, .18]; neuroticism M_β_ = .16, range = [. 10, .18]; openness to experience Mβ = .17, range = [.15, .23]; **Figure 2A**). These estimates approximate mean R^2^ values ranging from 1.0% for agreeableness to 3.6%, for extraversion. These estimates represent an improvement in prediction over past research (Gupta et al., 2024) for all traits besides agreeableness. Further, highly consistent estimates across cohorts indicates a limited role of birth year, nationality, and ascertainment method in prediction accuracy.

**Figure 2.**
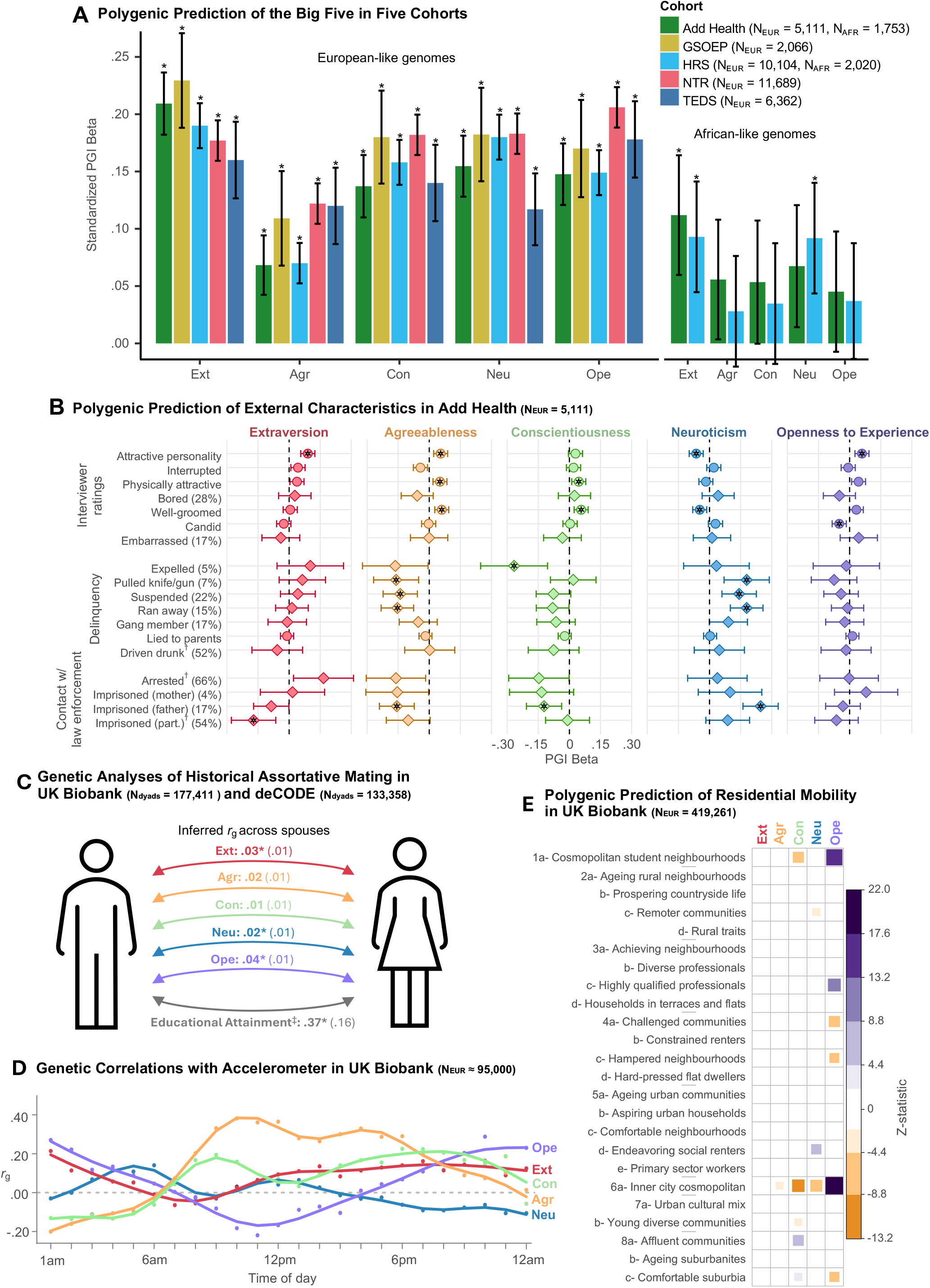
Polygenic prediction and additional genetic associations. Ext = Extraversion. Agr = Agreeableness. Con = Conscientiousness. Neu = Neuroticism. Ope = Openness to Experience. Panel A depicts standardized prediction of phenotypic personality trait scores among EUR and AFR participants in five cohorts using PGI weights derived from the EUR population-level GWAS. In panel B, betas for continuous traits are standardized and have point estimates depicted with circles, and betas for binary traits are logistic betas and have point estimates depicted with diamonds; percentage affirmative responses are noted in parentheses. † = questions asked only among a subgroup of participants (see Supplementary Text). Panel C depicts correlations between PGIs for each pair of family members. ‡ = Educational Attainment estimates come from Torvik et al. (2022), estimated across spouses in the MoBa cohort. In panel D, accelerometer activity indexes physical movement across a 24-hour period; see Grotzinger et al. (2022b) for complete details. Panel E depicts associations between PGI and census area of residence at age 50+, controlling for birthplace in addition to sex, age, array, and genetic principal components. Only associations that are significant at *p* < .01 after application of Benjamini-Hochberg false-discovery rate correction and sign-concordant within sibship are plotted. In all panels, error bars depict 95% confidence intervals. * = *p* < .01.

We also predicted personality traits among AFR participants in the Add Health and HRS cohorts, again using weights constructed from the highly powered EUR GWAS. PGI prediction is expected to substantially decrease when there are differences in genetic ancestry between discovery and target samples (Wang et al., 2023). Nevertheless, PGIs significantly predicted extraversion across both cohorts, and neuroticism in HRS (*p* < .01; **Figure 2A**), providing the first evidence for significant PGI prediction of personality traits beyond EUR individuals.

### Associations with Socially Relevant Behaviors and Important Life Outcomes

Building on epidemiological and longitudinal research that has demonstrated the breadth of personality’s associations, we quantified the widespread relevance of personality genetics to a plethora of socially relevant behaviors and important life outcomes (Bleidorn et al., 2019; Roberts et al., 2007; Soto, 2019; Wright et al., 2023) across tests of genetic correlation, PGI prediction, and Mendelian Randomization (MR) applied to EUR GWAS data. Results for AFR GWAS data are reported in the **Supplementary Text** and **Supplementary Figures S30-S31**.

### Genetic correlations

Genetic correlations estimated using LDSC indicate widespread genetic sharing between personality traits and health-relevant daily behaviors. Conscientiousness in particular was linked to reduced substance use, greater sports participation, and greater preference for low-calorie foods, as well as the downstream consequences of these behaviors: fewer spells in the hospital, healthier aging, and lower BMI (**Figure 3**; **Supplementary Tables S31-S32**). Personality was also genetically correlated with fluctuations in accelerometer-measured behavior across the day, with openness to experience linked to increased night-time activity and conscientiousness to increased activity during the day (**Figure 2D**; Guerreiro et al., 2024).

**Figure 3.**
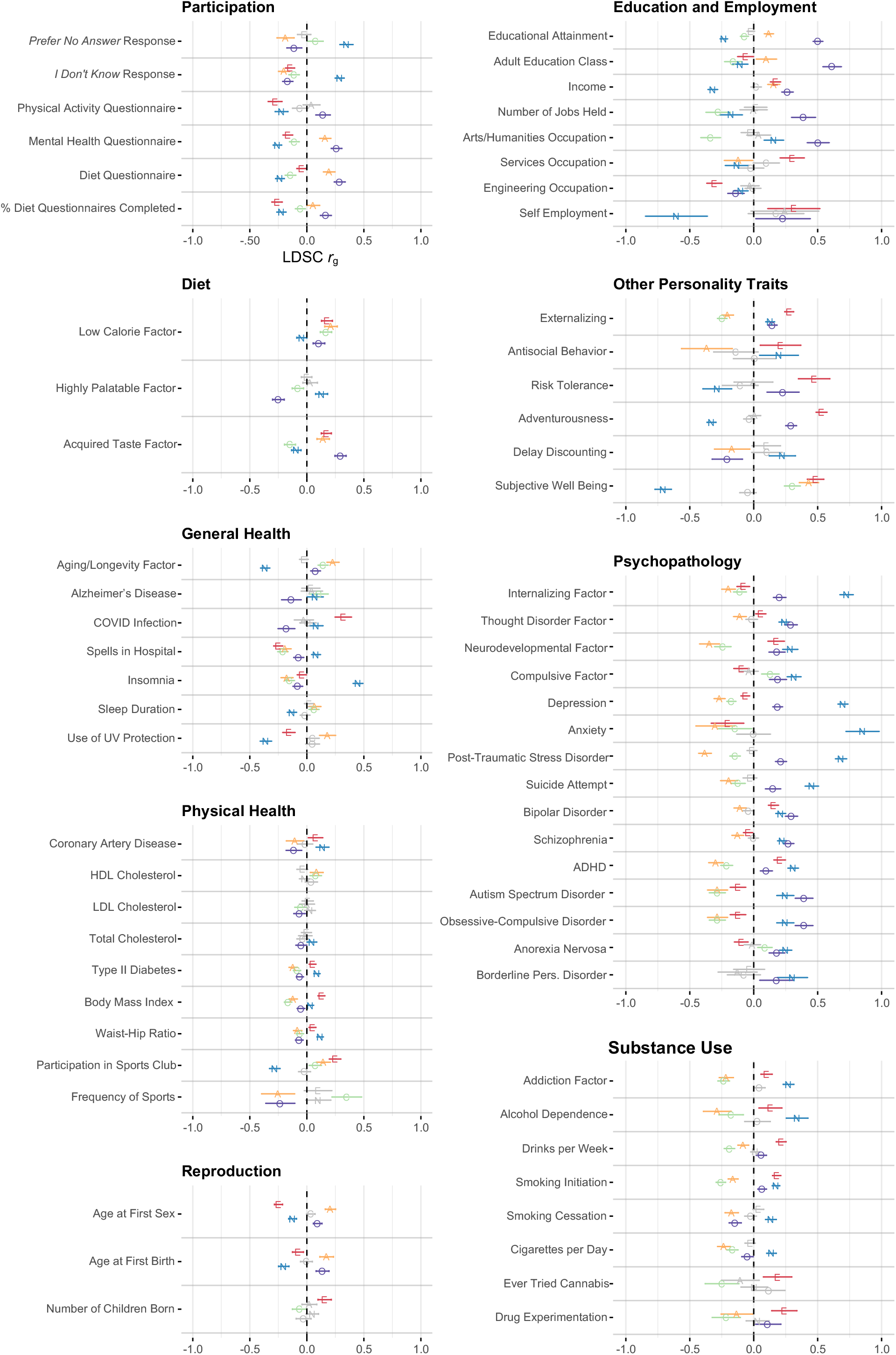
Genetic correlations between the Big Five personality traits and 67 behaviors and outcomes across 9 domains. E = Extraversion. A = Agreeableness. C = Conscientiousness. N = Neuroticism. O = Openness to experience. COVID = Coronavirus disease. HDL = High-density lipoprotein. LDL = Low-density lipoprotein. ADHD = Attention deficit hyperactivity disorder. Letters corresponding to personality traits represent correlation point estimates. Error bars represent 95% confidence intervals. Associations not significant at *p* < .01 are depicted in gray. Associations estimated using LDSC among EUR participants. Full description of each behavior and outcome are available in **Supplementary Tables S31 and S32.**

We found substantial genetic correlations between the Big Five and diagnosis liability for each of 10 psychiatric disorders and 4 transdiagnostic psychiatric disorder factors (**Figure 3**). Genetic profiles associated with neuroticism were associated with increased risk to all forms of psychopathology, particularly internalizing disorders, whereas genetic profiles associated with agreeableness displayed cross-cutting protective associations (Lo et al., 2017). We also identified broad patterns of genetic sharing between openness and psychopathology, which are notably discrepant from the small and inconsistent associations commonly reported in phenotypic research (Kotov et al., 2010; Widiger & Crego, 2019). Many other personality-psychopathology associations were specific and differentiated: Extraversion genetics were associated with lower risk for internalizing disorders but greater risk for neurodevelopmental disorders, and conscientiousness genetics were associated with lower risk for neurodevelopmental disorders but greater risk for compulsive disorders. These findings provide strong support for the close connection between personality traits and psychiatric disorders posited in psychiatric nosologies (Kotov et al., 2017).

Investigation of genetic correlations between personality traits and survey behavior in the UK Biobank indicated that personality plays a major role in research participation (**Figure 3**). In particular, genetic variants associated with openness to experience and agreeableness were associated with completing optional questionnaires, whereas genetic variants associated with neuroticism were associated with fewer questionnaire responses, more “*I don’t know*” and *“prefer no response”* answers, and higher rates of nonresponse after beginning a questionnaire. The genetics of personality may thus have a widespread, underacknowledged influence on sample composition and responses in published research that relies on survey data.

### PGI prediction of behaviors and outcomes

We examined PGI prediction of interviewer-rater qualities and delinquency, novel attributes that have not been examined in GWAS, in the Add Health cohort, a representative sample of young US adults (N = 5,110). We found that personality PGIs predicted how a young adult was perceived by their in-person interviewer during data collection (**Figure 2B; Supplementary Table S33**): Those with PGIs reflecting lower neuroticism and higher extraversion, agreeableness, and openness to experience were perceived to have a more attractive personality, and those with PGIs reflecting higher agreeableness and conscientiousness were perceived to be more well-groomed. In a subsample of at-risk Add Health participants, personality PGIs were associated with delinquent behavior and intergenerational contact with the prison system, (**Figure 2B; Supplementary Table S33**): high polygenic propensity for neuroticism and low polygenic propensity for agreeableness were associated with school suspension, running away from home, and ever pulling a knife or gun on another person.

Our socio-demographic analysis of UK Biobank participants (N = 419,261) linked personality PGIs to the characteristics of one’s residence in later adulthood, controlling for place of birth to reflect residential mobility. Focusing only on results that were significant in population-level analyses and sign-concordant in within-family analyses, genetic profiles indicative of higher openness to experience predicted moving into cosmopolitan professional areas and away from suburbia and challenged neighborhoods by middle-older adulthood (age 50+), whereas genetic profiles indicative of higher conscientiousness predicted moving away from cosmopolitan areas and into affluent suburban communities (**Figure 2E, Supplementary Figures S32-S33**). We present additional residential mobility analyses focused on urban/rural migration in the **Supplementary Text** and **Supplementary Figure S34**.

We also quantified personality similarity between spouses over recent generations. To do this, we estimated two PGIs from approximately balanced, non-overlapping halves of the GWAS discovery sample for each Big Five trait. We analyzed these halves with latent variable modelling to estimate genetic correlations among pairs of relatives that correct for PGI estimation error (**Supplementary Figure S36; Supplementary Table S34**). Spousal similarity is expected to induce genetic correlations on assorted traits among offspring that depart from their genetic relatedness, which can bias heritability estimates in standard population-based GWAS (Border et al., 2021; Horwitz et al., 2023). Inferred genetic correlations between parents, estimated using error-corrected PGI similarity meta-analyzed across spouses, sibling pairs, and cousin pairs from UK Biobank (N_dyads_ = 177,411) and deCODE (N_dyads_ = 133,358), indicated minimal historical assortative mating for each of the Big Five traits (Estimated parental *r*g = .01-.04; **Figure 2C**). Over past generations, birds of a feather have not flocked together: parents were only minimally more genetically similar in their personality traits, on average, than would be expected by chance, and far less genetically similar than has been estimated with respect to educational attainment (*r* = .37; Torvik et al., 2022). We present genetic correlations stratified by relationship and sample in **Supplementary Table S35** and phenotypic parental similarity estimates in the deCODE sample (N_pairs_ = 5,317) in **Supplementary Figure S35**.

### Mendelian randomization

To test causality and directionality in associations between personality traits and biobehavioral outcomes, we applied Mendelian Randomization (MR) tests, which leverage SNPs as instrumental variables in a natural experiment (Sanderson et al., 2022). Per MR best practices, we identified a subset of 13 outcomes most likely to comport with core assumptions of this method (**Supplementary Tables S36-S37**), we tested associations across three MR estimators (weighted median, MR-CAUSE, and weighted mode), and we considered associations significant only if they were directionally consistent across all three estimators with 95% confidence/credibility intervals that excluded zero across at least two of the three (see **Supplementary Text** and **Supplementary Figure 38** for complete information on MR methodology).

We found plausible causal effects for 33 exposure-outcome pairings (**Supplementary Table S38**). Twenty-four analyses indicated effects of personality on biobehavioral outcomes, providing novel evidence that personality traits modify physical health – for example, extraversion increased COVID infection risk, conscientiousness decreased BMI and reduced likelihood of smoking initiation and frequency of spells in hospital, and neuroticism decreased liability for a healthy aging/longevity factor. Furthermore, in line with theoretical perspectives that life experiences affect personality development (Roberts & Yoon, 2022), we found nine plausibly causal effects of biobehavioral outcomes on personality, encompassing broad multi-trait effects of educational attainment, increased BMI, and smoking initiation on personality. Reassuringly, we found no evidence for MR associations between personality and three negative controls (birthweight, number of sisters, and number of brothers) that would only be associated with one’s own personality through intergenerational confounding (residual gene-environment correlation). Our highly powered GWAS of each of the Big Five enabled these tests, providing valuable information on the potential causes and consequences of personality traits, which to date have been especially challenging to obtain using other methods (Grosz et al., 2020; Lucas, 2023). Nevertheless, future work should further triangulate on the MR-based causal inferences reported here with additional methods and data that permit strong causal inference (e.g. natural experiments; Bailey et al., 2024; Hammerton & Munafò, 2021).

### Evaluating Confounding Caused by Gene-Environment Correlation Polygenic prediction within dizygotic twin pairs

We used data from the Netherlands Twin Registry (N_families_= 2,956) and Twins Early Development Study (N_families_= 4,751) to predict personality traits within dizygotic twin pairs. These models predict personality from twin *differences* in PGIs, which we constructed from the primary EUR population GWAS meta-analysis, excluding the respective twin cohort used for PGI prediction. By holding family environment and parent genotype constant, and leveraging the randomness of intergenerational genetic transmission, these within-family comparisons substantially reduce genetic confounding associated with population stratification and assortative mating that may be present in population-level GWAS (Malanchini et al., 2024; Nivard et al., 2021; Selzam et al., 2019; Veller & Coop, 2024).

For all Big Five traits in both cohorts, within-twin-pair PGI differences robustly predicted the respective personality trait, with indistinguishable magnitude from population-level PGI prediction (Mean within-family β = 100.9% of population-level β, **Figure 4B**). This similarity between methods suggests no detectable confounding in population-level genetic effects on personality. This is in stark contrast with the notable predictive attenuation found in sibling PGI comparisons for other behavioral traits, such as educational attainment (mean within/population predictive strength ∼60%), cognitive ability (∼80%; Okbay et al., 2022; Tan et al., 2024), and externalizing psychopathology (∼75%; Karlsson Linnér, 2021).

**Figure 4.**
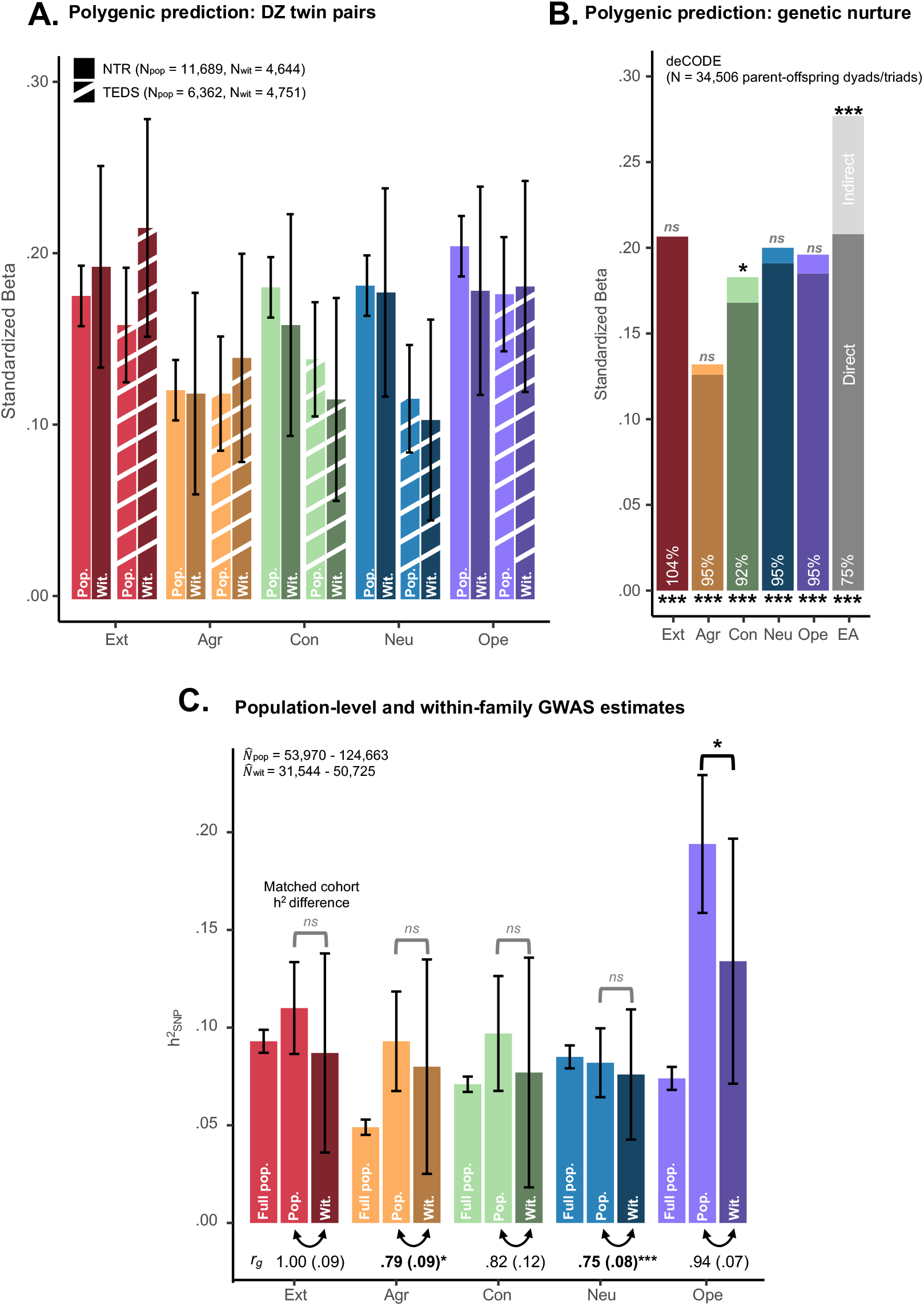
Evaluating environmental confounding in genetic associations with personality. In panel A, the magnitude of PGI prediction from the EUR population-level GWAS is presented for population-level and within-family analyses in two cohorts: Netherlands Twin Registry (NTR) and Twins Early Development Study (TEDS). In panel B, the magnitude of total PGI prediction from the EUR population-level GWAS among offspring in the deCODE cohort (with significance asterisks below the corresponding bar) is decomposed into directly transmitted and indirectly transmitted (non-inherited, with significance asterisks above the corresponding bar) genetic effects from parents (**Supplementary Table S39**). Comparison estimates for Educational Attainment (EA) come from a PGI constructed from the Okbay et al. (2022) GWAS of EA with deCODE and 23&Me cohorts held out. In panel C, h^2^_SNP_estimates estimated with LDSC from the within-family GWAS (“Wit.”) are presented alongside full EUR population-level GWAS (“Full pop.,” **Table 1**) and matched population-level (“Pop.”) GWAS in cohorts that contributed to within-family GWAS (**Supplementary Table S42**). *r*_g_ below each pair of traits indicates their genetic correlation estimated with LDSC, with deviation from *r*_g_ = 1 tested using a nested chi-square model in Genomic SEM. *p*-values above each pair of traits indicates results of a test for differences in heritability between population and within-family estimates. In all panels, error bars depict 95% confidence intervals and standard errors are in parentheses. *ns* = *p* ≥ .05. * = *p* < .05. ** = *p* < .01. *** = *p* < .001.

### Direct and indirect genetic effects on personality

We further tested for confounding using a complimentary method. We leveraged parent-offspring data from deCODE (N = 34,506) to decompose population-level PGI prediction into direct effects (genetic variants passed down from parent to offspring) and indirect effects (parental genetic variants not transmitted to the offspring). Across the Big Five, direct effects accounted for 96.2% of variance in in population-level PGI prediction, on average (**Figure 4; Supplementary Table S39**), indicating that genetic effects on personality operated nearly exclusive through direct transmission from parent to offspring. Only for conscientiousness were non-transmitted genetic effects significantly predictive of offspring personality (explaining 8.1% of the total predicted effect, *p* = .01). Non-transmitted effects did not differ in magnitude across mothers and fathers (**Supplementary Table S40**). In comparison, for educational attainment, only 75.1% of variance was explained by direct effects (**Supplementary Table S39**; Kong et al., 2018). After controlling for indirect effects, PGI prediction of extraversion by the extraversion PGI was similar in magnitude to PGI prediction of educational attainment by the educational attainment PGI. Thus, for personality traits, we find a negligible role of parental genetic nurture.

### Within-family genome-wide analyses

To conduct a maximally stringent test of genetic confounding, we supplemented our population-level GWAS with a within-family GWAS for each Big Five trait. Within-family GWAS accounts for effects of confounding at the level of individual SNPs (Friedman et al., 2021; Tan et al., 2024; Veller & Coop, 2024). To perform within-family GWAS, we assembled data across 12 contributing EUR cohorts (N = 31,544-50,725 across traits) that ascertained genetic data among family members using best-practices quality control (Tan et al., 2024), and we combined this with published within-family GWAS estimates of Neuroticism (Howe et al., 2022) (**Supplementary Table S41**; **Supplementary Figures S39-S43**).

For this within-family GWAS, LDSC-estimated h^2^_SNP_estimates ranged from 7.6% (SE = 1.6%) for agreeableness to 13.4% (3.2%) for openness to experience, evincing comparable magnitudes to those reported for the complete meta-analyses of population GWAS (**Figure 4**; **Supplementary Table S42**). To maximize comparability between within-family and population-level GWAS, we also estimated h^2^_SNP_using meta-analysis of population-level GWAS among the matched subset of 12 cohorts who contributed to the within-family GWAS (**Figure 4**). h^2^_SNP_comparisons across models, using *p* < .05 as a significance threshold to ensure identification of potential differences, indicated that for only openness to experience were matched population-level h^2^_SNP_estimates significantly greater than within-family h^2^_SNP_estimates (19.4% vs 13.4%, *p* = .02). This observed difference was driven largely by an inflated population-level h^2^_SNP_estimate for openness to experience relative to the full meta-analysis, and was specifically attributable to the Estonian Biobank sample (**Supplementary Table S41**). In contrast, previously-reported comparisons for other social and behavioral traits, such as educational attainment, depression, and household income, indicate LDSC h^2^_SNP_is attenuated by ≥ 50% in within-family data (Tan et al., 2024). LDSC-estimated genetic correlations between within-family and population-level GWAS summary data averaged *r*_g_ = .86 and were significantly less than 1.00 only for neuroticism (*r*_g_ = .75, SE = .08, *p* < .001) and agreeableness (*r*_g_ = .79, SE = .09, *p* = .03); these results indicate strong correspondence in genetic effect estimates between population-level and within-family analyses. The final piece of evidence for similarity between population-level and within-family genetic effects came from replication analyses of significant and suggestive effects (*p* < 1×10^−5^) from the full EUR population-level GWAS (excluding matched cohorts): on average, within-family estimates replicated at 95% the magnitude of population-level effect replication (see **Supplementary Table S42** for complete details).

## Discussion

In a major consortium effort assembling data from 46 cohorts covering 611K-1.14M EUR and AFR participants, we conducted highly-powered GWAS of each of the Big Five personality traits, producing dramatic gains in the number of discovered loci, validating powerful and robust polygenic indices, and comprehensively characterizing genetic architecture, confounding, assortative mating, and widespread genetic associations with socially relevant behaviors, health, and important life outcomes. These results overhaul the state of scientific knowledge on the genetic etiology of variation in human personality, establishing a rigorous basis for genetic inference and a fundamental role of personality genetics in the human condition.

Our results reveal that genetic associations with human personality are *generalizable* in several respects. Genetic effects on the Big Five are highly similar across geography, age groups, self-rated versus other-rated report, military service, and measurement instrument; furthermore, PGIs predicted personality traits with near-equivalent magnitude across the lifespan and in samples from different nations. Though the mechanisms by which genetic variance manifests in personality trait differences are sure to be numerous and complex, and may operate in part through dynamic developmental processes (Roberts & Jackson, 2008; Scarr & McCartney 1983), this observed similarity in genetic architecture and PGI prediction across groups suggests that widespread genetic inference is nonetheless possible. One key departure from this overall pattern pertains to agreeableness, which exhibited somewhat lower genetic correlations across geography and rater instrument, suggesting that mechanistic processes relating to this trait may be more varied and inference must be more contextualized.

We also demonstrated that genetic effects on human personality are relatively *unconfounded* by environmental factors that are shared across family members, including uncontrolled population stratification, assortative mating, and dynastic effects, each of which can inflate estimates of genetic effects (Veller & Coop, 2024). PGIs constructed using weights derived from population-level GWAS predicted each of the Big Five traits at indistinguishable strength in a standard population-level analysis and within-DZ twin pair analyses that account for these confounds. Parent-offspring analyses indicated that nearly all prediction was attributable to directly inherited genetic variants, rather than indirect effects of non-transmitted parental genotype. With the exception of openness to experience, estimates of h^2^_SNP_based on population GWAS were indistinguishable from those based on within-family GWAS that accounts for confounding. Effect size replication for top independent loci was very similar among population-level versus within-family hold-out samples. Finally, our comparisons of genetic correlations across relatives indicate that there has been minimal assortative mating on personality traits across recent generational time, extending decades of phenotypic research on romantic partner similarity (e.g. Horwitz et al., 2022; McCrae et al., 2008). Research in social science genomics has indicated that environmental sources of confounding play a major role in population-level GWAS of traits such as educational attainment, income, and cognitive performance (Howe et al., 2022; Okbay et al., 2022; Tan et al., 2024), introducing considerable challenge in obtaining estimates of direct genetic effects for these traits. In contrast, for personality, the lack of confounding indicates that population-level GWAS estimates overwhelmingly reflect direct genetic effects.

Functional genomic analyses and biological annotation indicated that despite their only modest genetic intercorrelations, genetic effects on each of the Big Five occur via overlapping molecular and cellular systems; for example, the prefrontal cortex was implicated in the genetics of each trait aside from agreeableness. Analyses also provide little support for neurobiological theories of the Big Five that posit trait-differentiated serotonergic and dopaminergic etiologies (Carver & Miller, 2006; DeYoung et al., 2021; Depue & Collins, 1999). Single cell gene expression data integrated with our GWAS suggests there is no straightforward reductive mapping from broad, multifaceted personality traits to specific neuron types. However, the availability of these high powered GWAS combined with ever-expanding access to temporal and spatial brain gene expression data will enable future developmental and system-specific analysis of neural personality etiology. Each of the Big Five demonstrated strong enrichment in protein truncating variant intolerant gene-sets specifically expressed in brain cells, suggesting that genetic variants with strong effects on personality are selected against (i.e. undergo negative selection), which is at odds with evolutionary theories that posit genetic variation in personality is solely maintained by inconsistent and varying selection pressures on personality that are in aggregate directionally neutral (balancing selection) (Penke & Jokela, 2016). Future research will be needed to further clarify the relative contributions of negative and balancing selection to variation in human personality traits.

Key limitations point to focal areas for future research. First, our within-family GWAS meta-analysis, while the largest of personality to date, was small compared to the population GWAS meta-analysis. Collection of additional within-family data, especially intergenerational family units (Davies et al., 2024) will provide further leverage to accurately and precisely identify loci indexing direct genetic effects. Fine mapping and experimental designs (e.g., Sanchez-Roige et al., 2023) will also be necessary to identify the causal variants within the identified loci. Second, we examined genetic correlations across only broad stratifying variables, and only among EUR and AFR individuals. More fine-grained sex- and age-stratified analyses and stratification by more expansive ancestral and cultural backgrounds will further refine inferences about the generalizability and differentiation of genetic effects on personality. Finally, we focused only on the very broad Big Five personality traits. Research with more granular personality trait data (e.g., of specific behavioral tendencies; Mõttus et al., 2017) will provide greater specificity of inference and permit more sophisticated multivariate analysis of the genetic patterning of personality trait structure.

Much of the scientific value of studying personality stems from its applicability to nearly all aspects of the lives that people lead. We extend this tenet by linking heritable variation in personality traits to core mental and physical health outcomes, labor market performance, and reproduction, with differentiated patterns that implicate certain traits (e.g. conscientiousness) as especially relevant to certain classes of outcomes (e.g. health behaviors). The genetics of personality also predict patterns of everyday behavior: how and when a person responds to surveys, where they move throughout their lives, and even the impression they make on others in a single interview session. Personality traits are highly stable over time (Bleidorn et al., 2022) and are resistant to simple experimental manipulation (Briley et al., 2018). Causal inference with respect to personality’s associations afforded by robust MR analyses is therefore especially valuable. MR results implicate personality traits as both causes and consequences of a range of health conditions. Given the pervasive relevance of personality to human experience, researchers across scientific fields will benefit from incorporating personality traits into models and theories when predicting and explaining biopsychosocial outcomes.

## Acknowledgements

T.S., M.G.N., and E.M.T-D. were supported by NIMH R01MH120219. E.M.T-D. was also supported by NIA R01AG073593. W.D.H., and C.X. were supported by a Career Development Award from the Medical Research Council (MRC) [MR/T030852/1] for the project titled “From genetic sequence to phenotypic consequence: genetic and environmental links between cognitive ability, socioeconomic position, and health.” T.K. was funded by the European Research Council (ERC Consolidator Grant awarded under the Horizon Europe framework, grant agreement 101087395. F.S. was supported by the Hector foundation II and by a 2023 NARSAD Young Investigator Grant (#31537) from the Brain & Behavior Research Foundation with support from the Families for Borderline Personality Disorder Research. R.C. was supported by the Jacobs Foundation grant no. 2023-1510-00 and the Research Council of Norway (grant number 325245). H.A. was supported by the research council of Norway (#274611, #324620). F.A.T. was supported by Research Council of Norway #26270. E.C.C. was supported by the research council of Norway (RCN; #274611) and the South-Eastern Norway Regional Health Authority (HSØ; #2021045). J.L. received financial support from the Strategic Research Council (SRC) established within the Academy of Finland (decision number: 352700). K.I. and U.Va. have been funded by Estonian Research Council’s personal research funding start-up grants PSG656 and PSG759. R.M. has been funded by Estonian Research Council’s team grant PRG2190. U.Võ. and T.E. have been funded by Estonian Research Council’s team grant PRG1291.

