## Supplementary Text and Figures for "Robust inference and widespread genetic correlates from a large-scale genetic association study of human personality"

*Ted Schwaba et al.*

**The PDF file includes:**

Methods and Supplementary Results, including Figs. S1 to S44
Contributing Cohort Information
References Not in Main Text

#### Table of Contents

##### Methods and Supplementary Results

**1. General information**
- 1.1. Cohorts
- 1.2. Measures
- 1.3. Software

**2. Population-level Association Meta-Analysis**
- 2.1. Cohort-specific Quality Control
- 2.2. GWAS Model
- 2.3. Quality Control Across Cohorts
- 2.4. Synthesizing Data Across Partially Overlapping Cohorts
- 2.5. Inverse-Variance Weighted Meta-Analysis
  - *Supplementary Figures S1-S10 (Manhattan plots of the Big Five in ancestry-stratified GWAS)*
- 2.6. X-Chromosome Analysis
- 2.7. Synthesis Across EUR and AFR participants
- 2.8. Replicating Genome-Wide Significant SNPs in Gupta et al. (2024).
- 2.9. Identifying Novel SNPs
- 2.10. Quantifying Overlap Among the Big Five

**3. Characterizing Common-Variant Heritability**
- 3.1. Estimating $h^2$SNP with Linkage Disequilibrium Score Regression
- 3.2. Genetic Correlations Across EUR and AFR Participants
- 3.3. Quantifying Heritability Across Cohorts with Random-Effects Meta-Analysis and Weighted Meta-Regression
  - *Supplementary Figure S11 (Meta-regressions of SNP heritability against scale reliability across cohorts)*
- 3.4. Characterizing Enrichment of Heritable Signal
  - *Supplementary Figures S12-S16 (Big Five gene set enrichment analyses)*
  - *Supplementary Figure S17 (Pairwise rank correlations of enrichment betas across the Big Five)*
- 3.5. Comparison of Genetic Architecture Across Grouping Variables
  - *Supplementary Figures S18-S22 (Genetic correlations across measurement instrument, geography, age, veteran status, and rater perspective)*
- 3.6. Genomic SEM Analyses Stratified by Measurement Instrument
  - *Supplementary Figures S23-S29 (Path diagrams, residual correlation matrices, GWAS results, and Qsnp analyses of instrument-stratified confirmatory factor analytic models)*

**4. Polygenic Prediction of Personality**
- 4.1. PGI Rationale and Cohorts
- 4.2. Method and Population-Level PGI model

**5. Associations with socially relevant behaviors and important life outcomes**
- 5.1. LDSC Correlations with External Phenotypes
  - *Supplementary Figure S30 (Genetic correlations between personality and health, psychopathology, and substance use among AFR participants)*
- 5.2. Polygenic Prediction of External Outcomes in Add Health

*Supplementary Figure S31 (Prediction of outcomes in the Add Health cohort from personality trait polygenic index among AFR participants)*
5.3. Residential Analyses
*Supplementary Figures S32-S33 (Residential preferences PGI heatmaps)*
5.3.1 Urban/Rural Migration Analyses
*Supplementary Figure S34 (PGI associations with moving or remaining in urban and rural areas in the UK Biobank cohort)*
5.4. Assortative Mating Analyses
*Supplementary Figure S35 (Phenotypic assortative mating correlations for the Big Five in the deCODE cohort)*
*Supplementary Figure S36 (Latent assortative mating model)*
5.5. Mendelian Randomization Analyses
*Supplementary Figure S37 (Causal diagram assumed by traditional Mendelian Randomization (MR) approaches)*
5.5.1 MR Method
5.5.2 Results and Causal Inference
**6. Evaluating Confounding Caused by Gene-Environment Correlation**
6.1. Within-family PGI Rationale and Model
6.2. Parent-offspring PGI Rationale and Model
6.3. Within-Family GWAS Cohorts and Methods
6.4. Within-family GWAS Quality Control
*Supplementary Figure S38 (Within-family Z-statistics before and after degrees of freedom correction among SNPs in the Minnesota Center for Twin and Family Research cohort)*
6.5. Inverse Variance-Weighted Meta-Analysis
6.6. Within-family GWAS Results
*Supplementary Figures S39-S43 (Big Five within-family GWAS Manhattan plots)*
6.7. Comparison of Population-Level and Within-Family Genetic Architectures
*Supplementary Figure S44 (Correlations among the Big Five in EUR population-level, EUR within-family, and AFR population participant data)*

**Contributing Cohort Information**

**References Not in Main Text**

### Methods and Supplementary Results

#### 1. General Information

1. 1. Cohorts

We incorporated data from 46 cohorts participating in the Revived Genomics of Personality Consortium (ReGPC). Each cohort provided data on at least one Big Five trait administered to a sample of genotyped participants; each analysis thus incorporates a subset of these 46 cohorts. Given the substantial growth in contributing sample sizes and advances in GWAS best practices since the previous Genomics of Personality Consortium (de Moor et al., 2012, 2015; van den Berg et al., 2016), data from those consortium cohorts were included if they were re-analyzed specifically for this study. Complete cohort-specific information on ascertainment, genotyping, and quality control is provided narratively in the **Contributing Cohorts** section at the end of this document, as well as in **Supplementary Tables S1 and S2.**

1.2 Measures

In each cohort, one or more Big Five personality traits were assessed via multi-item questionnaire scales (see **Supplementary Tables S1 and S2** for complete information). Scales qualified for study inclusion under four criteria (see SOP Analysis Plan document at https://osf.io/hgnsm/?view_only=f98d2f87f8b8472d9d49e42455701d23). First, we included data from validated questionnaires specifically designed to measure the Big Five, such as the Big Five Inventory (John et al., 1991) or the NEO Personality Inventory-Revised (Costa & McCrae, 2008). This comprised most of our data (k = 39 cohorts). We also included the extraversion and neuroticism scales of the Eysenck Personality Questionnaire (EPQ; Eysenck & Eysenck, 1984; k = 4 cohorts); phenotypic research and our genetic comparisons in the **Main Text** and section 3.6 (Genomic SEM Analyses Stratified by Measurement Instrument) confirm that these scales are highly congruent with extraversion and neuroticism as instantiated in the Big Five. We included the extraversion, conscientiousness, and openness scales of the HEXACO inventory (k = 1); these three traits are theoretically congruent with the Big Five (Lee & Ashton, 2004). Finally, we included data from Big Five trait scales created specifically for these analyses using factor analysis from item subsets of a broader personality questionnaire (k = 2). We excluded scales that measure personality traits outside the Big Five (e.g., psychoticism), measured personality with a single item, or measured narrower personality facets of the Big Five (e.g., sociability).

Typically (in k = 45/46 studies) participants responded to the questionnaire by self-reporting the extent to which various adjectives (e.g., "*Talkative"*) or phrases (e.g., *"Talks a lot"*) apply to them, using either binary logic ("*Yes*" or *"No*") or a multiple-point Likert scale (*"Disagree a lot" – Agree a lot*"). Most scales contained negatively-worded items assessing a trait's low pole (e.g., *"Quiet"*) that were reverse-scored. For each trait scale, we averaged responses across items to form a multi-item continuous composite score. In the ALSPAC cohort, participants' personality traits were reported on by their mother, and in supplementary EBB cohort data not incorporated in the discovery GWAS, participants' personality traits were reported on by a nominated close other. Comparisons reported in the **Main Text** confirm that it is appropriate to synthesize scores across these rater perspectives.

In cohorts that measured personality traits at multiple time points, we aggregated this longitudinal information into a maximally trait-like score by averaging a participant's scores across these measurement occasions and conducting analyses on this aggregated phenotype. To facilitate synthesis across cohorts, scores were standardized to a mean of 0 and a standard deviation of 1 prior to analyses.

1.3. Software

Analyses were conducted using the Howe and colleagues (2022) within-sibship analytic pipeline, the programs fastGWA (Jiang et al., 20199), PLINK (Purcell et al., 2007), PLINK2 (Chang et al., 2015), SBayesR (Lloyd-Jones et al., 2019), SNiPAR (Young et al., 2022), SLD4M (O'Connor et al., 2019), and R version 4.3.1 (R Core Team, 2023) using R data-management packages dplyr (Wickham et al., 2019), data.table (Dowle et al., 2019), and psych (Revelle, 2017), R visualization packages corrplot (Wei et al., 2017), ggplot2 (Wickham et al., 2019), and qqman (Turner, 2014), and R genomic analysis packages EasyQC (Winkler et al., 2014), GenomicSEM (Grotzinger et al., 2019), and ieugwasr and TwoSampleMR (Hemani et al., 2018).

#### 2. Population-level Association Meta-Analysis

2.1 Cohort-Specific Quality Control

For each Big Five trait, we conducted population-level genome-wide association analyses among participants with European-like (EUR) and African-like (AFR) genomes across the 22 autosomes and the X chromosome.

As a preliminary step, participants were stratified by recent genetic ancestry to minimize confounding due to population structure and allelic frequency differences. The strategy for genetic ancestry identification varies by cohort (see **Contributing Cohorts** below for specifics). Typically, analysts clustered participants by plotting multivariate vectors of local genetic variation, identifying maximally homogenous subgroups of participants, and then labeling these clusters with respect to similarity to 1000 Genomes Project superpopulations (1000 Genomes Project Consortium, 2015).

Analysts then implemented pre-imputation quality control according to the following specifications: genotypes were aligned to build 37/hg19 on the forward strand, excluding SNPs with Minor Allele Frequency (MAF) < 1%, Call rate <95%, those that deviated from Hardy-Weinberg Equilibrium ($p < 1x10^{-6}$), and those with poor clustering on visual inspection of intensity plots. Participants were excluded if they had low overall call rates (< 95%), excess autosomal heterozygosity or homozygosity, were duplicated samples, had gender inconsistent with sex (which is often indicative of mislabeled demographic data; Turner et al., 2011), or had chromosomal abnormalities.

Imputation of estimated genetic variants from genotyped data was done per-cohort; SNPs were removed with poor imputation quality (INFO < .40); other analyses used more stringent INFO thresholds when noted.

Across cohorts, 2.5 to 8 million SNPs passed these QC filters for each trait (see **Supplementary Tables S1 and S2** for information on each cohort).

2.2 GWAS Model

For each trait in each cohort, stratified by EUR and AFR participants, we regressed personality trait score, *y,* on each available autosomal biallelic SNP using an additive genetic model (SNP = 0, 1, or 2 effect alleles), controlling for chromosomal sex, age and age$^2$ (to account for nonlinear developmental trajectories of personality traits across the lifespan; Bleidorn et al., 2022), 20-year birth cohort (in cohorts with birth year range > 20 years), 10-20 genetic principal components (PCs) for ancestry, and, when appropriate, other covariates such as genotyping batch or questionnaire language (see **Supplementary Tables S1 and S2** for specific controls applied to each cohort), according to the following equation:

$$y = b_0 + b_1 SNP + \sum b_c x_c + e \, , \qquad \text{(Eq. 1)}$$

where $b_0$ is a regression intercept, $b_1$ is the linear effect of the allele account for the SNP, $b_c$ is the effect of covariate $x_c$ on the personality trait for covariates 1 through C, and *e* is a residual. In some cohorts, age and age$^2$ were highly collinear; in these cases, age$^2$ was excluded from the set of covariates.

In cohorts of unrelated participants, the above GWAS model was estimated using ordinary least squares (OLS). In cohorts that included data from related participants, analysts either a) used genetic information to estimate genetic relatedness and exclude one member of each first- or second- degree relative pair, or b) employed a method, such as a mixed model (using the software such as GCTA, Yang et al., 2011, or fastGWA, Jiang et al., 2019), to correct for the inclusion of related individuals.

2.3 Quality Control Across Cohorts

Following per-cohort analysis, a second uniform round of quality control was applied to data. SNPs were further restricted according to the following criteria: inclusion in the 1000 Genomes European/African Superpopulation SNP list, and SNP is biallelic. Effect (A1) and Noneffect (A2) alleles were aligned across cohorts using EasyQC (Winkler et al., 2014).

Cohort analysts were instructed to code participant scores such that high scores are indicative of high extraversion, high agreeableness, high conscientiousness, high neuroticism, and high openness to experience. To ensure that effects were properly coded in this direction, each trait in each cohort was entered into bivariate LDSC (Bulik-Sullivan et al., 2015) and genetically correlated with the large MoBa cohort. Effects were flipped ($b_1$ multiplied by -1) in traits with substantial negative genetic covariance with MoBa.

Cohort analysts were instructed to standardize phenotypes prior to GWAS. The squared standard error of a GWAS regression coefficient (*b*) is a function of the coefficient, sample size (*n*), and variances of the predictor ($\sigma_{SNP}^2$) and GWAS phenotype ($\sigma_y^2$), as follows:

$$(SE_b)^2 \approx \frac{\sigma_y^2 - \sigma_{SNP}^2 (b^2)}{\sigma_{SNP}^2 n}. \qquad \text{(Eq. 2a)}$$

Noting that variance of the SNP under Hardy-Weinberg Equilibrium is $2MAF \times (1 - MAF)$, we estimated the standard deviation of the GWAS phenotype ($\widehat{SD}_y$)to ensure that effects were indeed standardized in contributing cohorts for each SNP using its minor allele frequency (MAF), effect size, standard error of effect, and sample N.

$$\widehat{SD}_y = \sqrt{n \times SE_b^2 (2MAF \times (1 - MAF)) + b^2 (2\,MAF \times (1 - MAF))}. \qquad \text{(Eq. 2b)}$$

In cohorts where the median estimated variance (${\widehat{SD}_y}^2$) deviated substantially from 1, effects were re-standardized by dividing all betas and standard errors by the median estimated standard deviation.

2.4 Synthesizing Data Across Partially Overlapping Cohorts

Some groups contributed separate analyses on potentially-overlapping cohorts of participants (e.g., a participant may have contributed data for the same trait, potentially measured by a different instrument, to multiple cohorts). We empirically tested for overlap between these cohorts by correlating GWAS Z-statistics ($b$ / SE) across pairs of potentially overlapping cohorts, and we compared this value to the expected sampling correlation of the estimates, based on the rationale that a single cohort of participants has low power to detect true genetic effects, such that its GWAS estimates predominantly represent sampling variability, as opposed to true signal (as evidenced by the mean association chi-square for cohort-level GWAS rarely departing appreciably from the expectation under the null of 1.0). The expected sampling correlation between GWAS estimates $b_1$ and $b_2$ for cohorts 1 and 2 ($\sigma_{b1,b2}$) is:

$$\hat{\sigma}_{b1,b2} = r_{pheno} \frac{n_s}{\sqrt{n_1 n_2}}, \qquad \text{(Eq. 3)}$$

where $r_{pheno}$ is the phenotypic correlation among the overlapping participants, $n_s$ is the number of overlapping participants in the two cohorts, and $n_1$ and $n_2$ are the sample sizes in cohorts 1 and 2 respectively.

For three pairs of cohorts (two each collected as part of Generation Scotland, Lifelines, and FinnTwin), Z-statistic correlations approximately matched the expected sampling covariance and were substantially positive, indicating non-negligible participant overlap (see **Supplementary Table S3**). In these three cohort pairs, we computed the expected sampling covariance matrix (V) of each SNP by rescaling the sampling correlation matrix (R) by the respective standard errors of the GWAS coefficients (contained on the diagonals of a diagonal matrix D) for that SNP in the two cohorts as follows:

$$V = DRD. \qquad \text{(Eq 4a)}$$

We then meta-analyzed the estimates, taking into account the dependency of their estimation errors, using generalized least squares (GLS; Aitken, 1936) as follows:

$$b_{meta} = (X'V^{-1}X)^{-1}X'V^{-1}b\,, \qquad \text{(Eq. 4b)}$$

and

$$SE_{b_{meta}}^2 = (X'V^{-1}X)^{-1}\,, \qquad \text{(Eq. 4c)}$$

where $b$ is the vector of GWAS coefficients ($b_1$ and $b_2$ for cohorts 1 and 2) for a given SNP in each of the partially overlapping cohorts, $X$ is a vector of 1s (2 elements for two cohorts), $b_{meta}$ is the meta-analytic beta across the overlapping cohorts, and $SE_{b_{meta}}^2$ is its standard error. This synthesis consolidated the 46 contributing EUR samples into 43 total independent samples in addition to 10 independent AFR samples.

2.5 Inverse-Variance Weighted Meta-Analysis

To obtain the meta-analytic GWAS estimates $b_{meta}$, Standard Error ($SE_{b_{meta}}$), and Z statistic ( $Z_{b_{meta}}$) for each SNP for each personality trait across K independent cohorts, we implemented an inverse variance weighted meta-analysis in R, according to the following set of equations:

$$b_{meta} = \frac{\sum \frac{1}{SE_{b_k}^2} b_k}{\sum \frac{1}{SE_{b_k}^2}} , \qquad \text{(Eq. 5a)}$$

$$SE_{b_{meta}} = \frac{1}{\sqrt{\sum \frac{1}{SE_{b_k}^2}}}, \qquad \text{(Eq. 5b)}$$

and

$$Z_{b_{meta}} = \frac{b_{meta}}{SE_{b_{meta}}} . \qquad \text{(Eq. 5c)}$$

We also estimated imputation quality for non-genotyped SNPs (INFO), effect allele frequency (EAF) and MAF in the full meta-analytic sample using sample-sized weighted meta-analysis, according to the following set of equations:

$$INFO_{meta} = \frac{\sum n_k INFO_k}{\sum n_k} , \qquad \text{(Eq. 6a)}$$

$$MAF_{meta} = \frac{\sum n_k MAF_k}{\sum n_k} , \qquad \text{(Eq. 6b)}$$

and

$$EAF_{meta} = \frac{\sum n_k EAF_k}{\sum n_k} . \qquad \text{(Eq. 6c)}$$

We then compared each SNP's meta-analytic EAF with its EAF in the 1000 Genomes Project phase 3v5, pruning SNPs with a discrepancy greater than 10%, as suggested by Lloyd-Jones and colleagues (2019).

Because some cohorts estimated genetic associations using data from related participants and employed methods (such as mixed models) that correct for relatedness, reported sample size for each analysis may not correspond directly with the effective sample size. If the observed sample size is used in LDSC, and the effective sample size is lower than the observed sample size, heritability estimates will be deflated (Grotzinger et al., 2023). To account for this, we re-estimated sample size ($\widehat{N}$) for each SNP using the standard error of effect, and its expected variance under Hardy-Weinberg Equilibrium, according to the following equation:

$$\widehat{N} = \frac{1}{2 \times MAF \times (1 - MAF) \times SE_{b_{meta}}^2} , \qquad \text{(Eq. 7)}$$

where the numerator is the variance of the phenotype, which has been standardized to 1. Estimated sample size was capped at the summed reported sample size for each SNP, and SNPs with $\widehat{N} \leq 5{,}000$ were pruned from the final set of meta-analytic summary statistics. This resulted in a final set of between 9.3 and 10.7 million SNPs per trait per ancestry (see **Supplementary Table S4**).

We submitted these summary statistics to FUMA (Watanabe et al., 2017) to quantify the number of genome-wide significant lead SNPs. We thus define a lead SNP in line with FUMA as a common genetic variant (MAF ≥ .01) that is significant at $p < 5x10^{-8}$ (i.e., .05, Bonferroni-corrected for 1 million independent tests) and independent of other lead SNPs (linkage disequilibrium $R^2 < .10$) nearby on the genome (within a 250kb window). We present Manhattan plots of associations stratified by ancestry in **Supplementary Figures S1-S10**, below.

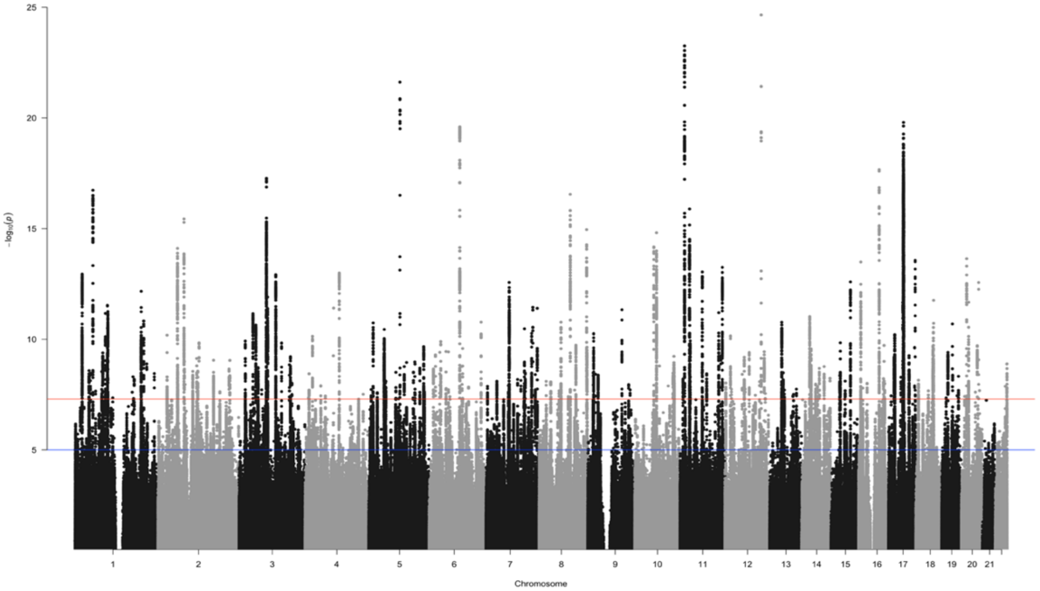


**Supplementary Figure S1. Manhattan plot of extraversion in EUR participants (N = 620,193).** Note: Blue line indicates $p = 5x10^{-6}$ and red line indicates $p = 5x10^{-8}$. *P*-values smaller than this threshold are considered significant.

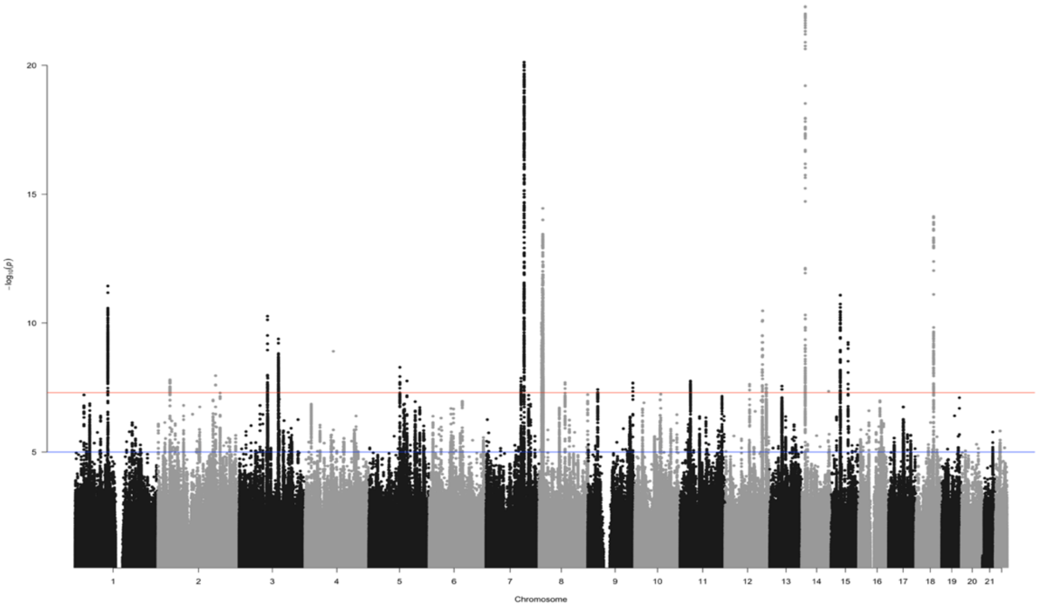


**Supplementary Figure S2. Manhattan plot of agreeableness in EUR participants (N = 567,556)** Note: Blue line indicates $p = 5x10^{-6}$ and red line indicates $p = 5x10^{-8}$. *P*-values smaller than this threshold are considered significant.

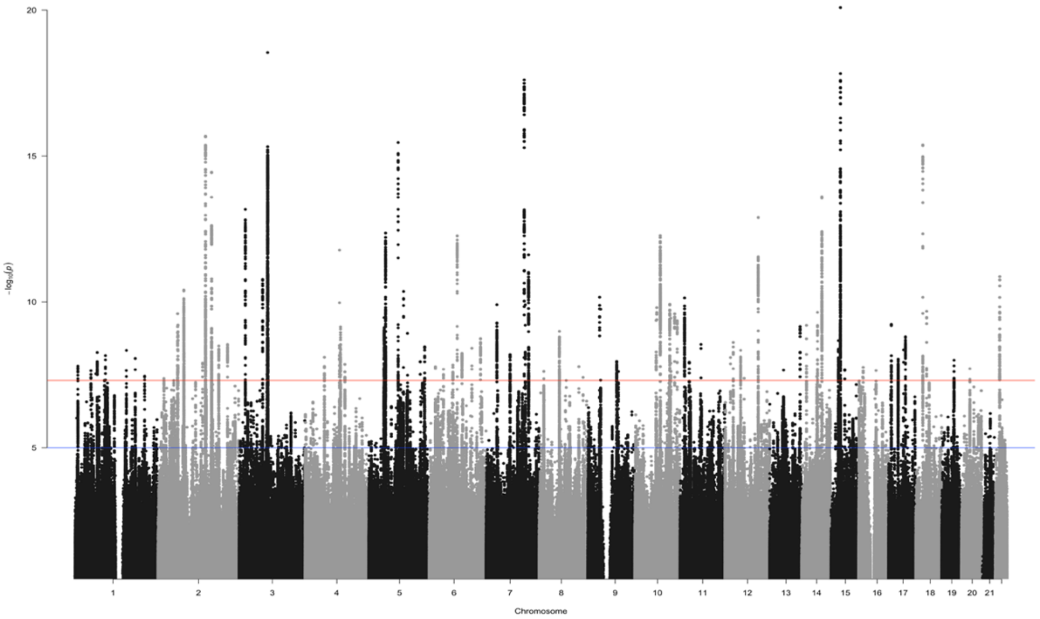


**Supplementary Figure S3. Manhattan plot of conscientiousness in EUR participants (N = 598,047)** Note: Blue line indicates $p = 5x10^{-6}$ and red line indicates $p = 5x10^{-8}$. *P*-values smaller than this threshold are considered significant.

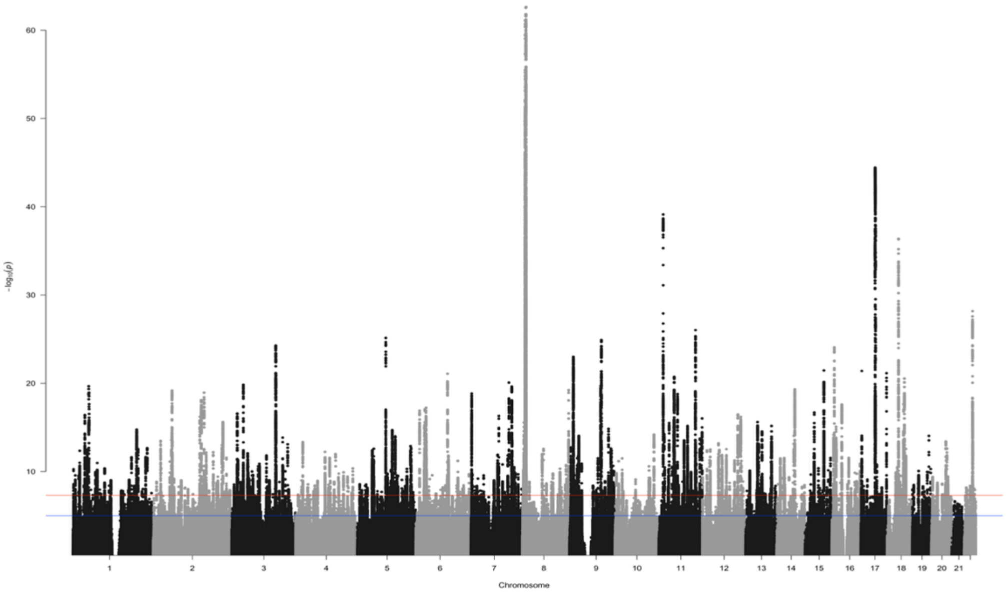


**Supplementary Figure S4. Manhattan plot of neuroticism in EUR participants (N = 1,088,604)** Note: Blue line indicates $p$ = 5x$10^{-6}$ and red line indicates $p$ = 5x$10^{-8}$. *P*-values smaller than this threshold are considered significant.

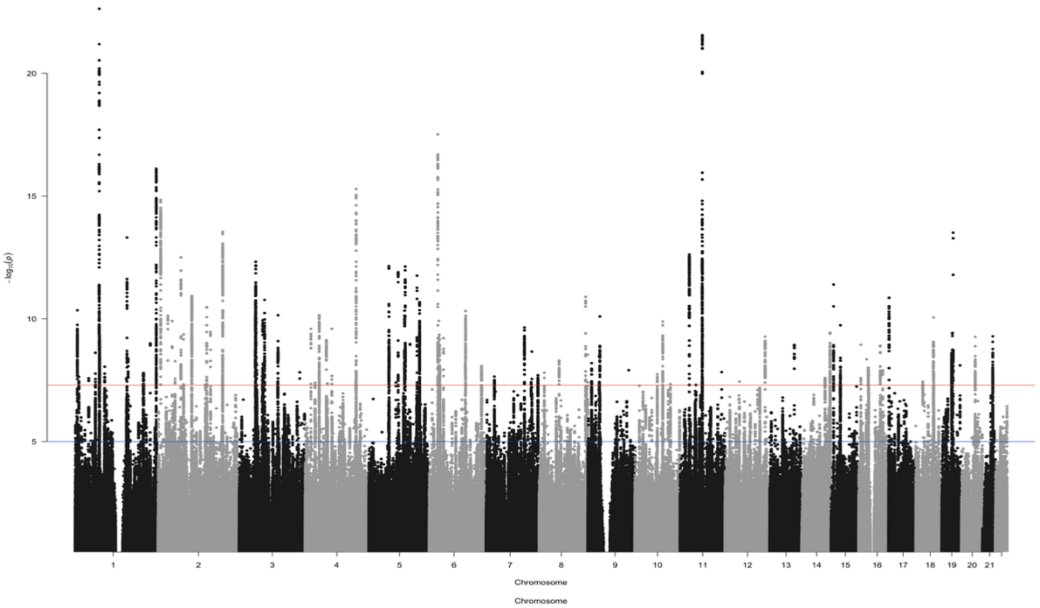


**Supplementary Figure S5. Manhattan plot of openness to experience in EUR participants (N = 568,489)** Note: Blue line indicates $p$ = 5x$10^{-6}$ and red line indicates $p$ = 5x$10^{-8}$. *P*-values smaller than this threshold are considered significant.

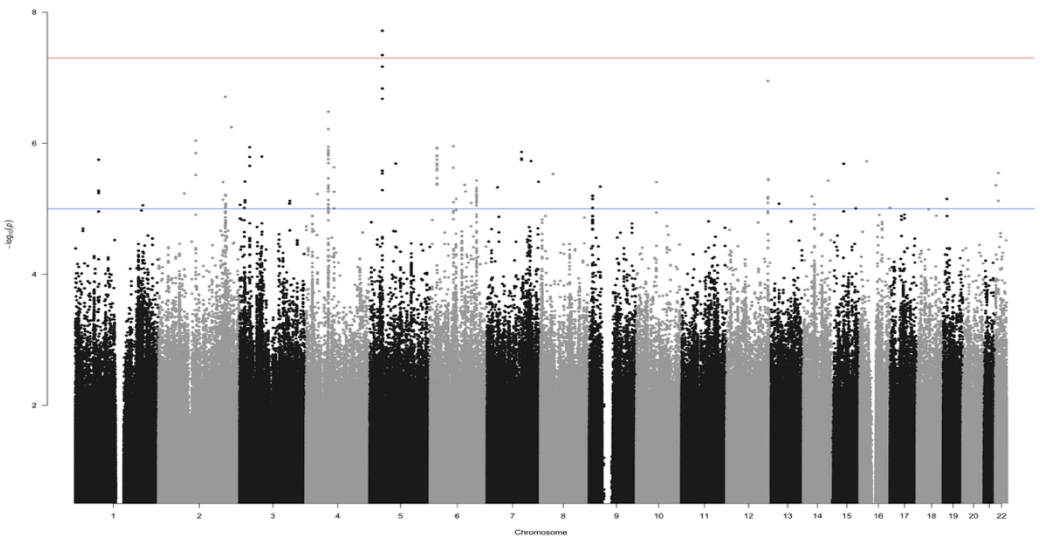


**Supplementary Figure S6. Manhattan plot of extraversion in AFR participants (N = 43,201)** Note: Blue line indicates $p = 5x10^{-6}$ and red line indicates $p = 5x10^{-8}$. *P*-values smaller than this threshold are considered significant.

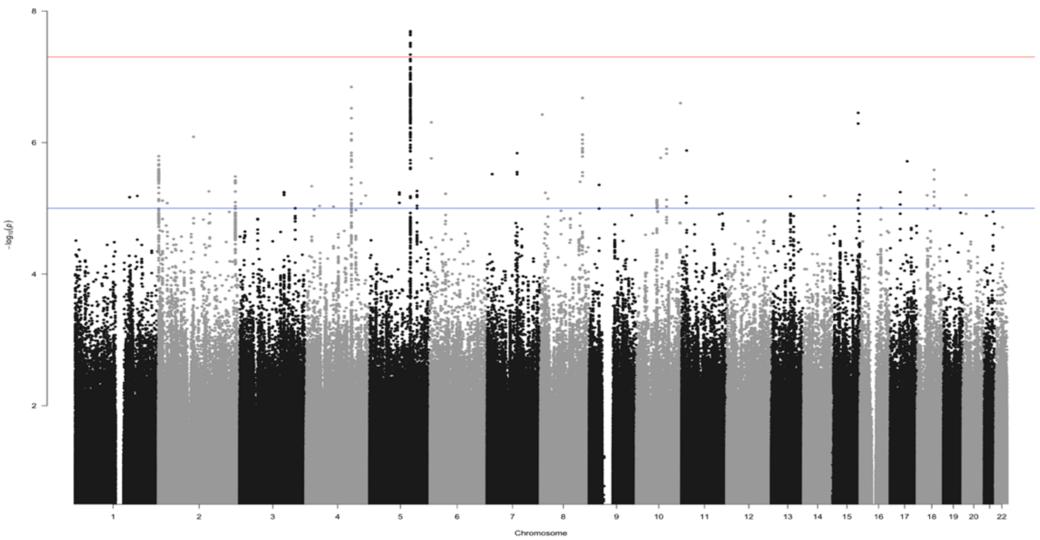


**Supplementary Figure S7. Manhattan plot of agreeableness in AFR participants (N = 43,482)** Note: Blue line indicates $p = 5x10^{-6}$ and red line indicates $p = 5x10^{-8}$. *P*-values smaller than this threshold are considered significant.

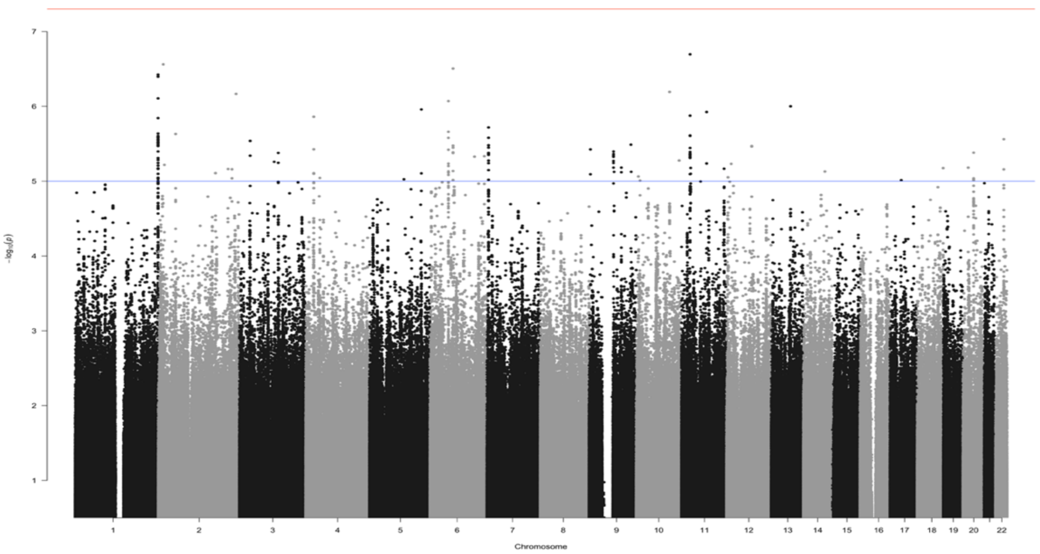


**Supplementary Figure S8. Manhattan plot of conscientiousness in AFR participants (N = 43,120)** Note: Blue line indicates $p$ = 5x10$^{-6}$ and red line indicates $p$ = 5x10$^{-8}$. $P$-values smaller than this threshold are considered significant.

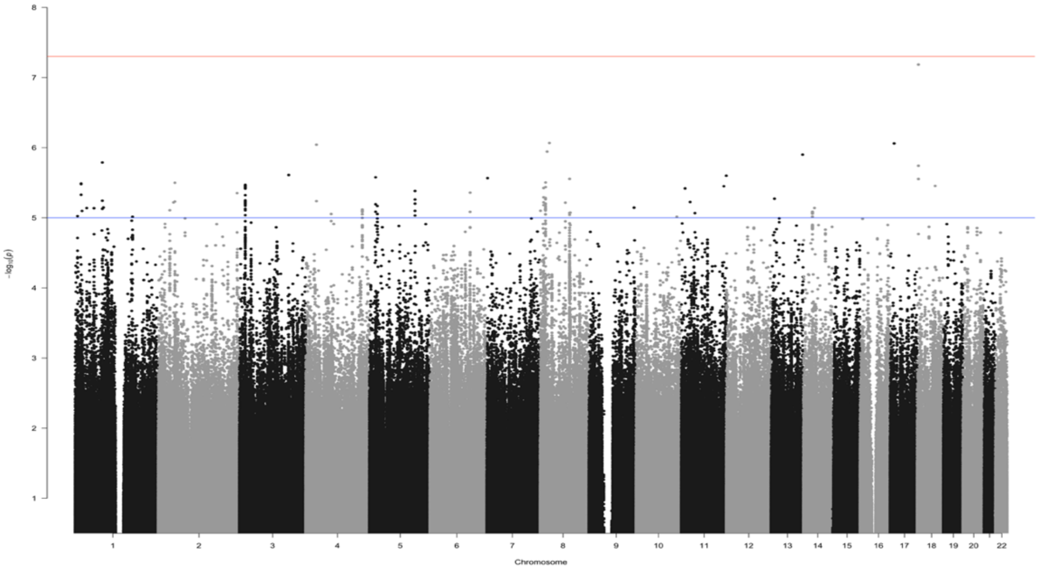


**Supplementary Figure S9. Manhattan plot of neuroticism in AFR participants (N = 48,110)** Note: Blue line indicates $p$ = 5x10$^{-6}$ and red line indicates $p$ = 5x10$^{-8}$. $P$-values smaller than this threshold are considered significant.

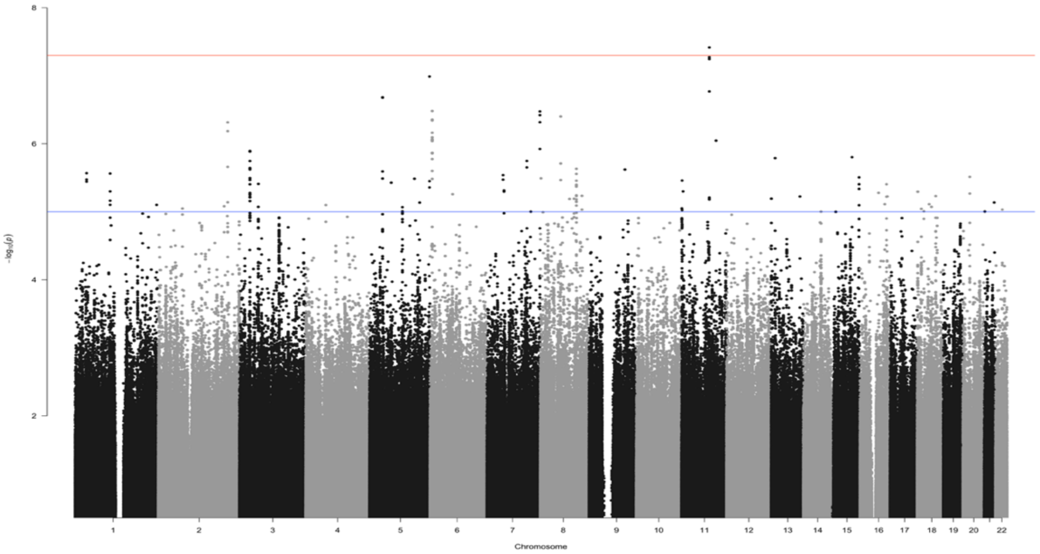


**Supplementary Figure S10. Manhattan plot of openness to experience in AFR participants (N = 43,498)** Note: Blue line indicates $p = 5x10^{-6}$ and red line indicates $p = 5x10^{-8}$. *P*-values smaller than this threshold are considered significant.

In EUR participants, on chromosomes 1-22, we identified 234 lead SNPs for extraversion (minimum *p*-value $4.11\times10^{-29}$), 32 lead SNPs for agreeableness (minimum *p*-value $5.21\times10^{-23}$), 104 lead SNPs for conscientiousness (minimum *p*-value $8.20\times10^{-21}$), 667 lead SNPs for neuroticism (minimum *p*-value $2.33\times10^{-63}$), and 113 lead SNPs for openness to experience (minimum *p*-value $2.32\times10^{-23}$). In AFR participants, we identified 1 lead SNP each for extraversion ($p = 1.92\times10^{-8}$), agreeableness ($p = 2.03\times10^{-8}$), and openness to experience ($p = 3.84\times10^{-8}$). We provide summary information on these GWAS results in **Supplementary Table S4**.

2.6 X-Chromosome Analysis

We estimated associations between X-chromosome SNPs and personality traits, in the six cohorts that contributed data incorporating this chromosome. Each of these six cohorts ascertained EUR participants. Quality control for these data followed the same procedure as described above. When possible, we conducted quality control on sex-stratified data. We excluded the two pseudo-autosomal regions on the X-chromosome from analysis, as few cohorts provided data from this genomic region.

In females, effects were coded as diploid (i.e., participants could have 0, 1, or 2 effect alleles, as with autosomal analyses above). In males, effects were coded as homozygous diploid, with dosage compensation (i.e., participants could have 0 or 2 effect alleles). In male samples without dosage compensation coding, betas and standard errors were divided by 2 to rescale effects equivalently to models with dosage compensation.

We then meta-analyzed X-chromosome associations together across male-stratified, female-stratified, and sex-combined samples following equation 5 above. We again re-estimated meta-analytic sample size for each SNP following equation 7. For male data, SNP variance in the denominator of this equation was instead estimated using $4MAF \times (1 - MAF)$ instead of $2MAF \times (1 - MAF)$ to account for dosage compensation.

Manhattan plots of X-chromosome results are presented in **Figure 1** of the main text. We identified one additional locus containing a lead SNP on the X chromosome, for agreeableness ($p = 9.83\times10^{-9}$; **Supplementary Table S6**). We note that as only six cohorts provided X-chromosome data, these analyses were comparatively underpowered relative to autosomal analyses.

2.7 Synthesis Across EUR and AFR Participants

To maximize GWAS discovery power, we combined results across EUR and AFR participants and identified additional independent lead SNPs in this transancestral sample. Specifically, we applied inverse-variance-weighted meta-analysis (equation 5) to synthesize the two sets of summary data for each trait: the k = 10 cohorts comprising analyses among AFR and EUR participants and the additional k = 36 cohorts comprising analyses among EUR participants. For this meta-analysis, we aligned SNPs to the 1000 genomes 3v5 EUR reference panel, as most participants (~95%) were of EUR ancestry, and stronger linkage disequilibrium among EUR compared to AFR superpopulations (Reich et al., 2001) means that aligning results to the EUR reference panel likely leads to conservative estimation of the number of lead SNPs. We present Manhattan plots of GWAS associations in this transancestry sample in **Figure 1** of the main text.

We then identified novel lead SNPs in this analysis. First, we submitted results to FUMA (Watanabe et al., 2017), which identified 111 potential novel lead SNPs aligned to the 1000 genomes 3v5 EUR reference panel. For each of these potential lead SNPs, we estimated a sample-size weighted ancestry-combined LD matrix (across 1000g 3v5 EUR and 1000g 3v5 AFR, including only SNPs that were included in both reference panels) that included the potential lead SNP and all other lead SNPs on that chromosome identified in the transancestry, AFR, and EUR stratified analyses. If this SNP was correlated with all other lead SNPs at $R^2 < .10$, we considered it a novel independent lead SNP beyond those identified in the ancestry-stratified analyses.

Of the 111 potentially novel lead SNPs identified in the transancestry GWAS, 103 remained LD-independent of all other lead SNPs in the synthesized transancestry LD matrix. We report the maximum LD of each of these SNPs in **Supplementary Table S10**.

To further characterize the independent strength of association of lead SNPs identified in this ancestry-combined sample, we conducted a conditional and joint analysis using the multiSNP() function in the GenomicSEM R package (Grotzinger et al., 2019). Specifically, we predicted each new lead SNP identified in the ancestry-combined sample jointly from each of the lead SNPs on that chromosome identified in any genome-wide analysis (EUR, AFR, and transancestry), using the transancestry LD matrix described above. Results of these analyses, presented in **Supplementary Table S11**, describe the extent to which each lead SNP is associated with each trait *over and above* other lead SNPs on that chromosome. As expected, many SNPs with $p$-values just below the genome-wide significance threshold were no longer significant after controlling for other lead SNPs, even if those other lead SNPs are nearly LD-independent.

In total, synthesizing across all GWAS results, we identified 1,257 lead SNPs. Breaking down by trait, we identified 258 lead SNPs for extraversion: 234 in EUR analyses, 1 in AFR analyses, and 23 additional lead SNPs in transancestry analyses, where 20 EUR lead SNPs were no longer significant. We identified 39 lead SNPs for agreeableness: 33 in EUR analyses (of which one was on the X chromosome), 1 in AFR analyses, and 5 additional lead SNPs in transancestry analyses, where 3 EUR lead SNPs were no longer significant. We identified 131 lead SNPs for conscientiousness: 104 in EUR analyses and 27 additional lead SNPs in transancestry analyses, where 14 EUR lead SNPs were no longer significant. We identified 703 lead SNPs for neuroticism: 667 in EUR analyses and 36 additional lead SNPs in transancestry analyses, where 27 EUR lead SNPs were no longer significant. Of the 703 neuroticism lead SNPs, 684 were independent of all other neuroticism lead SNPs regardless of genomic distance (i.e. they were not in long-range LD with any other neuroticism lead SNPs). However, 19 lead SNPs for neuroticism were in non-overlapping genomic loci (mean distance = 2,941,471b) but nonetheless were in LD ($r^2$ > .10), for example due to their location on inversion regions. This pattern has been found in previous neuroticism GWAS (Okbay et al., 2016). Finally, we identified 126 lead SNPs for openness to experience: 113 in EUR analyses, 1 in AFR analyses, and 12 additional lead SNPs in transancestry analyses, where 9 EUR lead SNPs were no longer significant. We note that this relative frequency of lead SNPs that were significant in EUR analyses but not transancestry analyses is similar to that in Gupta and colleagues (2024). Each of the lead SNPs across these analyses are presented in **Supplementary Tables S5-S9.**

2.8 Replicating Genome-Wide Significant SNPs in Gupta et al. (2024).

We quantified the similarity of these results to those from Gupta and colleagues (2024), the most recent GWAS of the Big Five personality traits. As with this study, Gupta and colleagues presented both ancestry-specific EUR and AFR and transancestry GWAS; for straightforwardness in comparison, we examined whether any of the lead SNPs across any of their analyses were significant in any of our analyses. If a lead SNP that they identified was genome-wide significant (GWS) in any of our analyses, or in the same 250kb genomic locus as a significant SNP in any of our analyses, we consider this a successful replication. Notably, the majority of Gupta and colleagues' sample comes from the Million Veterans Project cohort, and, for neuroticism, the UK Biobank cohort, which are also included in our data. Additionally, we note one point of methodological divergence between the two studies: Gupta and colleagues included multiallelic SNPs (i.e. a SNP with multiple reference alleles) in their GWAS, whereas we did not. Thus, we quantify the replication rate among biallelic SNPs.

In **Supplementary Table S12**, we present the results of our replication analyses, including the effects of all lead SNPs in Gupta and colleagues (2024), the effects of those SNPs in our study, and the effects of significant SNPs in the same locus in our study, if applicable. Overall, 207 of the 246 lead biallelic SNPs in Gupta and colleagues replicated in our study (84%): 9 of 12 extraversion lead SNPs, 3 of 3 agreeableness lead SNPs, 2 of 3 conscientiousness lead SNPs, 192 of 224 neuroticism lead SNPs, and 3 of 7 openness to experience lead SNPs. Among the 39 lead SNPs that did not replicate, the median *p*-value in our transancestry analysis was $9.05\times10^{-7}$, indicating that cases of non-replication were typically with SNPs whose *p*-values were just below the genome-wide significance threshold in our data.

2.9 Identifying Novel SNPs

We quantified whether lead SNPs indexed significant signal at a novel genomic locus or replicated significant signal found in a previous study. We aggregated lead and GWS SNPs (whichever were available) from previous Big Five GWAS studies with significant results: de Moor and colleagues (2012; including SNPs that were GWS in their discovery but not replication sample), Lo and colleagues (2017), Nagel and colleagues (2018; neuroticism only), Luciano and colleagues (2018; neuroticism only), Wu and colleagues (2024), and Gupta and colleagues (2024; including results significant in EUR, AFR, and transancestry analyses). We identified whether any of these GWS SNPs identified in past research were in the same 250kb locus as a GWS SNP in our results (Watanabe et al., 2017). If not, we consider this lead SNP to be novel.

Sixty-five percent of the lead SNPs – 823 of 1,257 – identified in this study were novel. in **Supplementary Tables S5-S9**, we categorize each lead SNP in this study as novel or replicating a previous finding. Most findings were novel, which is expected considering the comparatively low power of prior Big Five GWAS analyses. Specifically, for extraversion, 233 lead SNPs were in novel loci, and 25 replicated previously-identified significant loci. For agreeableness, 30 lead SNPs were in novel loci, and 9 replicated previously-identified significant loci. For conscientiousness, 129 lead SNPs were in novel loci, and 2 replicated previous loci. For neuroticism, the trait examined with the largest sample size in prior GWAS studies, 309 lead SNPs were in novel loci, and 394 replicated previous loci. Finally, for openness to experience, 122 lead SNPs were in novel loci, and 4 replicated previous loci.

2.10 Quantifying Overlap Among the Big Five

Finally, we quantified overlap in genetic signal across the Big Five, in two ways. First, we tallied the number of genetic loci tagged by the lead SNPs in analyses of each trait, and then we examined the overlap among those loci. As above, a genetic locus was defined as the 250kb region surrounding each lead SNP. Of the 931 total genomic loci tagged by the 1,257 lead SNPs in this study, 762 loci were unique to a single trait and did not overlap with any significant loci of other traits (**Supplementary Table S13**). The major exception to this general pattern of independence between the Big Five: a locus on the FOXP2 gene was associated with lower extraversion and neuroticism, and higher agreeableness and conscientiousness; this locus seems to be highly pleiotropic and has also been associated with borderline personality disorder (Streit et al., 2024).

We also quantified the general extent of genetic sharing genome-wide across the Big Five by estimating bivariate LD Score Regression (LDSC) associations across traits using data from EUR participants. LDSC is explained in full in sections 3.1 and 3.3 below. We standardized these genetic correlations using the rgmodel() function within Genomic SEM which provides SEs and p values of the true genetic correlation, as opposed to the standardized genetic covariance (Ennis et al., 2025; Tan et al., 2024). Results are presented in **Figure 1** in the main text. In general, the Big Five were mostly independent of one another, with similar correlations to those typically found in phenotypic research (e.g., Soto & John et al., 2017). As with phenotypic correlations, we found small but significant genetic correlations between agreeableness, conscientiousness, and low neuroticism, which is often referred to as the stability superfactor, and small but significant genetic correlations between openness and extraversion, often referred to as the plasticity superfactor (Digman, 1997; DeYoung, 2015). Finally, we identified a negative genetic correlation between extraversion and neuroticism, which is commonly found in phenotypic research (e.g., Soto & John, 2017). Overall, the low degree of overlap among the Big Five at the genetic level substantiates that these traits should be considered separately, as each indexes unique vectors of genetic variation. For further information on cross-Big-Five correlations, see section 6.5 below.

#### 3. Characterizing Common-Variant Heritability

3.1 Estimating SNP Heritability with Linkage Disequilibrium Score Regression

We estimated the SNP heritability ($h^2$SNP) of each personality trait using the univariate GenomicSEM extension of Linkage Disequilibrium Score Regression (LDSC; Bulik-Sullivan et al., 2015; Grotzinger et al., 2019). LDSC estimates the proportion of variance in a highly polygenic phenotype indexed by a set of common variants across the genome by regressing SNP GWAS effect sizes against their LD scores. SNPs with higher LD scores have proportionally higher odds of tagging a causal variant, so the slope of this regression indexes the heritability of a phenotype. Confounding due to population stratification and cryptic relatedness inflates GWAS effect sizes uniformly across the genome, so the intercept of this regression indexes and accounts for confounding (Bulik-Sullivan et al., 2015).

The SNPs included in LDSC analyses were the 1.2 million available in the HapMap3 reference panel. Because LD scores vary substantially across groups of people with different recent ancestral histories, these analyses were conducted with participants stratified into EUR or AFR groups. For analyses of EUR participants, LD scores were estimated from 1000 Genomes phase 3v5 EUR sample (Bulik-Sullivan et al., 2015). For AFR participants, we followed the same procedure using the 1000 Genomes phase 3v5 AFR sample to estimate LD scores.

First, we applied LDSC to estimate the $h^2$SNP of the meta-analytically synthesized GWAS results. We filtered SNPs to those with sample-size weighted meta-analytic MAF $\geq .01$ and INFO $\geq .90$ (equation 6, above). Because these GWAS results were synthesized using a fixed-effects framework, the $h^2$SNP of these associations indexes genetic effects that are shared across contributing studies, discarding the unique heritable variance in each study that is not shared with others.

As we discuss in the main text, LDSC estimated $h^2$SNP among EUR participants ranged from 4.9% (agreeableness) to 9.3% (extraversion). In contrast, $h^2$SNP among AFR participants was lower in magnitude, ranging from 1.6% (conscientiousness) to 5.1% (neuroticism; **Supplementary Table S4**).

To rule out that these differences in heritability were attributable to differences in samples across the two analyses, we re-ran $h^2$SNP analyses among the subset of 10 cohorts that ascertained both EUR and AFR participants (**Supplementary Table S15**). Even in this matched set of cohorts, the lower-bound of the 95% confidence interval of $h^2$SNP among EUR participants was greater than the upper bound of the 95% confidence interval of $h^2$SNP among AFR participants for extraversion (EUR 9.3% vs. AFR 8.0%), conscientiousness (EUR 5.4% vs AFR 5.2%), neuroticism (EUR 14.2% vs. AFR 9.4%), and openness to experience (EUR 7.6% vs AFR 6.7%). This suggests that $h^2$SNP as indexed by LDSC differs across participants with different ancestral histories. This may relate to the more complex LD structure among AFR participants compared to EUR participants, rendering directly genotyped SNPs less effective at indirectly tagging effects of non-genotyped SNPs, and to the greater amount of admixture among broadly AFR participants, which may render AFR LD scores less well-matched to the analytic sample (Bulik-Sullivan et al., 2015).

We applied Stratified LD Fourth Moments Regression (S-LD4M; O'Connor et al., 2019) to estimate the polygenicity of SNP effects on personality among EUR participants. Using GWAS summary statistics as input, S-LD4M describes how evenly the $h^2$SNP of a trait is spread across the genome by modeling the kurtosis of SNP effect sizes. In **Supplementary Table S14** we quantify the estimated effective number of independent common SNPs with effects on each trait.

3.2 Genetic Correlations Across EUR and AFR participants

We estimated transancestry LD scores and within-trait transancestry genetic correlations using POPCORN (Brown et al., 2016). Transancestry LD scores were calculated for 4,607,783 SNPs that were available in both the AFR and EUR ancestry groups in the 1000 Genomes Project (The 1000 Genomes Project Consortium, 2015), had a MAF > 0.01, and were not in the major histocompatibility complex region. LD scores were used as input with the EUR and AFR GWAS to estimate transancestry within-trait genetic correlations using the default regression-based model. Importantly, the regression-based genetic correlations estimated using POPCORN are not constrained between -1 and 1. The within-ancestry heritability p-value reported is for a test that the heritability is greater than 0 and the transancestry genetic correlation p-value reported is for a test that the genetic correlation is less than 1. Within ancestry heritability estimates and the transancestry genetic correlations were calculated from SNPs with available LD scores that were also present in the AFR and EUR GWAS. We computed both genetic effect and genetic impact LD scores and genetic correlations.

We present results of these analyses in **Supplementary Table S16**. All transancestry within-trait genetic correlations, for both genetic effect and genetic impact, did not differ significantly from 1; point estimates for correlations of genetic impact ranged from 0.635 (SE = 0.324) for neuroticism to 1.586 (SE = 6.965) for agreeableness. The wide range of plausible values was likely driven by the comparatively small sample size of analyses among AFR participants, highlighting the continued need for GWAS sample size increases in non-EUR participants.

3.3 Quantifying Heritability Across Cohorts with Random-Effects Meta-Analysis and Weighted Meta-Regression

We applied LDSC to estimate the heritability of each trait in each individual EUR cohort, leveraging the diversity of contributing cohorts with potentially-varying heritability estimates. We filtered SNPs in each cohort to those with MAF ≥ .01 and INFO ≥ .90. LDSC intercept and heritability estimates for each trait and each cohort can be found in **Supplementary Table S1**; . We note that for smaller cohorts (roughly, N < 10,000), heritability estimates have especially large standard errors; we caution against over-interpretation of estimates in each individual cohort.

We then applied fixed-effects and random-effects meta-analysis across heritability estimates for these cohorts. In a general sense, fixed-effects meta-analyses aggregate information about a parameter under the assumption that estimates from different studies are drawn from a single distribution with one true parameter, whereas random-effects meta-analyses relax this assumption to accommodate the possibility that effects are drawn from multiple distributions (Borenstein et al., 2007). We present these estimates, alongside LDSC $h^2$SNP estimates, in **Supplementary Table S17** (See **Supplementary Table S4** for complete results of meta-analytically combined LDSC $h^2$SNP estimates.

Across traits, LDSC and fixed-effects heritability estimates were highly similar, reflecting the similar set of assumptions underlying these fixed-effects models. In contrast, heritability as indexed by random-effects meta-analysis sometimes varied from these fixed-effects estimates: For agreeableness, random-effects heritability was significantly larger than LDSC-estimated heritability (.074, SE = .010 versus .049, SE = .002), indicating heterogeneity in heritability estimates across contributing cohorts. The variability in heritability across cohorts, quantified in a random-effects model using the $\tau$ parameter, ranged from .019 for neuroticism to .046 for openness to experience.

Foundational psychometric theory posits that measurement error attenuates a measure's statistical signal (Spearman, 1910; Novick et al., 1971) This has previously been demonstrated for the heritability of personality using twin samples (Jang et al., 1998): less reliable scales tend to be less heritable, and scales with greater reliability tend to be more heritable. $h^2$SNP estimates across many cohorts that measured personality using differentially reliable instruments allowed us to test for the first time whether there is an association between a personality measure's reliability and its $h^2$SNP.

To test this, we estimated six weighted meta-regressions where the $h^2$SNP in each cohort was regressed on the reliability of its personality measure, indexed in terms of Cronbach's alpha and accounting for each cohort's relative power. Specifically, for each of the Big Five traits, we estimated the following equation:

$$h^2_{SNP} = b_0 + b_1 \times (\alpha - 1) + e \, , \qquad \text{(Eq. 8a)}$$

with each estimate weighted according to

$$weight = \frac{1}{\mathrm{SE}^2_{h^2}} \, . \qquad \text{(Eq. 8b)}$$

In this equation, $B_0$ is an intercept describing the trait's heritability at an alpha of 1.0, $B_1$ is a slope parameter describing the association between variability in the scale's internal consistency ($\alpha$) minus 1 and its heritability, and $h^2SE^2$ is the SNP heritability standard error for each cohort. We also estimated a sixth meta-regression using the lme4 package in R (Bates, 2010), where we predicted heritability from internal consistency in a mixed linear model with a random intercept, with heritability estimates for each trait nested within studies, and estimates weighted by both the inverse sampling variance of the heritability estimates and the inverse number of estimates contributed by that cohort, to appropriately account for cohorts that contributed multiple estimates across traits, as in the following equation:

$$h^2_{SNP\, i,j} = B_{0,j} + B_1\left(\alpha_{ij} - 1\right) + e_{ij} \quad . \qquad \text{(Eq. 9a)}$$

where $i$ is each individual $h^2$SNP estimate and $j$ is each cohort, with each estimate given weight that is inversely proportional to the number of Big Five traits it measured ($Ntraits$), ranging from 1 to 5:

$$weight = \frac{1}{h^2SE^2 \times t\ traits} \, . \qquad \text{(Eq. 9b)}$$

Specific $\alpha$ estimates for each trait measured with each scale can be found in **Supplementary Table S1**. We note that for many scales, estimates of alpha reliability were obtained from externally published scale validation estimates using different cohorts, introducing measurement error that likely attenuated meta-regression associations somewhat.

As shown below in **Supplementary Figure S11**, we identified a robust association between a measure's reliability and its $h^2$SNP: across traits, slope $b$ = .067 (SE = .007), for extraversion, $b$ = .137 (.017), agreeableness, $b$ = .079 (.014), conscientiousness, $b$ = .082 (.018), neuroticism, $b$ = .128 (.023), and openness, $b$ = .116 (.017).

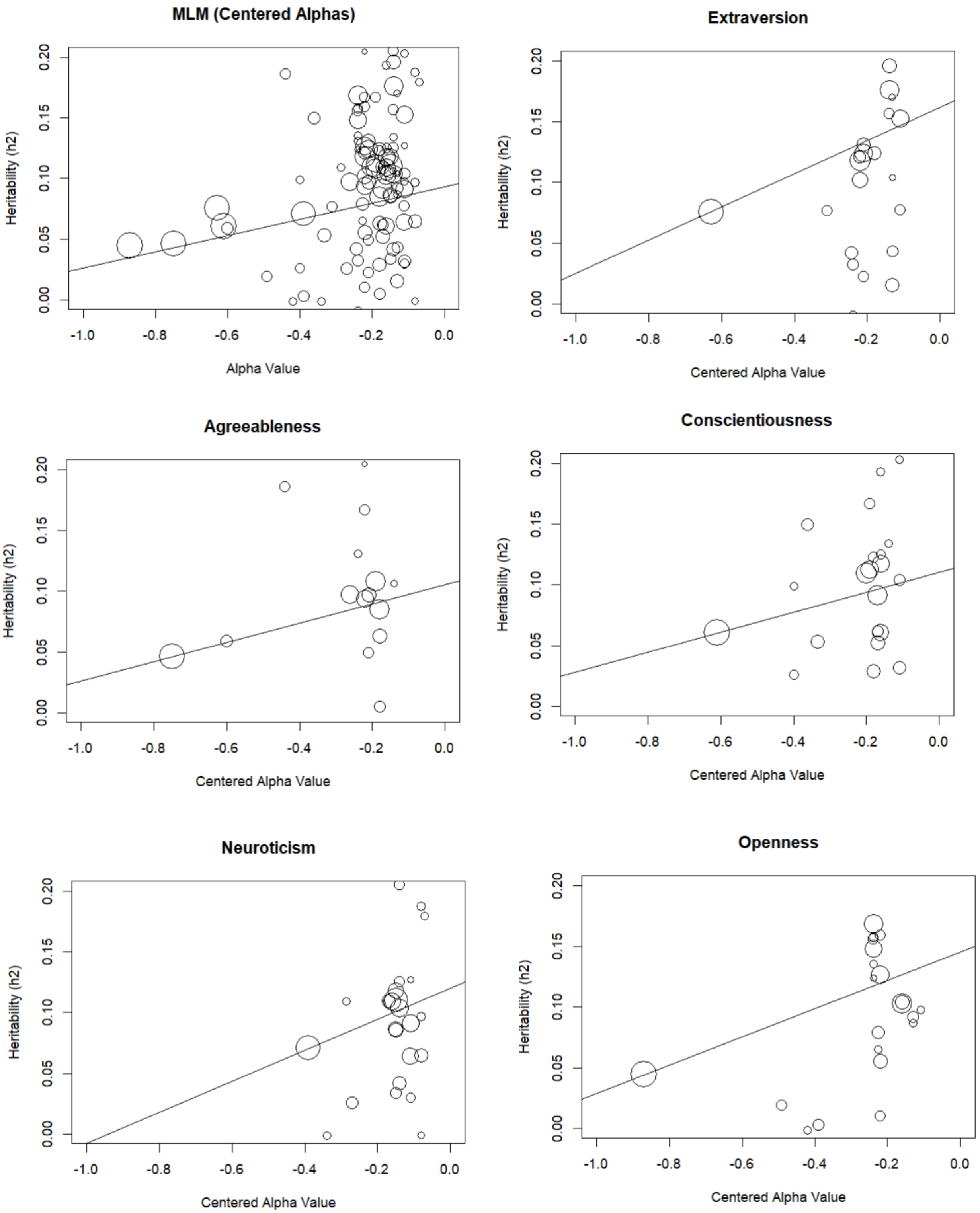


**Supplementary Figure S11. Meta-regressions of SNP heritability against scale reliability**

**across cohorts.** Note: MLM = Multilevel Model. Centered alpha values were estimated by subtracting 1 from all alpha values, such that an alpha of 0.0 represents perfect scale reliability.

Because we re-scaled the X axis by subtracting 1 from alpha reliability estimates, the Y-intercept is transformed to an estimate of $h^2$SNP if scale reliability = 1, with a corresponding standard error estimate. These results suggest that a scale disattenuated of measurement error would have $h^2$SNP of, for extraversion, 16.2% (SE = 1.0%), agreeableness, 10.5% (1.0%), conscientiousness, 11.0% (1.0%), neuroticism, 12.0% (0.8%, openness to experience, 14.5% (1.2%), and across traits, 9.3% (1.0%).

Overall, these findings indicate that the $h^2$SNP of the Big Five traits among EUR participants varies substantially across combinations of cohorts and traits, and covaries systematically with measurement error. On the low end, the heritability of meta-analytically synthesized agreeableness data was 4.9%. On the high end, the heritability of a perfectly reliable extraversion scale was 16.2%. This variability complicates the possibility and utility of identifying a single "true" $h^2$SNP for the Big Five personality traits. We note that future research should pay close attention to the reliability of measurement instruments and, when possible, measure personality with instruments that maximize reliability.

##### 3.4 Characterizing Enrichment of Heritable Signal

To further characterize personality's genetic architecture, we submitted EUR GWAS summary statistics to MAGMA (de Leeuw et al., 2015) in order to identify annotations enriched for heritable signal, for each Big Five trait. We filtered SNPs to those with INFO ≥ .80. The gene sets included in this analysis came from Akingbuwa and colleagues (2022), which include 1) protein-truncating variant (PTV)-intolerant (PI) genes ascertained by selecting genes with pLI>0.9 from the Genome Aggregation Database (gnomAD), which are under stringent selection, 2) genes expressed in brain cell types and neurons, ascertained from human and mouse brain tissue, 3) the intersection of PI genes and brain-expressed genes (i.e., PI genes expressed in the brain), and 4) gene sets defined by synaptic processes and cellular composition. Gene sets and their sources and or construction are described in full in Akingbuwa et al., 2022.

In **Supplementary Tables S18-S22**, we present results of analyses for each combination of trait and gene set, and in **Supplementary Figures S12-S16** below, we visualize those results. In general, number of significant enrichment sets scaled with sample size and heritability: there were more significant associations for neuroticism and fewer for agreeableness. Content-wise, the intersection of brain genes and genes intolerant to protein truncation showed stronger enrichment than just brain genes. These patterns have previously been found for psychiatric disorders, such as schizophrenia (Akingbuwa et al., 2022), implicating shared biological pathways in the etiology of personality and psychopathology. Additionally, these results replicate findings from previous studies which find that neuroticism heritability is enriched among excitatory and inhibitory neurons in the brain (Bryois et al., 2020) and extend them to the other Big Five traits. We further characterize these enrichment findings in the main text.

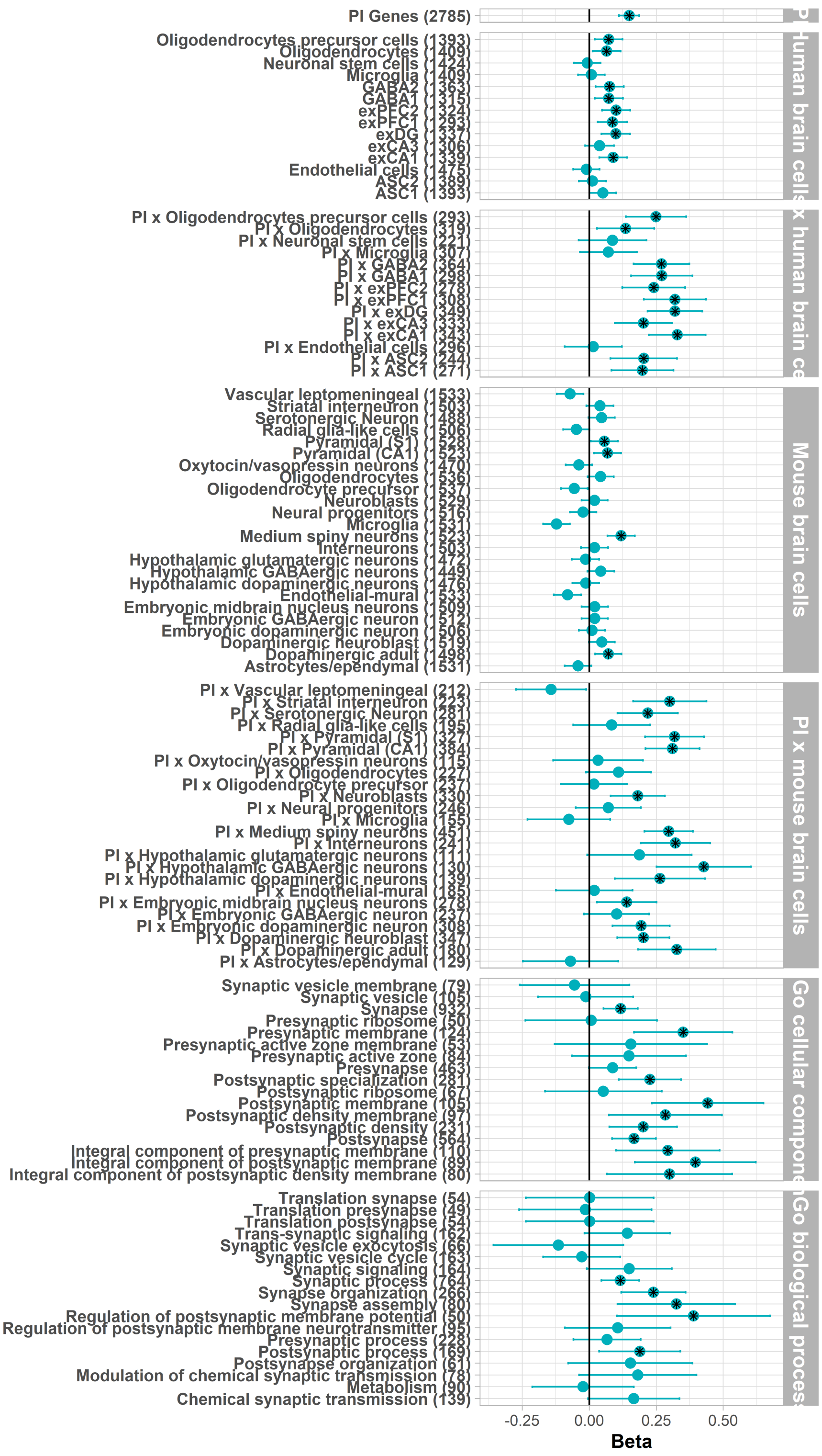


**Supplementary Figure S12. Extraversion gene-set heritability enrichment analyses**. PI = Protein-Truncating Intolerant genes. An X indicates the intersection of two gene sets. Number of genes in each set are in parentheses. Associations with a black star are significant after false-discovery-rate correction (Benjamini & Hochberg, 1995).

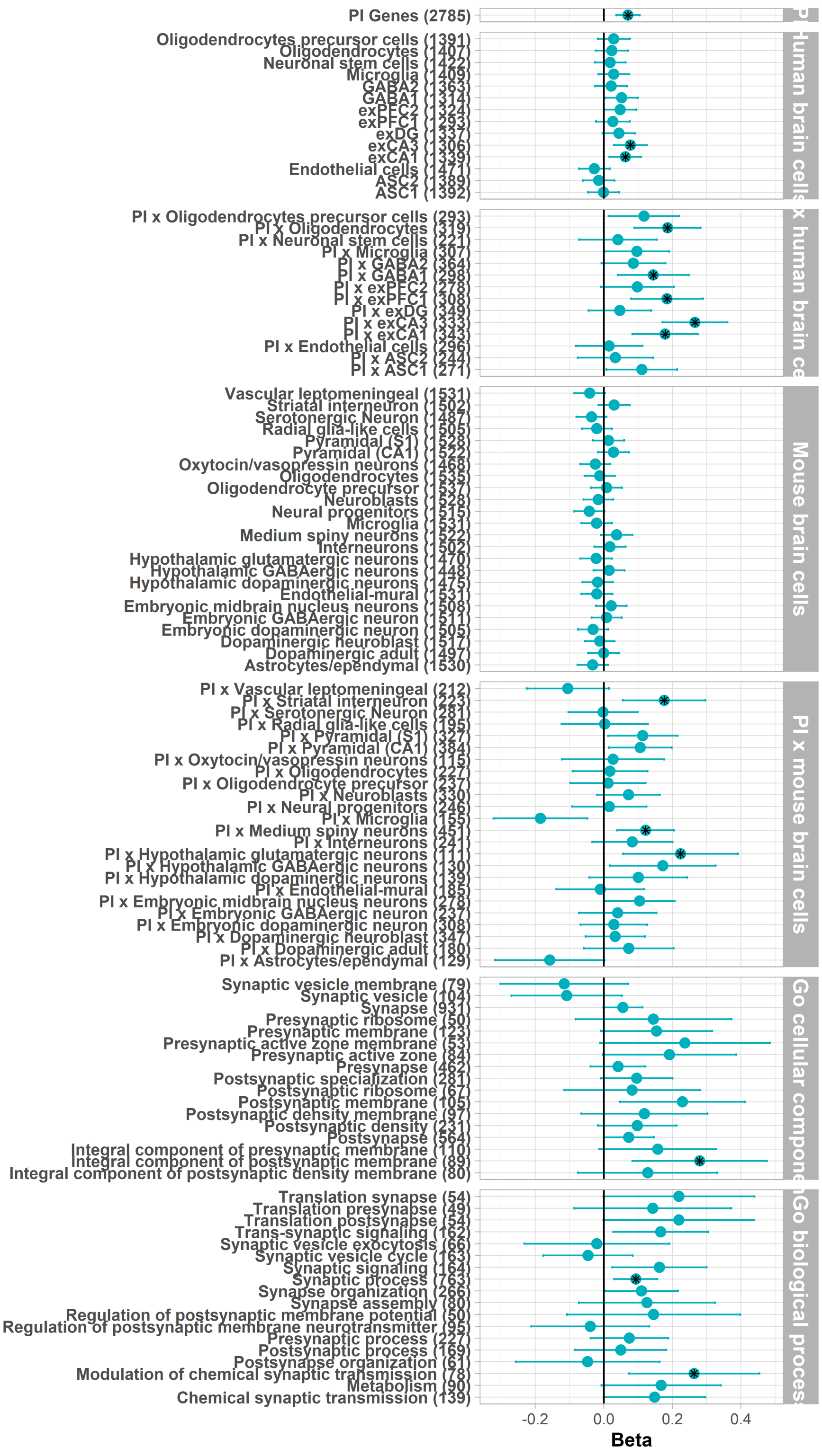


**Supplementary Figure S13. Agreeableness gene-set heritability enrichment analyses**. PI = Protein-Truncating Intolerant genes. An X indicates the intersection of two gene sets. Number of genes in each set are in parentheses. Associations with a black star are significant after false-discovery-rate correction (Benjamini & Hochberg, 1995).

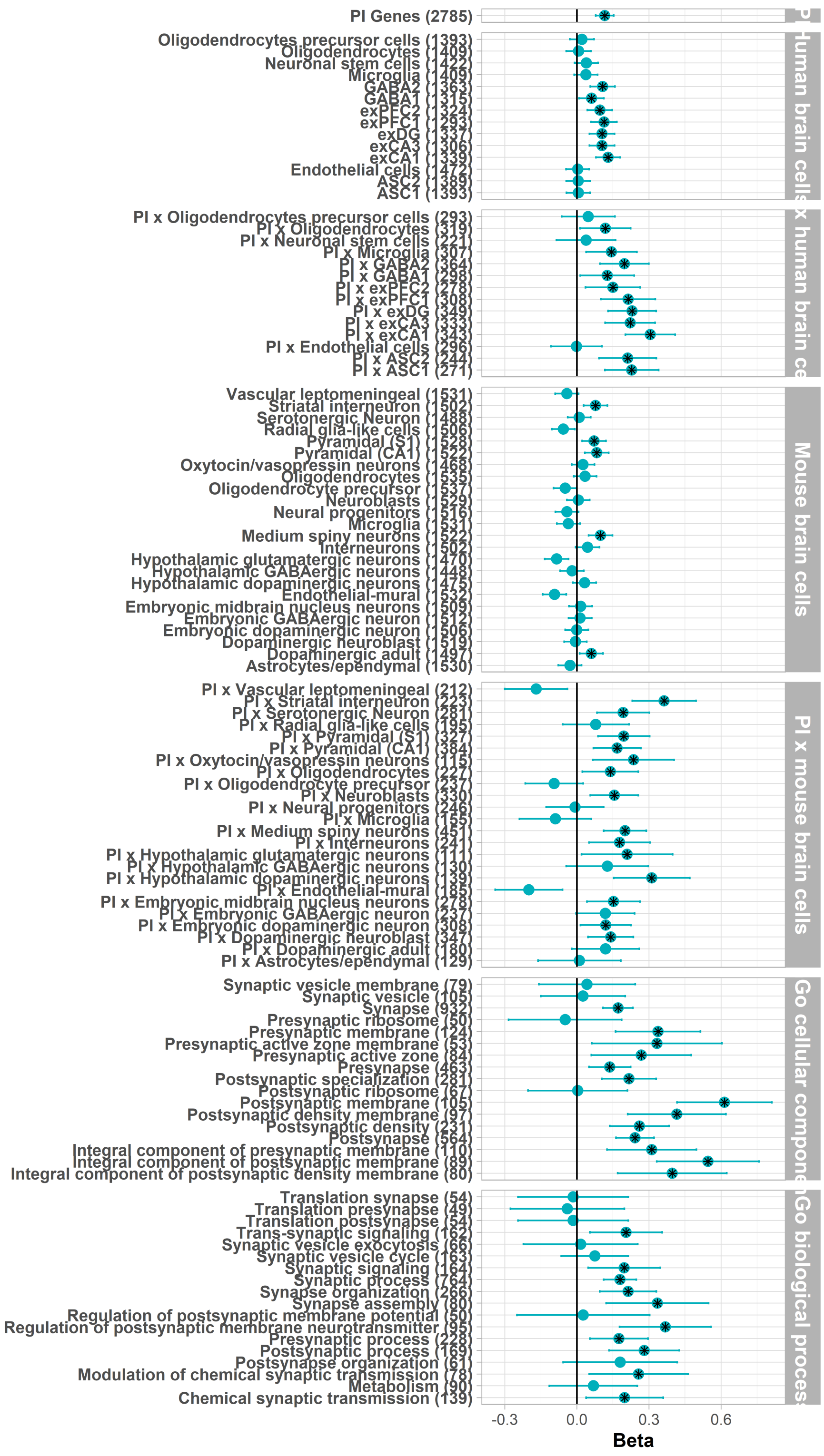


**Supplementary Figure S14. Conscientiousness gene-set heritability enrichment analyses**. PI = Protein-Truncating Intolerant genes. An X indicates the intersection of two gene sets. Number of genes in each set are in parentheses. Associations with a black star are significant after false-discovery-rate correction (Benjamini & Hochberg, 1995).

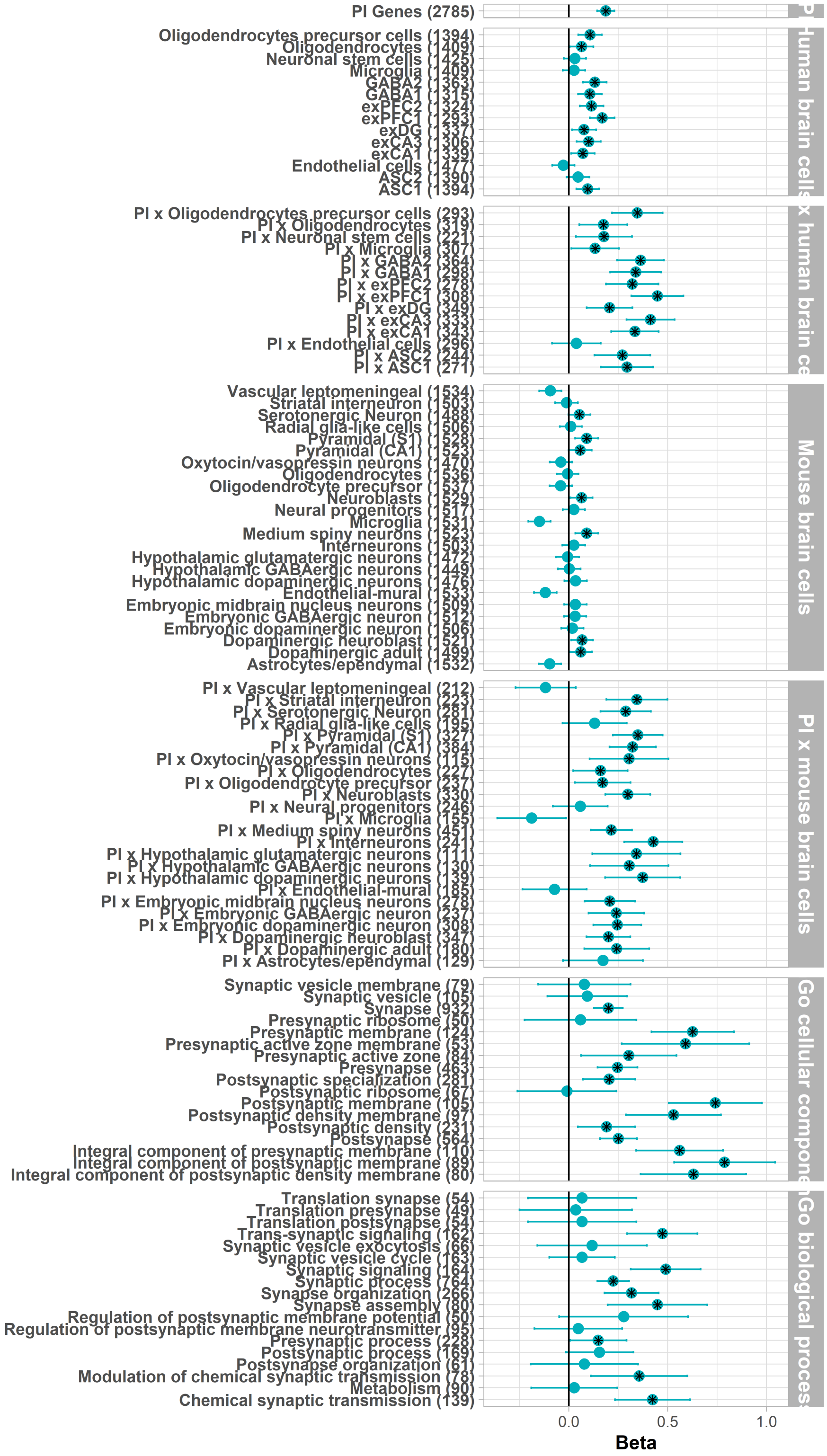


**Supplementary Figure S15. Neuroticism gene-set heritability enrichment analyses**. PI = Protein-Truncating Intolerant genes. An X indicates the intersection of two gene sets. Number of genes in each set are in parentheses. Associations with a black star are significant after false-discovery-rate correction (Benjamini & Hochberg, 1995).

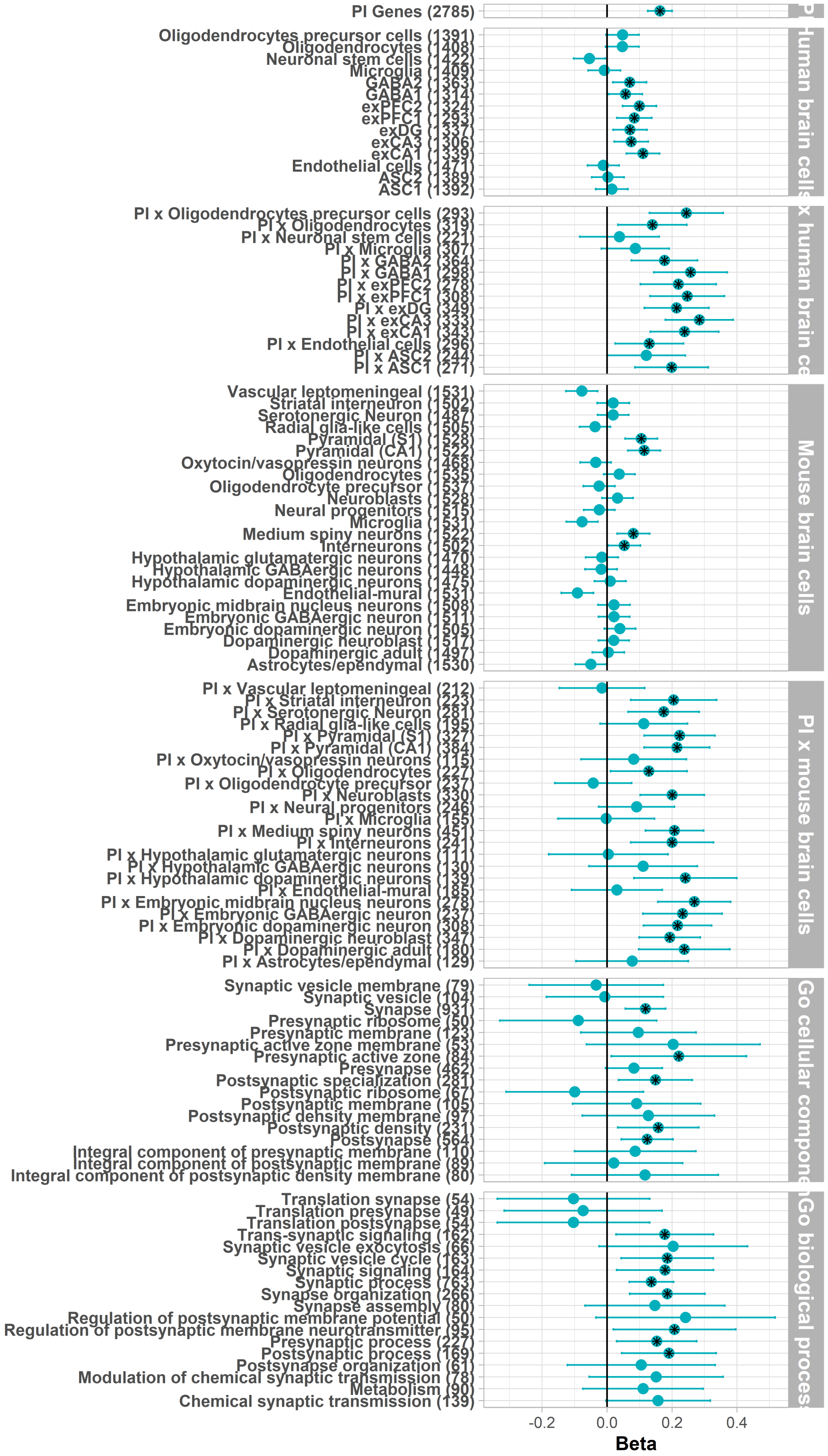


**Supplementary Figure S16. Openness to experience gene-set heritability enrichment analyses**. PI = Protein-Truncating Intolerant genes. An X indicates the intersection of two gene sets. Number of genes in each set are in parentheses. Associations with a black star are significant after false-discovery-rate correction (Benjamini & Hochberg, 1995).

We then indexed the similarity of enrichment across the Big Five, using pairwise rank correlations of enrichment betas from the analyses presented above. We visualize these correlations in **Supplementary Figure S17**. In general, patterns of enrichment were similar across the Big Five, though enrichment was less similar for pairs of traits involving agreeableness and openness to experience, especially for the pairing of openness to experience and agreeableness ($\rho$ = 0.48).

We also calculated z-statistics across the 112 gene sets to test for significant differences in annotation enrichment between pairs of traits. We found that agreeableness and neuroticism had the most significantly differently enriched gene sets (46), with extraversion and openness to experience the fewest (4). Agreeableness and neuroticism generally showed the most differentiated enrichment with the other traits. In terms of most differentiated gene set pairings for each Big Five trait, agreeableness showed the most differential enrichment with neuroticism (46 gene set annotations), conscientiousness also with neuroticism (19), extraversion with agreeableness (18), neuroticism with agreeableness (46), and openness to experience with neuroticism (19). Finally, in terms of similarity, agreeableness was most similar to conscientiousness (number of differentially enriched gene sets = 13), conscientiousness to extraversion (6), extraversion to openness to experience (4), neuroticism to extraversion (9), and openness to experience to extraversion (4).

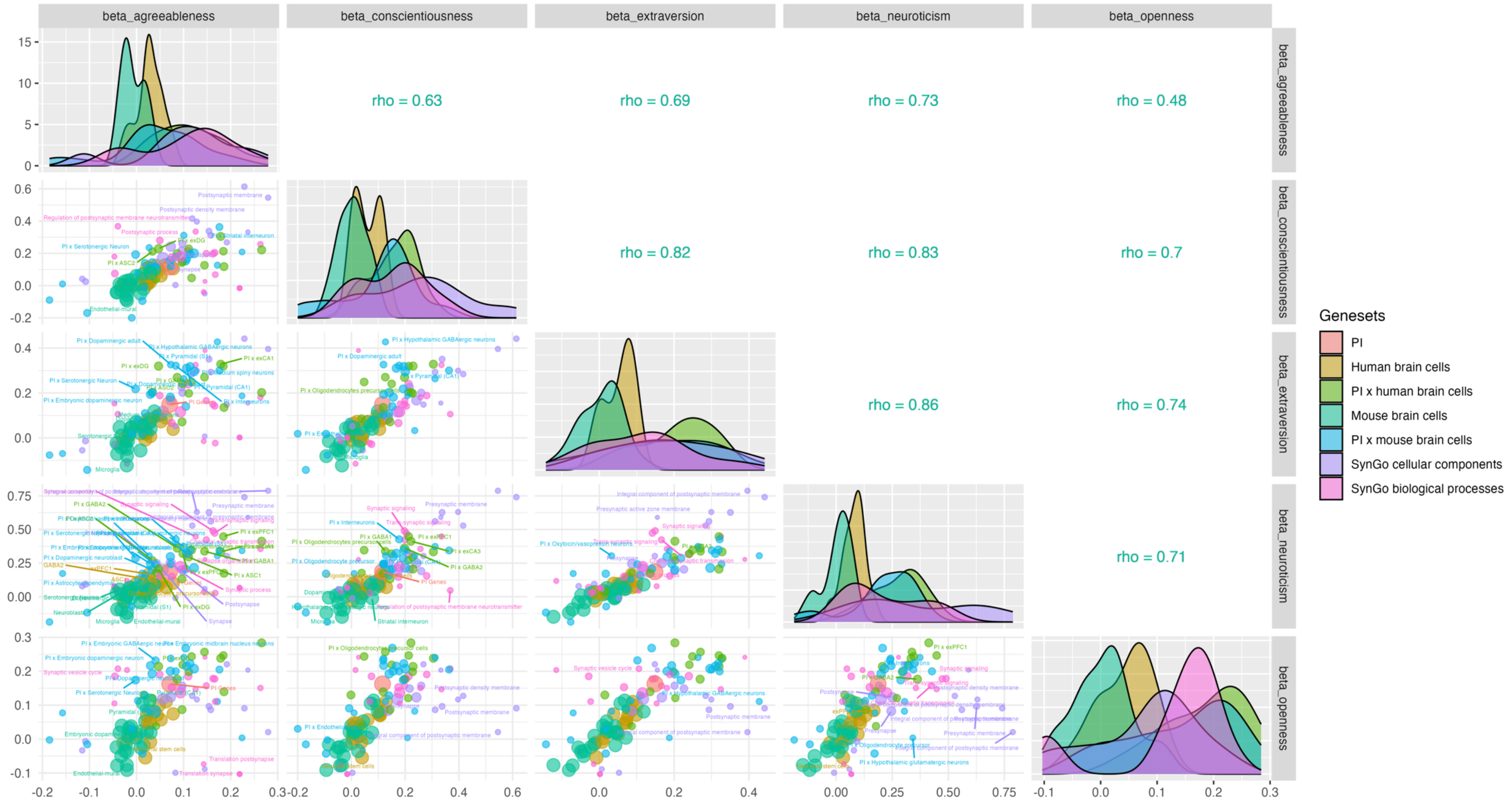


**Supplementary Figure S17. Pairwise rank correlations of enrichment betas across the Big Five.** Note: PI = Protein-truncating Intolerant genes. An X indicates the intersection of two gene sets. Larger points have larger weights, scaled by the inverse product of the standard errors of the gene set association betas. Gene sets with significantly differing enrichment across pairs of traits are labelled.

##### 3.5 Comparison of Genetic Architecture Across Grouping Variables

We compared the genetic architecture of each of the Big Five personality traits across five vectors of variation that differed substantially across cohorts: geographic region, age group, rater perspective, military veteran status, and measurement instrument. Because genetic covariance analysis requires the highly powered GWAS from ancestrally homogeneous cohorts, these analyses were restricted to participants of European-like ancestry.

For measurement instrument, geographic region, and age group, we constructed groups for comparison using a uniform procedure. First, we identified grouping variables that maximized the number of mutually exclusive stratification groups while ensuring that 1) each group would have a sample size of over N = 15,000, allowing sufficient power for genetic correlations across groups, 2) when possible, each individual group contained multiple cohorts of participants, to minimize idiosyncratic effects of an individual cohort, and 3) when possible, each group included cohorts that varied on other grouping variables (e.g., the young age group contained cohorts that completed a variety of questionnaires and lived in different geographic regions). These grouping strategies were chosen to maximize the specificity of inference that could be drawn from comparison analyses. To compare genetic effects across rater perspectives, we contrasted Estonian Biobank self-reported GWAS estimates with Estonian Biobank other-reported GWAS estimates. To compare genetic effects across military veteran status, we contrasted Million Veteran Program GWAS estimates with GWAS estimates pooled across the other 45 cohorts.

Then, for each Big Five trait, among the cohorts included in each group, we conducted a GWAS using same method as for the main analyses. We then compared the genetic architecture across groups using bivariate LD score regression (LDSC; Bulik-Sullivan et al., 2015). In addition to quantifying $h^2$SNP for each phenotype, Bivariate LDSC quantifies the genetic covariance across two phenotypes by regressing the product of their SNP GWAS effect sizes against their LD scores. As with univariate LDSC, SNPs with higher LD scores have proportionally higher odds of tagging a variant that is associated with both phenotypes, so the slope of this regression indexes the covariance between two phenotypes; this slope can be rescaled by the heritability of both phenotypes (i.e., by each phenotype's variance) so that it is expressed as a genetic correlation that ranges from -1 to 1. Confounding due to population stratification and cryptic relatedness, and sample overlap across the two phenotypes, inflates GWAS effect sizes uniformly across the genome, so the intercept of this regression accounts for these sources of confounding and unknown and varying degrees of genetic overlap across the two phenotypes (Bulik-Sullivan et al., 2015). Critically, because LDSC examines covariance in effects across genetic variants, rather than covariance in effects across participants, associations can be estimated across mutually exclusive groups (e.g., to compare participants from the US with those from continental Europe).

First, we compared the similarity of genetic signal across geographic regions. Personality trait structure can vary across cultures and languages (McCrae et al., 2005; Saucier et al., 2014); the genetic architecture of personality may similarly vary across geographic regions. We structured our grouping across geographic regions to also stratify by language, when possible, leading to four groups: the United States (k = 12 cohorts, max N = 49,091), the United Kingdom and Australia, which both speak English but are not individually large enough to form their own group (k = 11 cohorts, max N = 500,558; max N minus the UK Biobank = 26,803), continental

Europe (k = 11 cohorts, max N = 56,510), and Nordic countries (k = 6, max N = 60,646). Details on cohort geography are available in **Supplementary Table S1.**

Results of cross-geographic comparisons (**Supplementary Figure S18**) indicated clear evidence of convergent and discriminant validity for four of the Big Five traits. The major exception was agreeableness, which was less strongly intercorrelated across groups. Among Nordic participants, agreeableness scores were strongly correlated with extraversion scores, and among UK/AUS participants, agreeableness scores were not strongly correlated with agreeableness scores in other geographic regions (we note that the Nordic-group MoBa cohort, the UK/AUS-group TEDS cohort, and the UK/AUS-group Generation Scotland cohort each provided personality data using the IPIP questionnaire, creating collinearity between cross-geography and cross-questionnaire comparisons; see cross-questionnaire comparisons below). These findings indicate that the genetic architecture of agreeableness may be particularly variable across western geography.

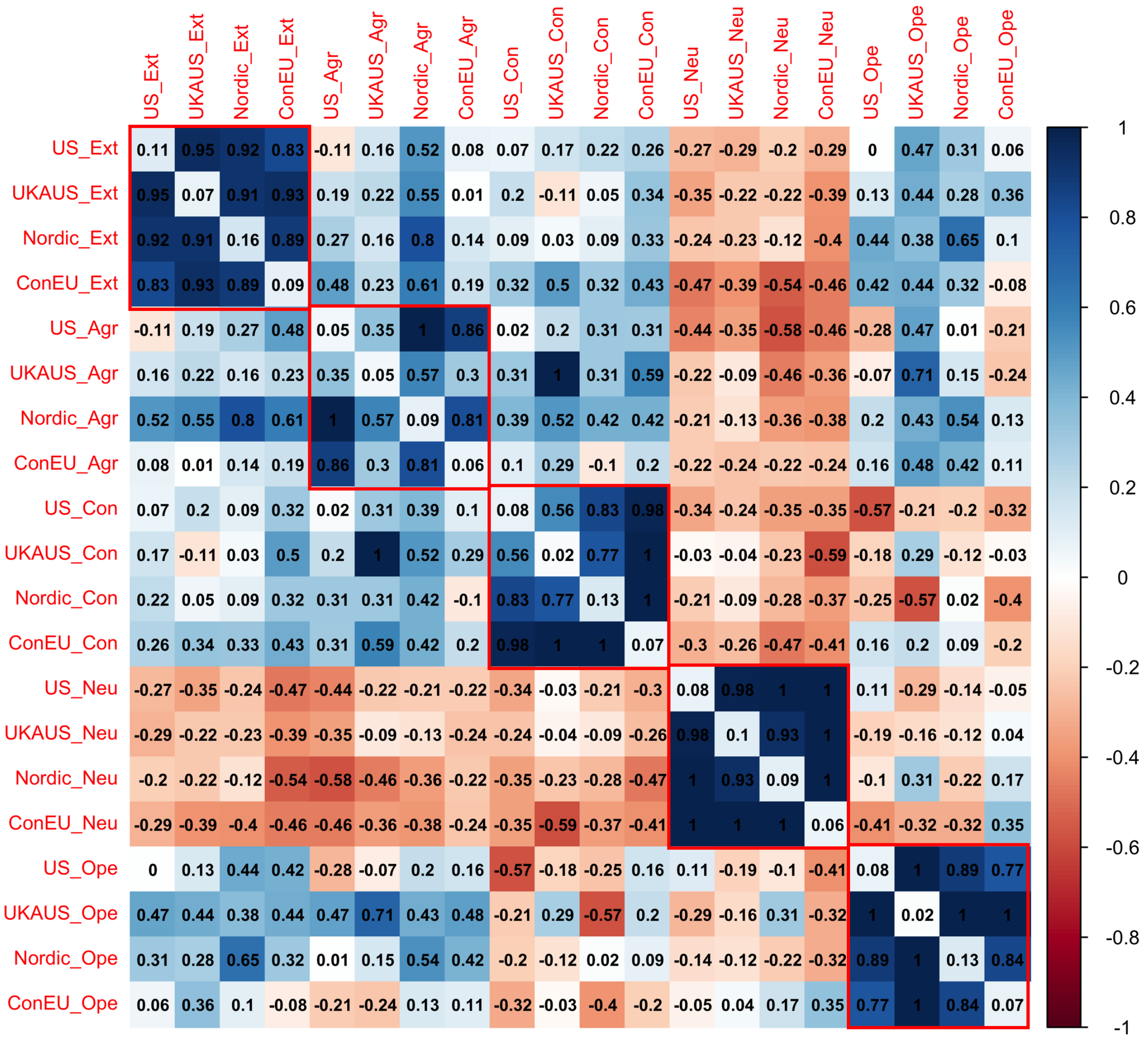


**Supplementary Figure S18. LDSC genetic correlations across geographic regions**. US = United States. UKAUS = United Kingdom and Australia. ConEU = Continental Europe.

Heritability of each scale is presented on the diagonal. Red squares denote genetic correlations among measures of the same trait.

We also compared correlations across age groups. Twin and family research has identified strong but imperfect genetic continuity in personality development across the lifespan, including a trend of decreasing heritability with age, as environmental variance increases (Tucker-Drob & Briley, 2014). For the first time, we examined whether the molecular genetic architecture followed similar developmental trends, and we conducted age-group comparisons not possible in phenotype data that requires sample overlap. Because many participating cohorts ascertained participants with wide variance in age, we grouped cohorts by comparing their mean age +/- 1.5 SD, which includes roughly 87% of participants in each cohort, and identified break points that would permit maximally large sample sizes within each age group while retaining few cohorts with participants who overlapped across age groups. The young age group contained cohorts where ~87%+ of participants were 25 or younger at mean age of personality measurement (k = 6, max N = 15,454). The middle adulthood age group contained cohorts between ages 26 and 64 (k = 6, max N = 54,414), and the older adulthood age group contained cohorts over age 65 (k = 5, max N = 18,624).

For each trait, we found general continuity in genetic architecture across age groups (**Supplementary Figure S19**), replicating results of past twin and family research (Tucker-Drob & Briley, 2014). However, unlike twin and family research, which finds decreases in broad-sense heritability with age, we did not find any systematic decreases in the $h^2$SNP of any trait with age. Given the many differences across cohorts beyond age, we refrain from drawing inferences from individual pairwise correlations. In general, these results set the stage for future investigations with fine-grained age groups and stratified GWAS within a cohort, to obtain maximally comparable estimates across groups that can be used to refine lifespan theories of personality genomics.

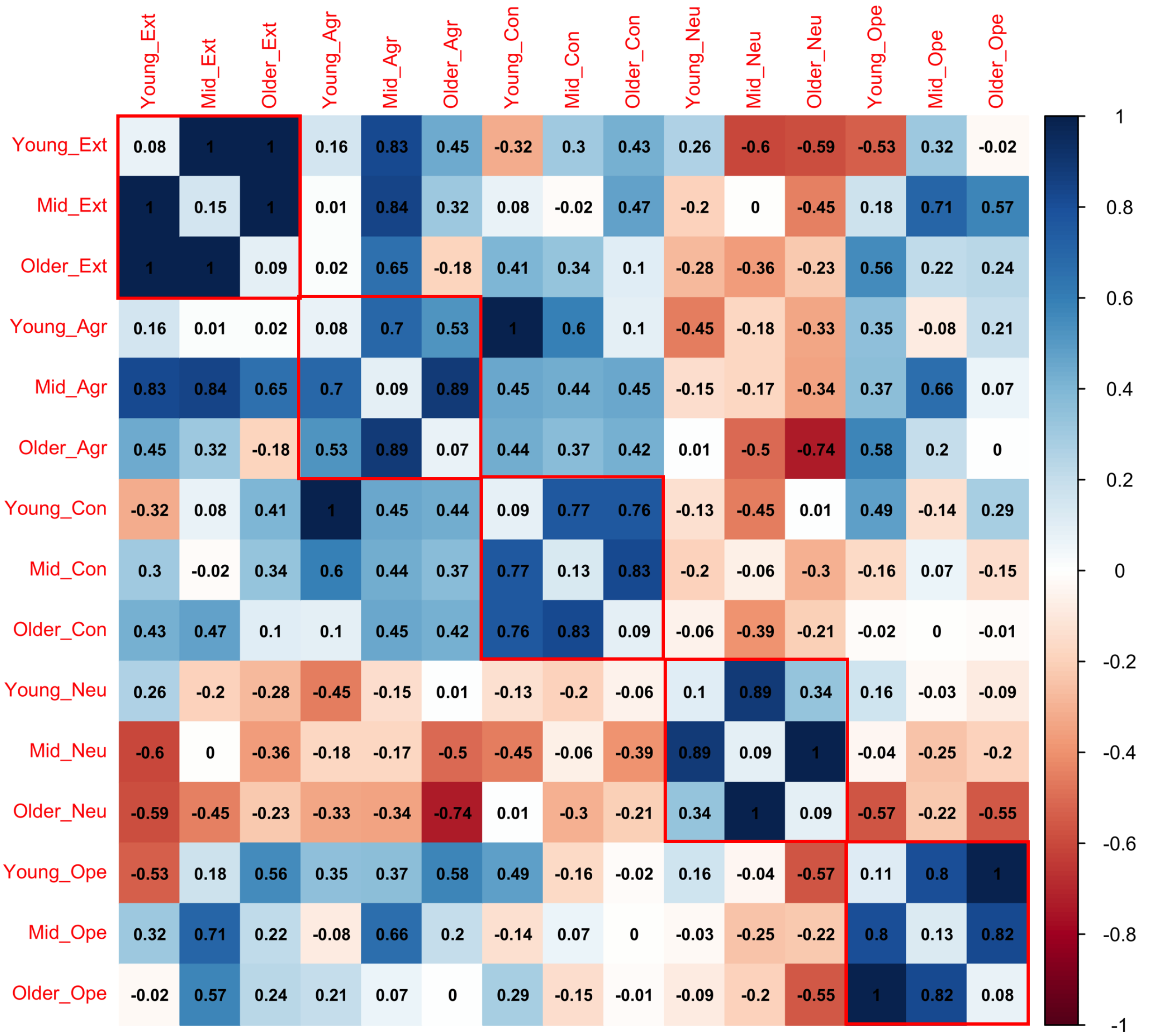


**Supplementary Figure S19**. **LDSC genetic correlations across age groups.** Note: Young = Mean cohort age +-1.5SD ≤ 25. Mid = Mean cohort age +-1.5SD = 26-64. Older = Mean cohort age +-1.5SD ≥ 65. Heritability of each trait in each group is presented on the diagonal. Red squares denote genetic correlations among measures of the same trait.

We structured our next comparison, of veteran status, to highlight the Million Veterans Project, which ascertained participants who had prior military service (Gupta et al., 2024). Notably, this cohort made up the majority of the most recently published GWAS, and so comparing the genetic architecture between the Million Veterans Project cohort and the remaining ReGPC contributing cohorts also allows us to evaluate the extent to which the genetic architecture of personality is similar to past published estimates. Importantly, there may be veterans in other ReGPC cohorts; no information was collected about veteran status in any of the other participating cohorts. Comparisons between the MVP cohort and the other 45 ReGPC cohorts illustrate general similarity across the two groups, as can be seen in the lower-left ribbon of **Supplementary Figure S20.** These correlations suggest strong similarity between Gupta and colleagues' discovery cohort and ours. Heritability was generally higher in the ReGPC cohorts than in the MVP cohort, which is expected given the comparatively low reliability of the ten-item measure administered to MVP participants.

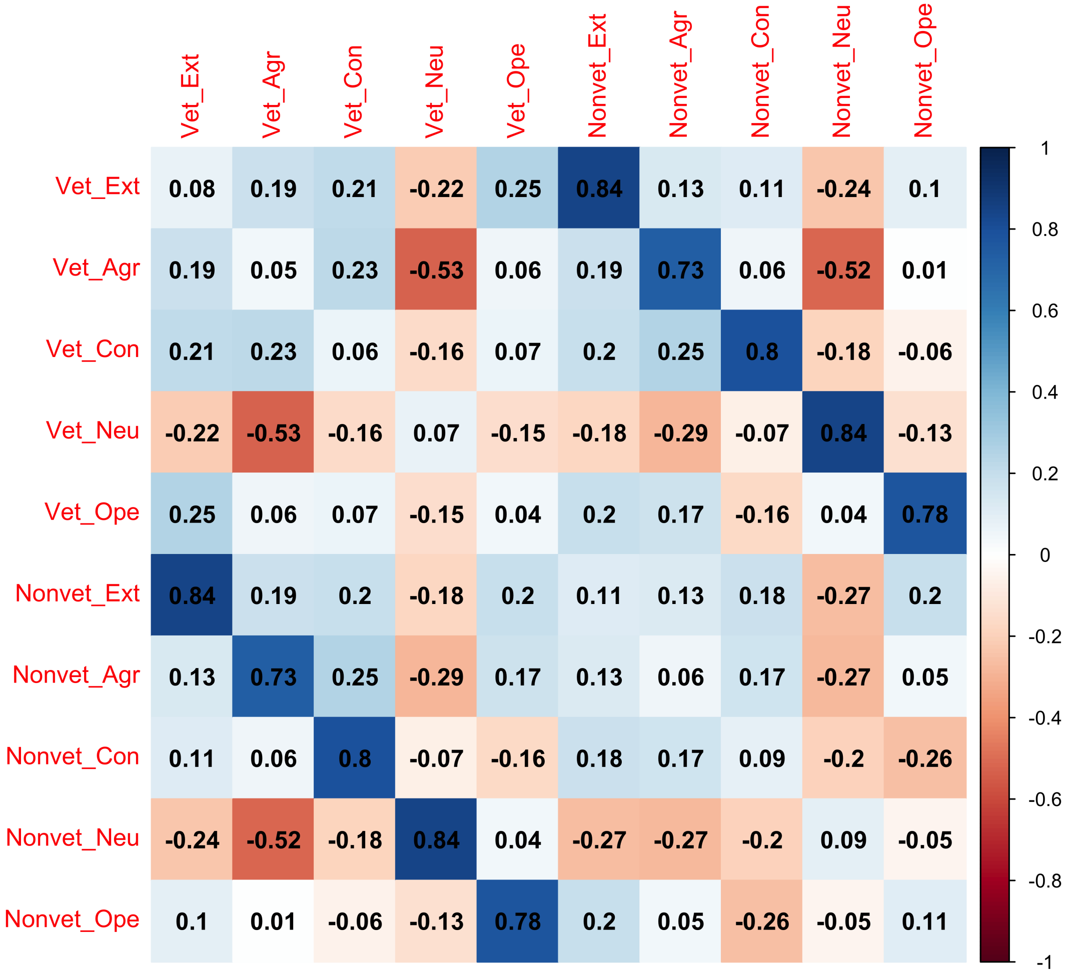


**Supplementary Figure S20. LDSC genetic correlations between the Million Veterans Program cohort and the other 45 ReGPC cohorts.** Note: Vet = Million Veterans Program. Nonvet = The Remaining 45 ReGPC Cohorts. Heritability of each trait in each group is presented on the diagonal.

We compared the genetic architecture across rater perspectives using data from the Estonian Biobank (EBB). The EBB measured personality traits from two perspectives: participants reported on their own personality traits (max N = 73,983), and they also nominated an informant to provide a second rating of their personality, from the perspective of a close other (max N = 20,269). This allowed us to examine the extent to which genetic signal was similar across rater perspectives. In general, people and close others provide similar but imperfectly correlated ratings on a target's personality (Kenny & West, 2010; Vazire, 2010). Extending this to molecular genetic data for the first time, we find generally high correlations between self-rated and other-rated genetic architectures (**Supplementary Figure S21**), especially for the traits of extraversion, agreeableness, and openness to experience. That is, people tend to see themselves quite similarly to how others see them, both phenotypically and in terms of genetic associations. These findings indicate that, though personality is typically assessed via self-report, these self-report measurements correspond with how a person is perceived by close others, reflecting a shared psychological reality.

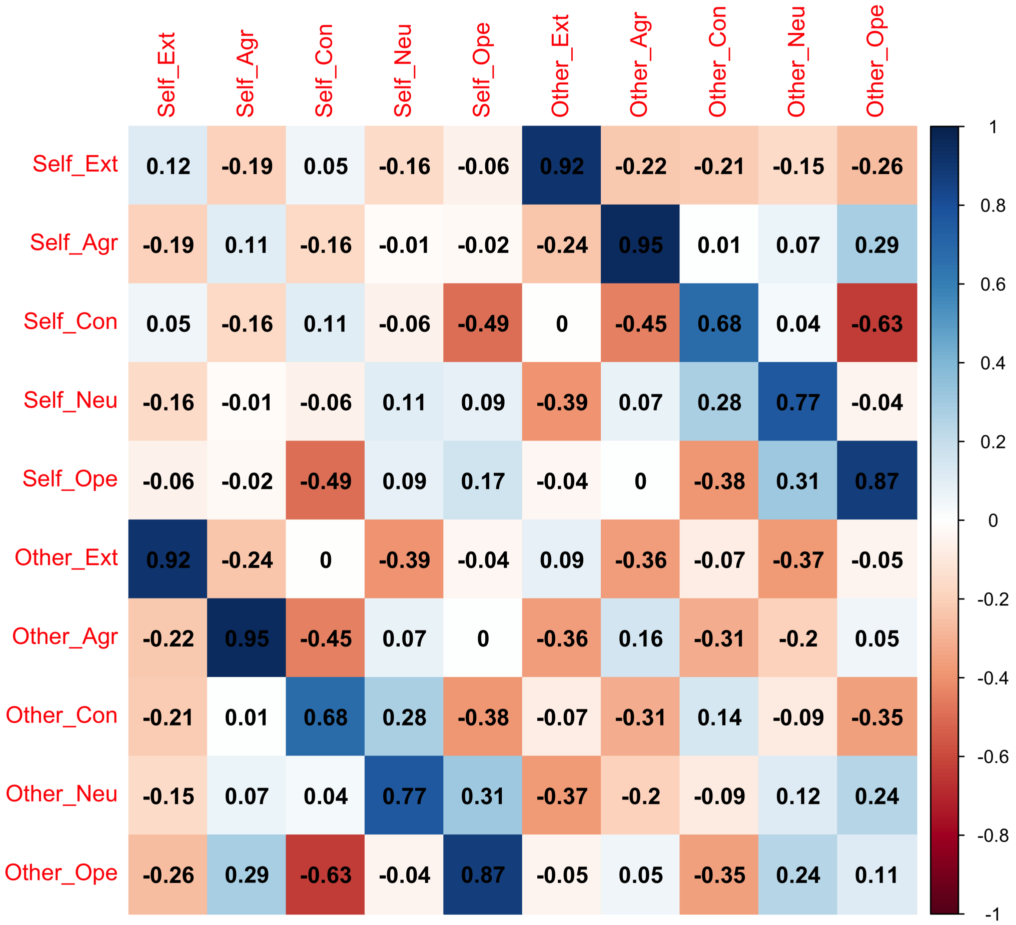


**Supplementary Figure S21. LDSC genetic correlations between self-rated and other-rated personality trait measurement in the Estonian Biobank**. Note: Heritability of each trait in each group is presented on the diagonal.

Finally, we compared the similarity of genetic signal across questionnaires. Though the Big Five is the dominant structural paradigm in modern personality psychology, content of individual questionnaires varies depending on the theories espoused by each scale's creators, with phenotypic correlations for the same Big Five trait ranging as low as $r$ = .60 across instruments (Soto & John, 2017; Thalmayer et al., 2011). Thus, it is an open question whether scales designed to measure the same trait will demonstrate high levels of genetic overlap in genomic data. Most cohorts completed personality questionnaires belonging to one of four questionnaire families. Eight cohorts (max N = 96,966) completed Big Five Inventory (BFI) scales, including the BFI and BFI-short. Four cohorts (max N = 481,307) completed the Eysenck Personality Questionnaire (EPQ) or Eysenck Personality Inventory extraversion and neuroticism scales, which are theoretically and empirically congruent with their corresponding Big Five traits. Four cohorts (max N = 47,175) completed Big Five scales drawn from the International Personality Item Pool, and 19 cohorts (max N = 117,524) completed NEO personality Inventories (NEO), including the NEO-Personality Inventory-Revised and the shorter NEO-Five Factor Inventory. Finally, Estonian Biobank participants completed the 100 Nuances of Personality inventory (max N = 73,983). Details on which cohorts were administered which questionnaires can be found in **Supplementary Table S1**.

As we show in **Supplementary Figure S22**, for extraversion, neuroticism, conscientiousness, and openness, we observed strong convergent validity ($r_g$s ~.75 and above) across personality scales measuring the same trait. For agreeableness, convergent validity across measures was moderate ($r_g$s ~ .40-.75), with the exception of the genetic correlation between IPIP agreeableness and 100NP agreeableness (0.28). We also generally observed good discriminant validity between scales measuring different traits. Exceptions included moderate genetic associations between IPIP and 100NP measures of agreeableness in relation to the extraversion scales (ranges 0.35-0.58 and -0.18-0.48, respectively). The association between IPIP agreeableness and extraversion may be explained by the inclusion of sociability-relevant content in IPIP agreeableness inventories, as exemplified by the item *"am interested in people*"). Heritability did not vary substantially across scales measuring the same trait. Overall, the pattern of genetic correlations indicates that heritable signal in the Big Five is largely robust across measurement instruments.

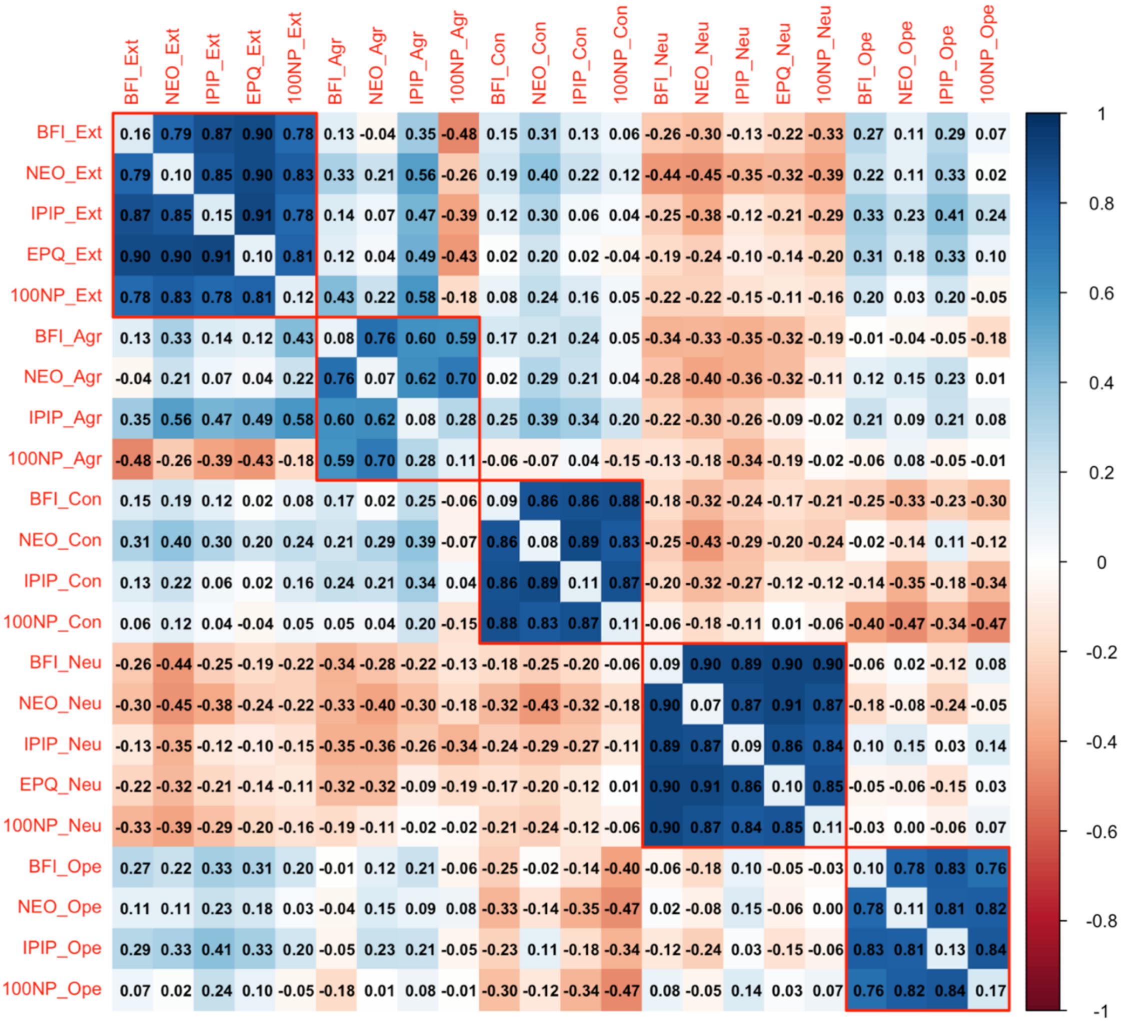


**Supplementary Figure S22. LDSC genetic correlations between personality measurement instruments**. BFI = Big Five Inventory. NEO = NEO Inventory. IPIP = International Personality Item Pool. 100NP = 100 Nuances of Personality. EPQ = Eysenck Personality Questionnaire. Heritability of each questionnaire is presented on the diagonal. Red squares denote genetic correlations among measures of the same trait.

<u>3.5 Genomic SEM Analyses Stratified by Measurement Instrument</u>

To investigate the extent to which genetic effects on personality may differ by measurement instrument, we formally modeled the multivariate genetic architecture of the 22 instrument-stratified Big Five GWAS phenotypes using Genomic Structural Equation Modeling (Genomic SEM; Grotzinger et al., 2019). We first fit a confirmatory five-factor model conforming to *simple structure,* in which each instrument loaded only on the factor corresponding to the Big Five trait that it was designed to measure, with no cross loadings modeled (**Supplementary Figure S23**). In this model, standardized loadings of each indictor on its corresponding factor were generally high, with the factors correlating only modestly. However, model fit was poor ($\chi^2(199) = 29{,}596.05$; CFI = .675; SRMR = 0.128), suggesting that additional model complexity was warranted, e.g. to reflect influences on each measure by off-target traits.

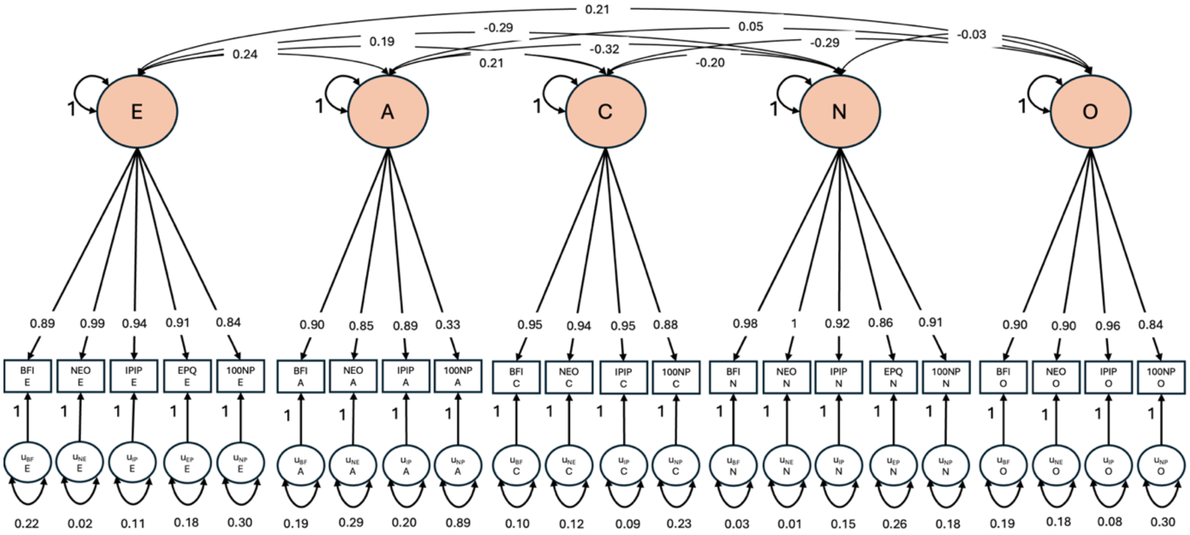


**Supplementary Figure S23. Confirmatory five-factor model of measurement-instrument stratified GWAS of the Big Five, conforming to simple structure.** Standardized estimates are displayed.

We employed an iterative data-driven approach to extend the confirmatory model to include additional free parameters. At each iteration, we inspected the residual genetic covariance matrix (calculated as the difference between the observed and model-implied genetic covariance matrices) to identify clusters of sign-consistent (all positive or all negative) residual covariances. These clusters may reflect unaccounted for dependencies between a single measure of a Big Five trait and all measures of another Big Five trait. We then modelled these dependencies by adding a corresponding cross-loading to the factor model and examined its residual covariance matrix in turn. We proceeded until there were no remaining clusters of residual genetic covariances where at least one residual genetic covariance exceeded |0.035| (which corresponds to a residual correlation of .35 for two traits with $h^2_{SNP} = .10$).

In **Supplementary Figure S24**, we display the residual genetic correlation matrix for the confirmatory simple structure model. (Note that although we used residual genetic covariances to guide model modification, we present residual genetic correlations for ease of interpretation). We observed a consistent pattern of negative residuals between 100NP agreeableness and measures of extraversion ($r_g$ = 0.56, -0.35, -0.48, -0.54, -0.25 for the BFI, IPIP, NEO, EPQ, and 100NP, respectively), suggesting that 100NP agreeableness may also reflect low extraversion. This may, in part, be attributable to the fact that 100NP agreeableness is measured solely using negatively worded items. We therefore specified 100NP agreeableness to also load on the extraversion factor and refit the model using the procedure described above.

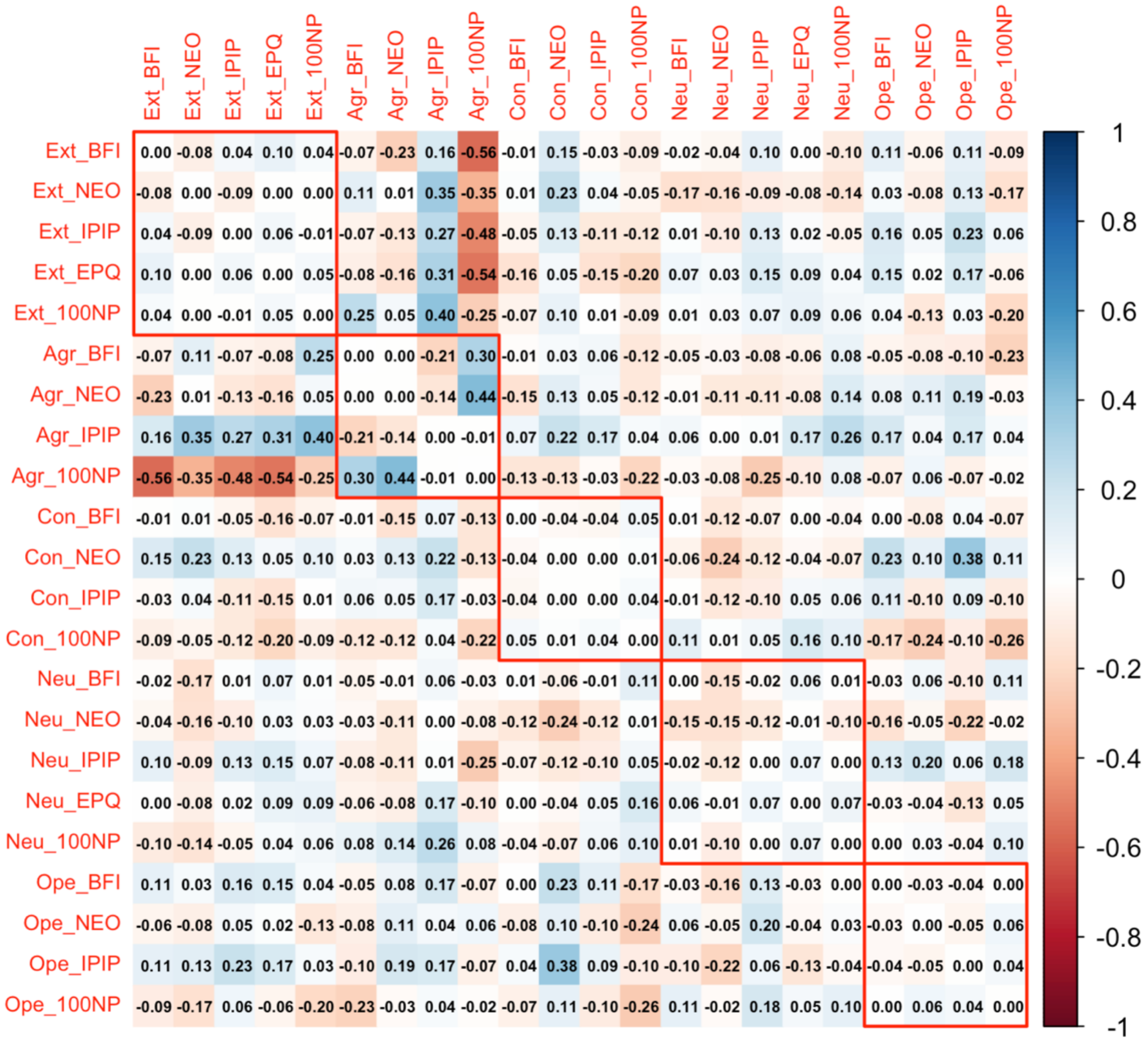


**Supplementary Figure S24. Residual Genetic Correlation Matrix for the Confirmatory Simple Structure Model.** BFI = Big Five Inventory. IPIP = International Personality Item Pool. NEO = NEO Inventory. EPQ = Eysenck Personality Questionnaire. 100NP = 100 Nuances of Personality. Red squares denote residual genetic correlations among measures of the same trait.

We display the final factor model of measurement-instrument stratified GWAS of the Big Five in **Supplementary Figure S25**, with its residual genetic correlation matrix in **Supplementary Figure S26**. As compared to the residual genetic correlation matrix displayed in **Supplementary Figure S24**, substantial clustered residuals are no longer apparent. Moreover, in comparison to the poor model fit of the simple structure model ($\chi^2(199)$ = 29,596.05; AIC = 29,704.05; CFI = .675; SRMR = 0.128), the final model displayed much improved fit for the Big Five model with cross-loadings ($\chi^2(194)$ = 14,282.05; AIC = 14,400.05; CFI = .844; SRMR = 0.087).

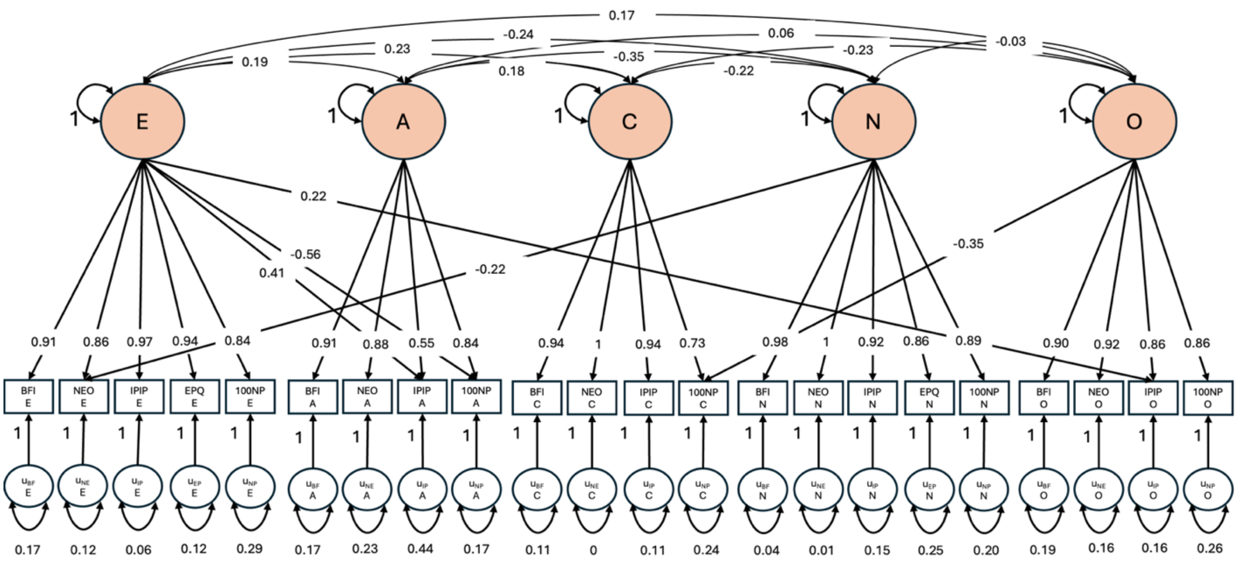


**Supplementary Figure S25. Final factor model of measurement-instrument stratified GWAS of the Big Five, allowing for cross-loadings.**

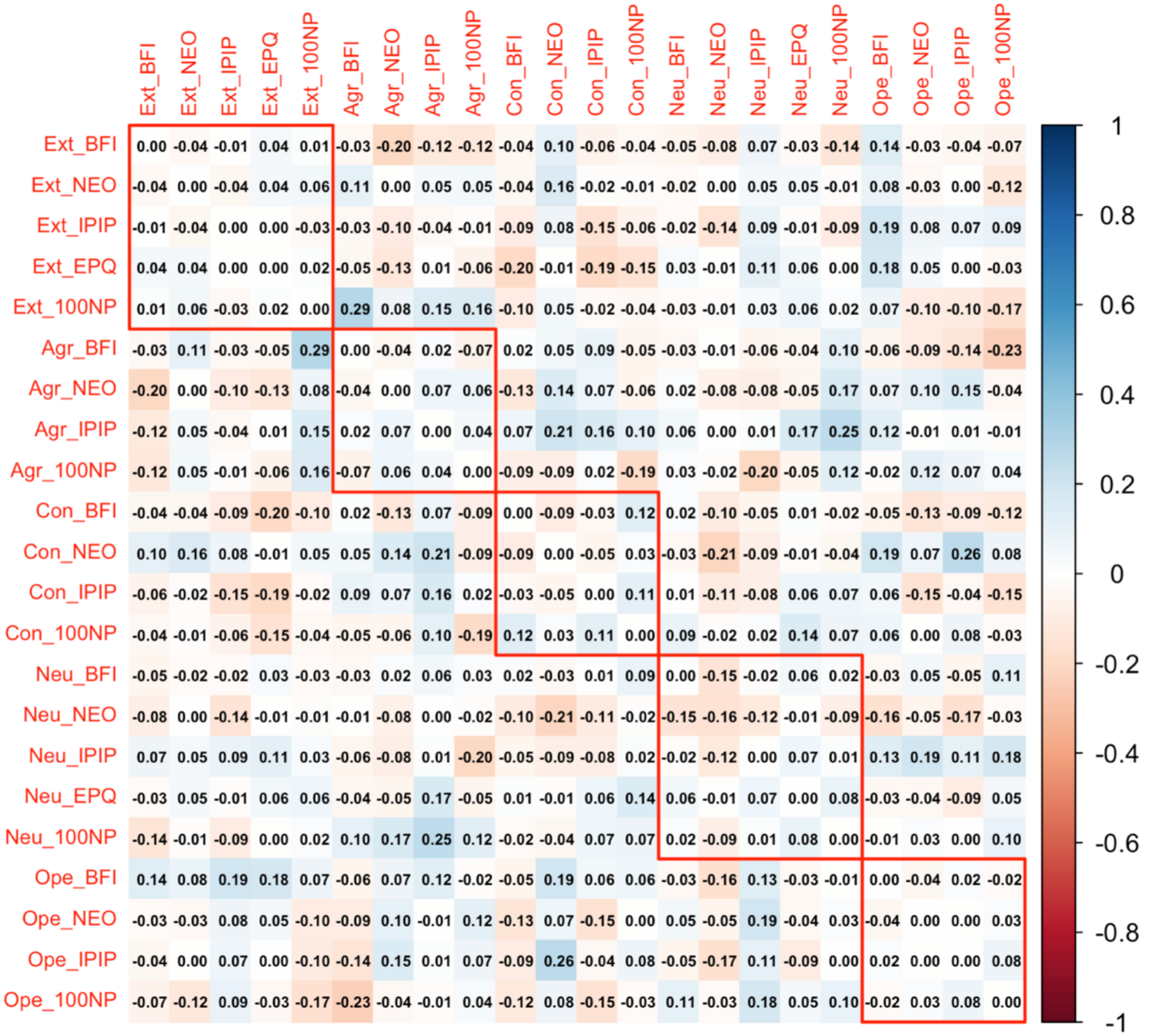


**Supplementary Figure S26. Residual genetic correlation matrix from final factor model after allowing for cross-loadings.**

We also conducted an exploratory factor analysis (EFA) on the LDSC-derived genetic correlation matrix of all 22 measurements (**Supplementary Table S28**). Note that, unlike confirmatory factor models fit within Genomic SEM, such EFAs are not optimized using the LDSC V matrix (which contains information about the magnitude and dependencies of estimation errors in the genetic correlation matrix) and should therefore be viewed as only approximate. We observed good agreement between the EFA cross-loadings and the cross-loadings from our second model – the only exception being IPIP openness loading on the extraversion factor (present in our final model but not the EFA solution).

To illustrate the value of this iterative model fitting procedure, we again consider 100NP agreeableness. In the poor fitting simple structure model (**Supplementary Figure S23**), we observed that 100NP agreeableness had a very low loading on its latent factor (0.33). However, in the final model, in which the 100NP agreeableness is allowed to additionally cross-load on extraversion, its loading on the agreeableness factor is strong (.84). In other words, failing to model the effect of low extraversion on 100NP agreeableness produced a misimpression that 100NP agreeableness does not reflect agreeableness genetics well.

3.5.1 Genome-Wide Association Models with SNP effects

We extended the Genomic SEM analysis to conduct multivariate GWAS using both the confirmatory five factor model with simple structure (depicted in **Supplementary Figure S23**) and the final factor model with cross-loadings (depicted in **Supplementary Figure S25**). These multivariate GWAS differ from the primary GWAS reported in the main text in three important respects. First, these models specify effects of SNPs to simultaneously predict each of the Big Five personality trait factors rather than one trait at a time. Second, in these models, each of the Big Five are modeled as latent variables composed of the common variance across instrument-stratified GWAS meta-analyses (**Supplementary Figure S22**), rather than as observed variables across the full GWAS meta-analytic sample. Omission of cohorts that did not employ the major measurement instruments for which we conducted instrument-stratified GWAS reduces power for these analyses. Third, estimation of effects in the context of multivariate GWAS model allows us to compute an omnibus $Q_{SNP}$ statistic for each SNP in each of the two multivariate GWAS (degrees of freedom = 17: 22 possible SNP-phenotype path regressions minus 5 SNP-factor regressions) converted to a 1df $\chi^2$ statistic. $Q_{SNP}$ is an index of heterogeneity in SNP-phenotype relations beyond what we would expect under the factor model (Grotzinger et al., 2019; Clapp Sullivan et al., 2023). When $Q_{SNP}$ is high, the expectation that the genetic variant operates on the individual GWAS phenotypes exclusively via the factors is violated.

Results from both multivariate GWAS are summarized in **Supplementary Table S29** and Manhattan plots for each Big Five personality trait and for $Q_{SNP}$ from a multivariate GWAS using the final factor model with cross-loadings are displayed in **Supplementary Figure S27**. Results were similar for both models. Both in terms of number of lead SNPs and in terms of the mean association $\chi^2$ (1) compared to $Q_{SNP}$ $\chi^2$ (1), there was overall a great deal more association signal than $Q_{SNP}$ signal, indicating that the SNP effects on the individual measures largely plausibly act via the factors.

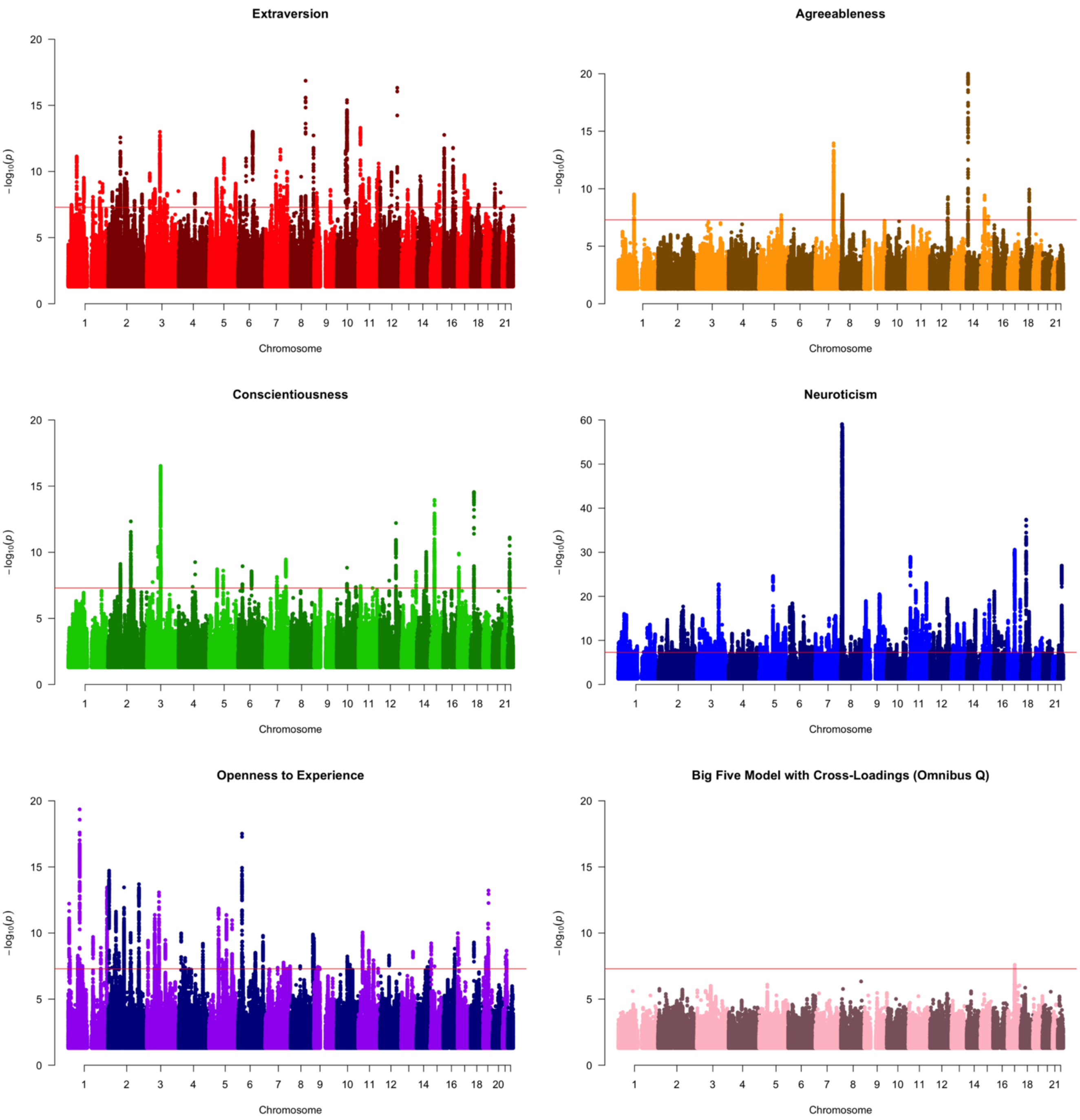


**Supplementary Figure S27. Manhattan plots of for each Big Five personality trait and for $Q_{SNP}$ from measurement-instrument stratified GWAS using the final factor model with cross-loadings.** Genome wide significance is denoted with a red line at p = $5\times10^{-8}$. Note that the Y-axis scale for neuroticism is scaled differently than the other panels.

As depicted in the bottom-right panel of **Supplementary Figure S27**, we identified one genome-wide significant lead SNP for omnibus $Q_{SNP}$ in the final factor with cross-loadings model, rs8712: effects of this SNP were significantly heterogenous across measurement instruments. Using the National Institute of Health's LDlink module (Machiela & Chanock, 2015), we found this SNP to be in strong LD with a lead SNP for neuroticism (rs62062288; $R^2$ = .94) and a lead SNP for extraversion (rs62057151; $R^2$ = .94. These SNPs were also in high LD with one another ($R^2$ = .90), where the trait-increasing allele for extraversion was associated with the trait decreasing allele for neuroticism. The FUMA GWAS catalog (Watanabe et al., 2017) indicated the lead SNP for omnibus $Q_{SNP}$, rs8712, has been found to be associated with a wide range of traits beyond personality, including cognitive performance, lung function, Parkinson's disease, educational attainment, alcohol consumption, snoring, and breast cancer. Thus, this locus appears to be especially pleiotropic, linked to multiple impactful outcomes.

To visualize the heterogeneity of effects of lead SNP rs8712 across instruments, we present plots of that SNP's effects on the individual measures as a function of the corresponding factor loadings (**Supplementary Figure S28, left panel**) and, for comparison purposes, representative hits for each of the Big Five traits, which tend to have much lower omnibus $Q_{SNP}$ values (**Supplementary Figure S29**). Because rs8712 is in LD with genome-wide significant hits for extraversion and neuroticism, we display the estimates specifically for measures loading on those factors representing those traits, and we produce the plots for the corresponding extraversion and neuroticism hits (**Supplementary Figure S28, middle and right panels**). Note that for ease of interpretation, estimates used for these plots were standardized with respect to the genetic variance of each measure, such that a loading of 1.0 corresponds to all of the genetic variance in the corresponding measure accounted for by the factor. Multivariate GWAS estimates for these SNPs are reported in **Supplementary Table 30**. In these figures, regression lines correspond to the model expectations, with the slope corresponding to the estimated SNP effect on the respective factor. The extent of scatter of the points around each line depicts the heterogeneity in effects relative to their expectations (greater scatter corresponds to higher $Q_{SNP}$). These plots can both be used to visualize heterogeneity and identify specific indicators for which SNP betas deviate considerably from expectations. For rs8712, there is considerable scatter around the expectations. Of note is the near zero effect of this SNP on 100NP extraversion, despite that measure's strong loading on extraversion, and similarly, the near zero effect of this SNP NEO neuroticism despite that measure's strong loading on neuroticism. In comparison, there is tight correspondence between SNP effects and their expectations for representative low $Q_{SNP}$ genome-wide significant hits for each of the Big Five.

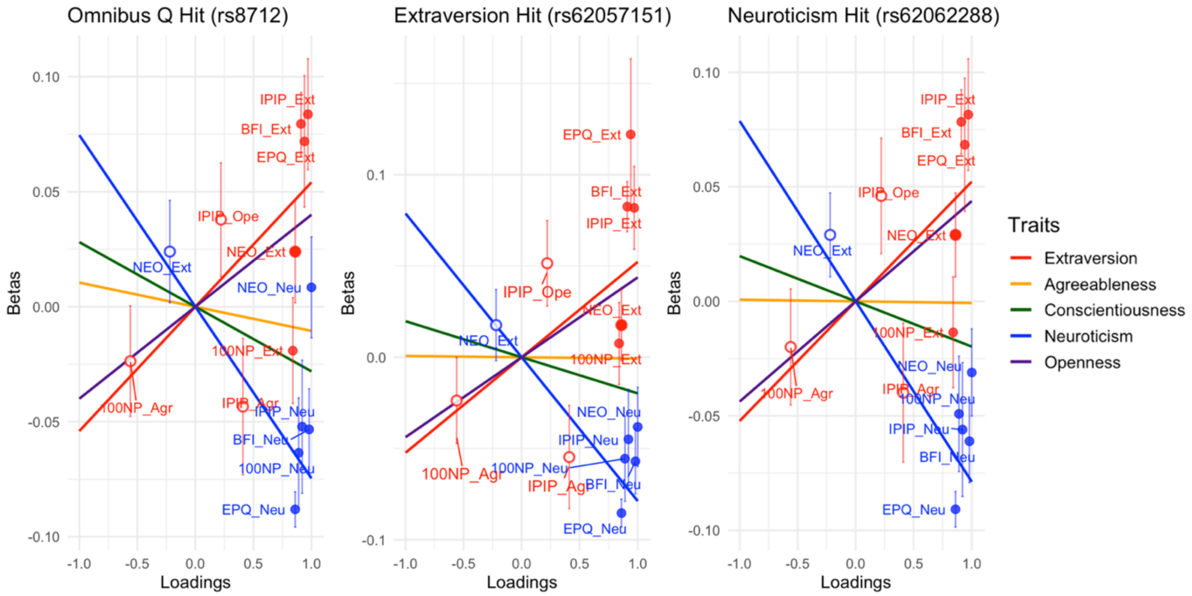


**Supplementary Figure S28. Plots of SNP betas on extraversion and neuroticism measures as a function of corresponding loading for a genome-wide-significant $Q_{SNP}$ that is in high LD with Extraversion and Neuroticism hits (left panel), and corresponding plots for the extraversion and neuroticism hits (middle and right panels).** Both betas and loadings are standardized with respect to the genetic variance of the corresponding measure. Points are colored according to the trait on which the loading corresponds to. Solid points are primary loadings. Unfilled points are cross-loadings (from a measure of another trait). Error bars are standard errors. The superimposed lines have slopes equal to the estimated SNP effect on the respective trait and represent the model-based expectations. The diffuse scatter of the points around their expectations is illustrative of the implausibility of assumption that the genetic variant operates on the individual measures via the factors (i.e. high $Q_{SNP}$).

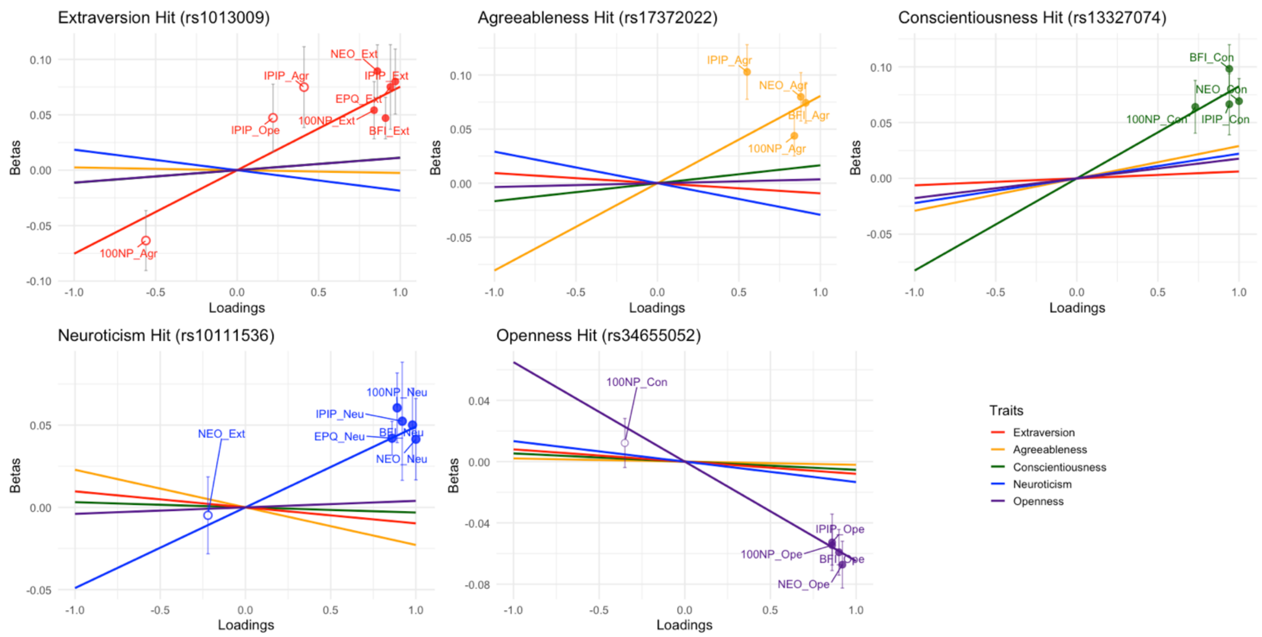


**Supplementary Figure S29. Plots of SNP betas on each individual Big Five measure as a function of that measure's loading for a representative genome-wide significant hit with low $Q_{SNP}$ for each trait.** Both betas and loadings are standardized with respect to the genetic variance of the corresponding measure. Points are colored according to the trait on which the loading corresponds to. Solid points are primary loadings. Unfilled points are cross-loadings (from a measure of another trait). Error bars are standard errors. The superimposed lines have slopes equal to the estimated SNP effect on the respective trait, and represent the model-based expectations. The comparatively steep slope of the line corresponding to factor for which the SNP is a hit, illustrates the specificity of the SNP effect on the factor, and the tight scatter of the points around their expectations is illustrative of the plausibility of assumption that the genetic variant operates on the individual measures via the factors (i.e. low $Q_{SNP}$).

#### 4. Polygenic Prediction of Personality

4.1 PGI Rationale and Cohorts

Polygenic Indices (PGIs) are individual-level summary scores calculated as the sum of many genetic variants' weighted associations with a given phenotype (Burt, 2024). We leveraged the marked increase in power from our discovery GWAS over past studies of personality genomics to estimate PGIs and quantify their predictive accuracy across five independent cohorts: the National Longitudinal Study of Adolescent to Adult Health (Add Health), German Socioeconomic Panel (GSOEP), Health and Retirement Study (HRS), Netherlands Twin Register (NTR), and Twins Early Development Study (TEDS).

Add Health is a large, representative cohort (N ~ 20,000) of young adults assessed through adolescence and early adulthood. Data on personality and outcomes related to juvenile delinquency were assessed in five waves from 1995 – 2018, with the first wave in 1995, the second wave in 1996, the third wave from 2001-2002, the fourth wave from 2008-2009, and the fifth wave from 2016-2018. A subset of participants (N ~ 3,000) provided genetic data during the fourth wave of assessment. For three phenotypes in Add Health (Ever driven drunk, ever arrested, and ever in prison) a large proportion of participants were coded as a "legitimate skip," meaning they were unlikely to have done the behavior based on previous survey responses. We coded skip responses as NA, resulting in a sample of participants for these three traits that had higher prevalences. Sample sizes for each trait including skips are as follows: Ever driven drunk (N = 6,312 (4,153 skips)), Ever arrested (N = 7,648 (6,103 skips)), ever in prison (N = 9,123 (6,418 skips)).

In total, we estimated PGI associations in Add Health participants with European (N = 5,111) and African (N = 1,753) ancestry that provided personality, genetic, and delinquency data.

The German Socioeconomic Panel (GSOEP) is a representative cohort of Germans that have been assessed annually since the 1980s. GSOEP participants provided personality information every four years from 2005-2021 using an adapted German short version of the Big Five Inventory (Hahn et al., 2012). A subset of these participants also participated in the GSOEP Innovation Cohort, where they contributed genotype data (Koellinger et al., 2023). We estimated PGI associations among the N = 2,066 participants who contributed both personality and genomic data that passed strict quality control procedures as described by Koellinger and colleagues.

Finally, the HRS, NTR, and TEDS cohorts are described in the **Contributing Cohorts** section below. These cohorts also contributed to the GWAS discovery analyses. To ensure that there was no sample overlap between discovery and prediction target, we re-estimated genome-wide associations for each of these cohorts with the target prediction cohort removed from the discovery sample, and we used these independent discovery summary statistics for PGI prediction in each cohort.

Testing PGI associations across five different cohorts allowed us to quantify the robustness of these results across age, instrument, geography, and ancestry. With respect to age, Add Health, NTR, and TEDS ascertained younger participants (mean age < 26), HRS ascertained older participants (ages 51+), and GSOEP ascertained a representative lifespan cohort. With respect to instrument, Add Health measured personality with the 20-item IPIP inventory, HRS measured personality with the 31-item MIDI inventory, NTR used the 60-item NEO-FFI 3, TEDS used the 30-item BFI-30. With respect to nationality, Add Health and HRS sampled US residents, GSOEP ascertained German residents, NTR ascertained Dutch residents, and TEDS ascertained English residents. All five cohorts ascertained participants of European-like ancestry, and Add Health and HRS additionally ascertained participants of African-like ancestry, which allowed us to benchmark cis-ancestry and trans-ancestry prediction (sample sizes of the AFR discovery GWAS were not large enough to use AFR discovery data to estimate PGI weights).

4.2 Method and Population-Level PGI Model

We derived PGI weights from EUR GWAS summary data using SBayesR (Lloyd-Jones et al., 2019), which has been benchmarked as the most effective method for highly polygenic traits that likely follow a mixed distribution of effect sizes across alleles. SBayesR is a Bayesian method that estimates PGI weights by sorting SNPs into a distribution of effect sizes. For each of the Big Five, we estimated PGI weights with HapMap3 SNPs sorted into a distribution (pi) of 95%, 2%, 2%, and 1%, with effect sizes (gamma) scaled at .00 (no effect), .01, .10, and 1.00. The LD reference for these models was calculated for HapMap3 SNPs using 10,000 unrelated people with European Ancestry in the UK Biobank (Lloyd-Jones et al., 2019). We excluded the MHC region from these analyses. SBayesR models can fail to converge when estimating weights from summary data with varying sample sizes across SNPs; this was the case for openness to experience, extraversion, and neuroticism. To ensure model convergence, we pruned SNPs outside 1SD of the max sample size for those traits. In total, we created 5 polygenic index weights, one for each of the Big Five traits, using 712,425 (neuroticism) to 1,131,471(conscientiousness) SNPs per trait.

Then, we applied these PGI weights to estimate a score for each participant, using PLINK2 (Chang et al., 2015). Each cohort ascertained personality at multiple occasions; to maximize trait influences and minimize state influences on personality scores for each participant, we estimated participant scores as the mean across assessment waves. We then regressed participants' standardized personality trait score (*y)* on their standardized polygenic score, controlling for their mean age across waves they contributed personality data, their biological sex, age, age$^2$, 10 ancestral principal components (PCs), and any other necessary covariates (e.g., genotyping chip) as follows:

$$y = b_0 + \ b_1 PGI + \sum b_c x_c + e \ , \qquad \text{(Eq. 10)}$$

where $b_0$ is a regression intercept, $b_1$ is the standardized effect of the PGI, $b_c$ is the effect of covariate $x_c$ on the personality trait for covariates 1 through C, and *e* is a residual.

We refer to this as *population-level* PGI prediction, as it predicts variation in personality from variation in genotype in the population. The focal output of this model is the $b_1$ parameter, which represents the standardized association between the PGI (genotype) and the Big Five trait score (phenotype) after controlling for covariates. In the three cohorts of unrelated participants (Add Health, GSOEP, and HRS), standard OLS standard errors were obtained. In the two cohorts with related participants (NTR and TEDS), we estimated this model with cluster-robust standard errors, with family ID serving as the cluster, to account for non-independence of data across family members. Results of these analyses are described in the **Main Text.**

**5. Associations with socially relevant behaviors and important life outcomes**

5.1 LDSC Correlations with External Phenotypes

To quantify the widespread relevance of personality genetics, we identified 61 external phenotypes that had previously been GWASed in participants with recent European-like ancestry with publicly available summary statistics. We consider these phenotypes in terms of nine clusters: Diet, education and employment, general health, participation, personality, physical health, psychopathology, reproductive health, and substance use. Full information on each external phenotype, including reference and sample size, is presented in **Supplementary Table S31**.

For each phenotype, we estimated bivariate LDSC genetic correlations with the EUR Big Five summary statistics, as described in section 3.3 above. Results from these tests are described in the **Main Text** and **Supplementary Table S32**.

For 13 of these 65 phenotypes, we were also able to estimate bivariate genetic correlations among participants with recent African-like ancestry, with both Big Five and external phenotype summary statistics estimated using AFR data, using LD scores estimated from the 1000 genomes 3v5 AFR data. We visualize these associations in **Supplementary Figure S30** below.

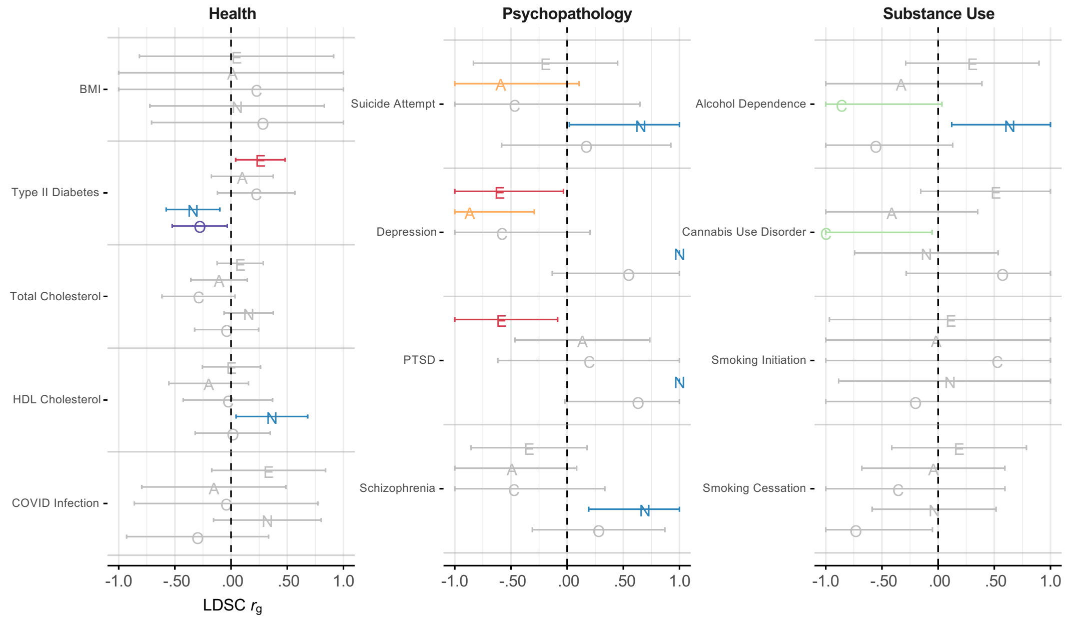


**Supplementary Figure S30. Genetic correlations between personality and health, psychopathology, and substance use among participants of African-like ancestry.** Note. E = Extraversion. A = Agreeableness. C = Conscientiousness. N = Neuroticism. O = Openness to Experience. Error bars depict 95% confidence intervals. Correlations not significant at $p < .05$ depicted in gray. Correlations estimated using Linkage Disequilibrium Score Regression.

Overall, results of genetic correlations among participants with recent African-like ancestry had substantially wider confidence intervals than in EUR analyses, which is attributable to smaller GWAS sample sizes in these analyses and lower heritability of phenotypes in AFR participants (see section 3.1 above). When considering the 95% confidence intervals of these analyses alongside those from EUR analyses, we observed strong compatibility with results across ancestry. In both sets of results, neuroticism was positively correlated with liability for all psychopathology outcomes at $p < .05$, extraversion was negatively correlated with PTSD and depression liability, and conscientiousness was negatively correlated with alcohol dependence and cannabis use disorder liability. We observed one phenotype where significant results differed in sign across ancestries: in EUR data, Type-II Diabetes was positively associated with neuroticism, but in AFR data, it was negatively associated with neuroticism. This difference was trait-specific, as Type-II Diabetes had a negative genetic correlation with openness to experience and a positive genetic correlation with extraversion in both groups.

5.2 Polygenic Prediction of External Outcomes in Add Health

When possible, it is ideal to index genetic sharing between two phenotypes using bivariate LDSC, as estimates of genetic correlations using this method are unbiased by measurement error. However, this requires GWAS summary data, which may not be available for a certain phenotype. Add Health in particular has measured external outcomes spanning delinquency, interviewer-rated behavior, and contact with the criminal justice system, which are germane to personality yet have not been GWASed. Thus, we supplemented bivariate LDSC associations with PGI analyses in the Add Health dataset, among EUR (max N = 5,111) and AFR participants (max N = 1,753).

Participants in Add Health were interviewed in-person by a trained interviewer, who then was asked to rate the participant they interviewed. Specifically, they were asked to rate the interviewees' physical attractiveness and the attractiveness of their personality on a scale from 1 (very unattractive) to 5 (very attractive), as well as how well-groomed the interviewee was. Interviewers also rated how candid the respondent seemed in the interview from 1 (very candid) to 4 (not candid), indicated whether the respondent ever seemed bored or embarrassed during the interview, and recorded the number of times the interviewee interrupted the interviewer.

We also incorporated 8 measures of juvenile delinquency measured in Add Health. A subset of at-risk Add Health participants were asked whether they had ever been expelled from school, received an out-of-school suspension, driven while drunk, run away from home, been initiated into a named gang, or had ever been arrested or taken into custody by police. Participants were also asked how often they had lied to their parents or guardians in the past 12 months.

Finally, Add Health participants provided data on contact with the prison system: whether they had ever served time in a jail, prison, juvenile detention center, or other correctional facility, and whether their biological mother or biological father had ever spent time in jail or prison.

To predict each variable from personality PGI, we estimated an extension of Equation 10. In this model, the right-hand side of the model was identical, and, on the left-hand-side, the predictor was each external outcome rather than personality PGI. For continuous outcomes, we report the beta coefficient of a model in which both the PGI and outcome were standardized. For binary outcomes, we report the beta coefficient of a logistic regression model. Certain variables were measured at multiple occasions: how often participants lied to their parents and interviewer-rated grooming, candidness, physical attractiveness, personality attractiveness, and number of interruptions. In these cases, the outcome variable was calculated as the mean score on this variable across all waves that a participant provided data. Other variables represent cumulative outcomes (e.g., "*Have you ever…*"). For these outcomes, we predicted participant scores at their most recent measurement occasion.

We present results of prediction in participants of European-like ancestry in the **Main Text**. Below (**Supplementary Figure S31),** we present results of prediction in participants of African-like ancestry. In general, analyses had much lower power in AFR than EUR analyses, due to the smaller prediction sample and discrepant ancestry between the EUR GWAS discovery sample and the AFR prediction target sample. As a result, no associations between the personality PGIs and outcomes in Add Health were significant. All point estimates of PGI associations in the Add Health dataset can be found in **Supplementary Table S33.**

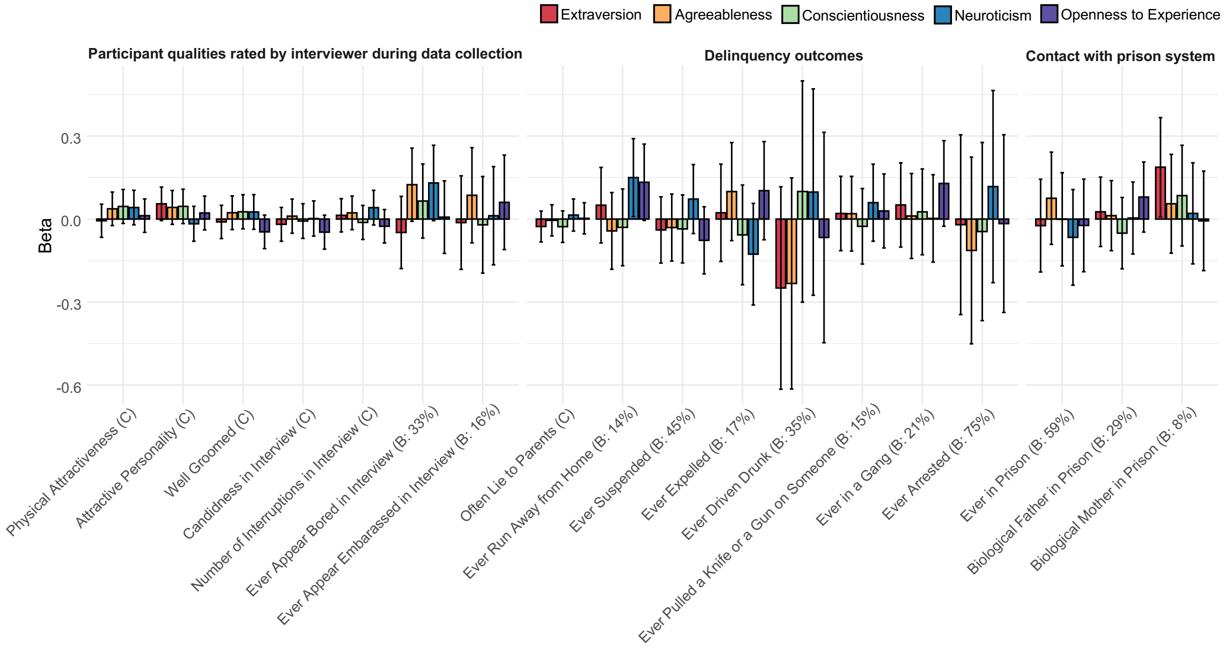


**Supplementary Figure S31. Prediction of outcomes in the Add Health dataset from personality trait polygenic index among participants with recent African-like ancestry (N = 1,753)**. Note: Effect sizes with continuous outcomes (labeled with a C) are standardized, whereas associations with binary outcomes (labeled with a B, with percentage affirmative answers) are in terms of logistic betas. Error bars depict 95% confidence intervals. No associations are significant at $p < .01$.

5.3 Residential Analyses

People make choices regarding their socio-demographic residential environment throughout their lives. These choices involve both economic trade-offs (i.e., *"Where can I afford to live?")* and preferences (i.e., *"With which geo-social environment do I have an affinity?"*). Such decisions have implications for both health and wealth, as the chosen environment affects exposure to environmental risks and social/educational opportunities. Personality may influence both the ability to afford specific socio-demographic environments and one's affinity towards them.

To test associations between personality genomics and residential environment, we analyzed data from 419,261 UK Biobank participants of European ancestry, born in the UK, and aged 39-70 at the time of enrollment. For each participant, we computed PGIs for all five personality traits (see section 4.2 above for PGI estimation). For the Neuroticism PGI, we held out participants from the UK Biobank, as well as Generation Scotland and the Lothian Birth Cohorts 1921 and 1936, to conservatively account for an unknown degree of participant overlap among these cohorts.

For our dependent variables, we obtained the approximate Lower Layer Super Output Areas (LSOA) of participants' current addresses (smaller geographic regions containing approximately 400-1,200 households) (https://www.ons.gov.uk/methodology/geography/geographicalproducts/areaclassifications). Based on these LSOAs, we derived local urbanicity and UK Census-based residential area classifications. These classifications are categorical groupings of British neighborhoods derived by clustering based on 60 census variables, covering demographics, household composition, housing nature, socioeconomic composition, and employment. The resulting clusters include descriptive labels, such as “Comfortable Suburbia” or “Cosmopolitan Student Neighborhoods.” Where needed, we redrafted the category labels to enhance transparency for those unfamiliar with UK demographics.

For our first analysis, we regressed area classification on personality PGI, controlling for five principal components, age, sex, genotyping array, and including Middle Layer Super Output Areas (MSOAs) at birth as a fixed effect. MSOAs are geographic regions of approximately 2,000-6,000 households with 5,000-15,000 individuals; adding them as control variables therefore tests whether personality PGI is related to residential affordability and preferences later in life, comparing individuals to their childhood neighbors to ensure the association reflects affordances and preferences from birth (i.e., not confounded by childhood socioeconomic status or environment). We applied a false-discovery rate correction to $p$-values of these tests and considered adjusted $p$-values $< 0.01$ as statistically significant.

As we show in **Supplementary Figure S32,** we found widespread associations between openness to experience, neuroticism, and conscientiousness PGIs and residential area classification in late adulthood. Openness to experience PGI was positively associated with residing in “Cosmopolitan Student Neighborhoods,” “Inner City Cosmopolitan” areas, and “Highly Qualified Professionals’” areas, and negatively associated with various constrained and suburban residential areas. Neuroticism PGIs displayed mostly inverse associations relative to openness to experience, with low neuroticism (but not high openness to experience) associated with living in areas classified as “Prosperous Countryside Life” and “Affluent Communities.” Conscientiousness PGIs were negatively related to cosmopolitan neighborhoods and positively related to suburban living.

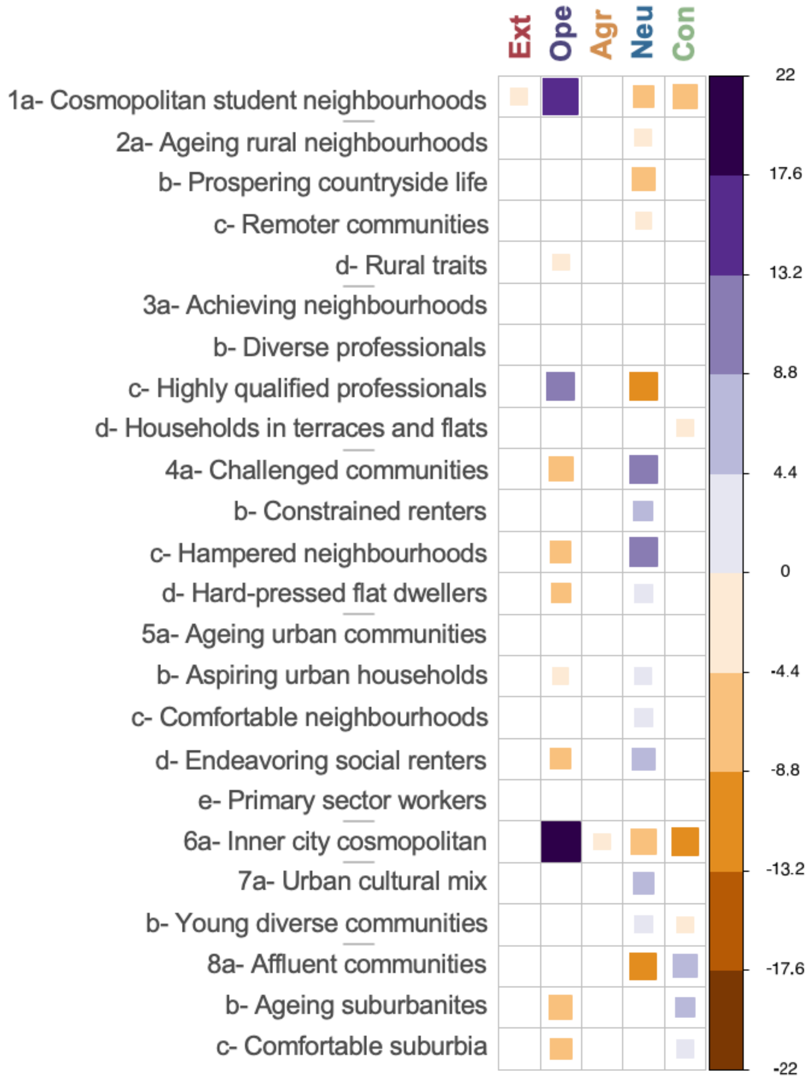


**Supplementary Figure S32. Residential preferences PGI heatmap.** Heatmap based on the Z-statistics (right-hand bar) of the association between Big Five PGI and 22 area classifications. Associations are estimated controlling for 5 principal components, sex, age, genotyping array, and a fixed effect for MSOA region at birth.

We then repeated Analysis 1 within 20,517 sibling pairs, where the power for within-sibling analysis of a binary outcomes (e.g., residence in a certain area) was determined solely by discordant pairs. For individual area classifications, we identified between 100 and 3,602 discordant pairs (median 1,280 pairs), meaning these analyses were underpowered. We then filtered the results from Analysis 1 based on effect sign concordance in the within-sibling analysis and a within-sibling Z-statistic above 1. This filtering provides stricter control for childhood SES and other confounders, as families share potential socioeconomic confounders more intimately than individuals born in the same MSOA-level neighborhood.

When applying this filtering, most associations between neuroticism PGIs and area classifications were pruned (See **Figure 4 panel E in the Main Text**), suggesting that they may be the result of environmental confounding, but other associations persisted, especially for openness to experience and conscientiousness.

Finally, we repeated Analysis 1, now including median home prices at the MSOA level in 1995, 2000, 2005, and 2010 to control for the effects of personality on an individual's ability to afford specific socio-demographic neighborhoods. The remaining effect plausibly reflects socio-cultural preference, rather than affordability. Controlling for local housing costs in the 1995-2010 timeframe attenuated the association between are-classification and personality PGI relative to analysis 1, and this attenuation was not specific to any one personality domain. We present this correlation heatmap in **Supplementary Figure S33**.

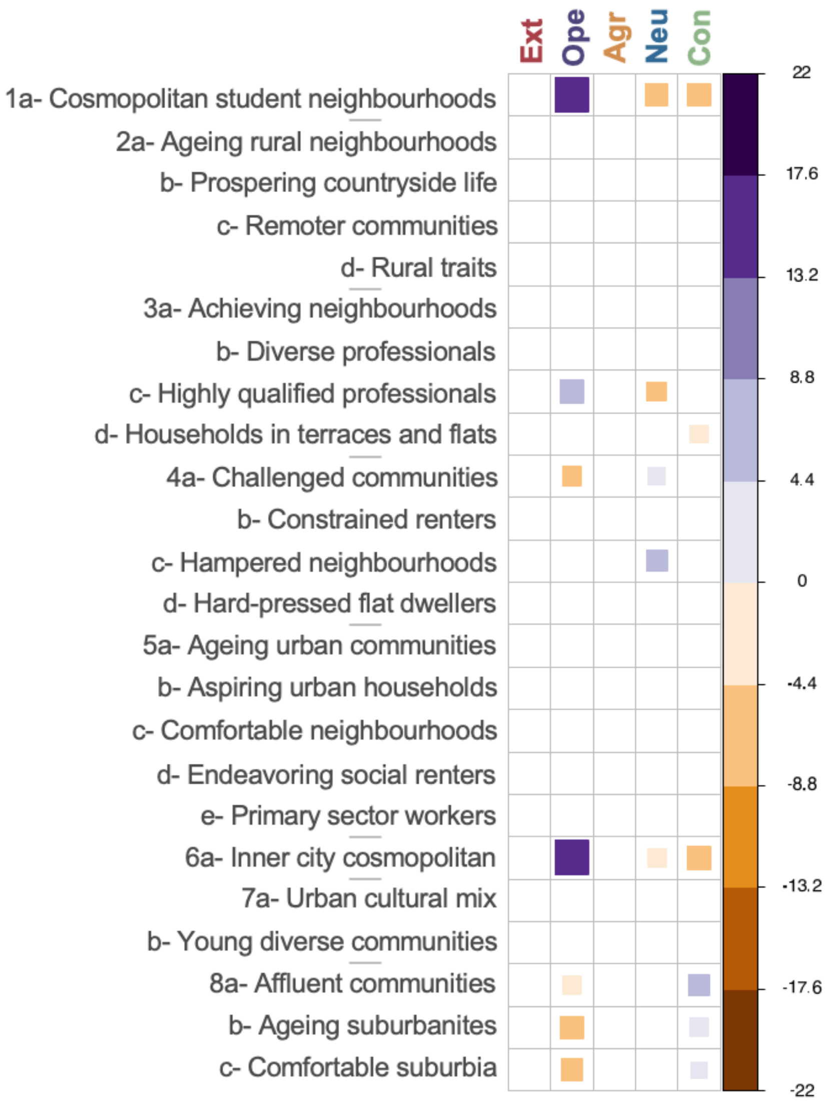


**Supplementary Figure S33. Residential preferences PGI heatmap controlling for median home prices.** Heatmap based on the Z-statistics (right-hand bar) of the association between Big Five PGI and 22 area classifications. Associations are estimated controlling for 5 principal components, sex, age, genotyping array, and a fixed effect for MSOA region at birth, plus median home prices in 1995, 2000, 2005, and 2010.

5.3.1 Urban/Rural Migration

We also explored the extent to which personality PGIs predicted movement out of rural and urban areas, or continued residence in them, among 265,410 UK Biobank participants. Location was classified as urban or rural using the 2011 Rural-Urban Classification for Local Enterprise Partnership Areas based on Census Output Areas (https://www.gov.uk/government/statistical-data-sets/local-enterprise-partnerships-leps-rural-urban-gis-shapefiles). We categorized participants into four bins according to whether they were born in a rural or urban area and whether they currently reside in a rural or urban area, and we predicted category membership from population-level EUR personality PGI, controlling for age and ancestral principal components.

As we visualize in **Supplementary Figure S34**, PGIs for each of the Big Five predicted residential movement across the lifespan. Similar to the residential preference PGI analyses above, participants who inherited genetic profiles associated with higher openness to experience were more likely to move to urban areas from rural birthplaces, and genetic profiles associated with conscientiousness and agreeableness predicted movement to rural areas from urban birthplaces. Complementing above residential preference results, participants who inherited genetic profiles associated with higher conscientiousness were also more likely to remain in rural areas (current residence and place of birth in rural areas), whereas openness to experience PGI was associated with lower likelihood of remaining in rural areas but not associated with remaining in urban areas. This helps further contextualize the genetics of openness to experience as especially related to residential niche-picking via leaving rural areas. These associations were highly similar, albeit with wider confidence intervals, across males and females.

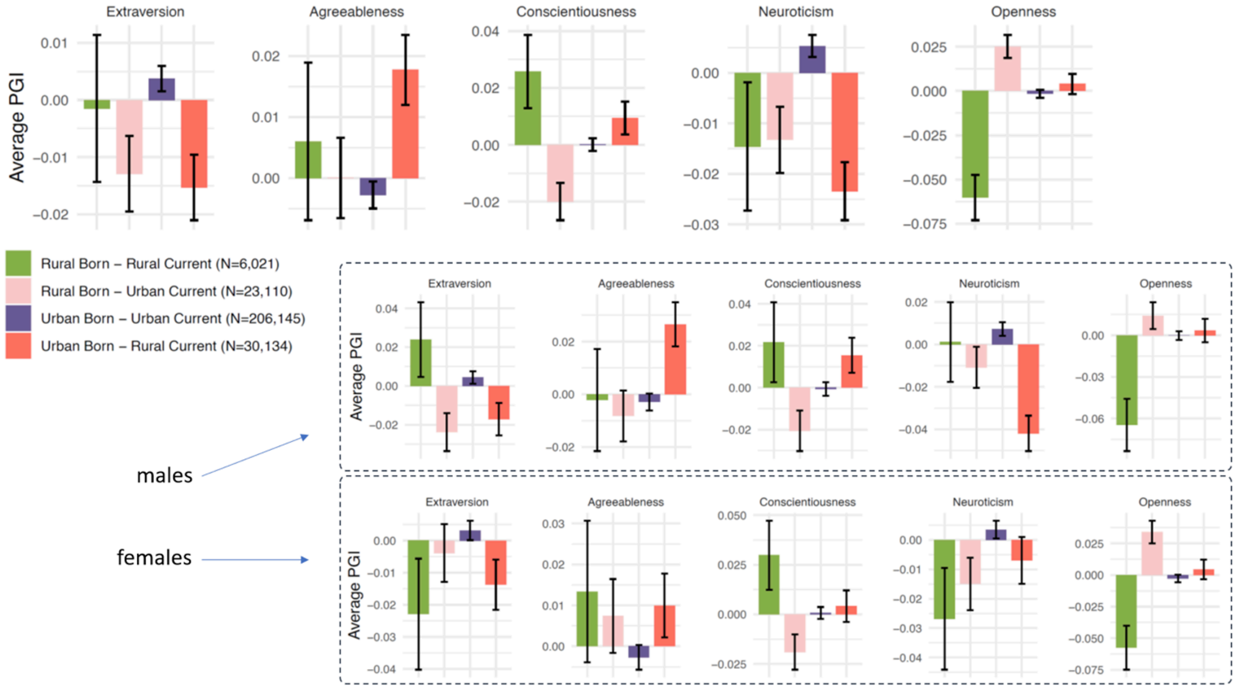


**Supplementary Figure S34. PGI associations with moving or remaining in urban and rural areas in the UK Biobank.** Error bars represent 95% Confidence Intervals.

5.4 Assortative Mating

Assortative mating (AM), the tendency of parents to resemble each other on a given trait, is a widespread human phenomenon (Horwitz et al., 2023). For personality traits, however, there is evidence for very little assortative mating: parents tend not to resemble each other in their phenotypic personality. In **Supplementary Figure S35** below, we visualize this phenomenon in the deCODE cohort (N = 5,317 mate-pairs). In this cohort, AM was greater for openness to experience than other traits, replicating past research (Horwitz et al., 2023; McCrae et al., 2008).

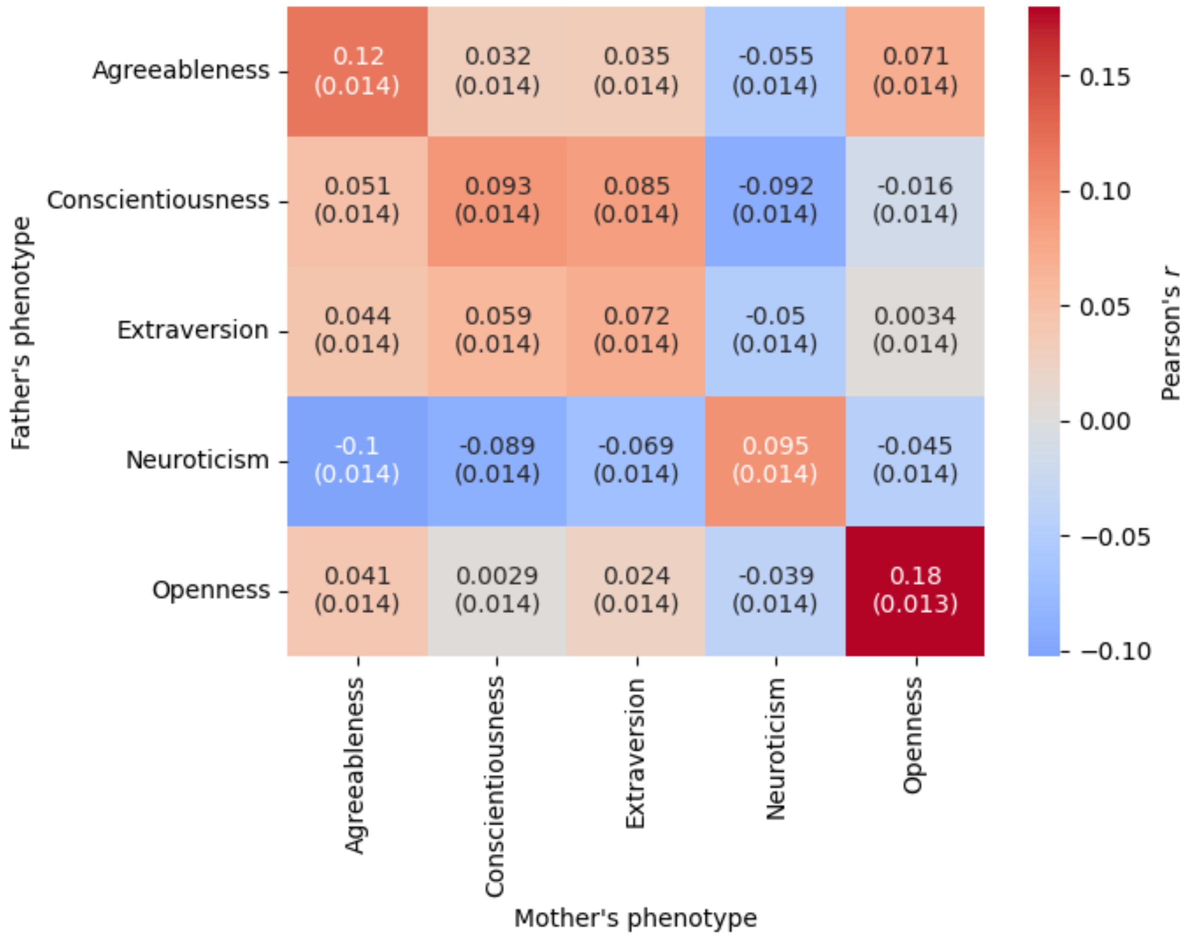


**Supplementary Figure S35. Phenotypic assortative mating correlations for the Big Five in the deCODE cohort.** Depicted are phenotypic cross-mate correlations for 5,317 unique mate-pairs from the Icelandic population. Mates were defined as individuals known to have children together in the deCODE genealogical database. Each square shows the correlation coefficient with standard error in parenthesis.

AM can also confound genetic inference. It induces genetic correlations across the genome that affect estimates of heritability and genetic correlations with other traits (Border et al., 2021). Recent research has quantified AM using molecular genetic data, which tests the extent to which AM has existed over generational time (Torvik et al., 2022).

We developed a novel model to quantify historical AM using genomic data. The closest antecedent to the latent PGI model proposed here is Tucker-Drob (2017), which outlines error correction in PGI analysis as a latent variable model. In this model, two PGI estimates for the same trait are derived from disjoint discovery GWAS (with no sample overlap and approximately the same sample size). Here, we expand this framework to include two PGI estimates for a single trait in a pair of individuals, who may be siblings, cousins, spouses, etc. Utilizing pairs simplifies identification, which is advantageous. Notably, using PGIs, it is unnecessary to measure the trait in the target cohort, a feature that is both powerful and essential for datasets that have not measured a focal phenotype, such as the Big Five personality traits. We depict the structural equation model path diagram for this method in **Supplementary Figure S36**.

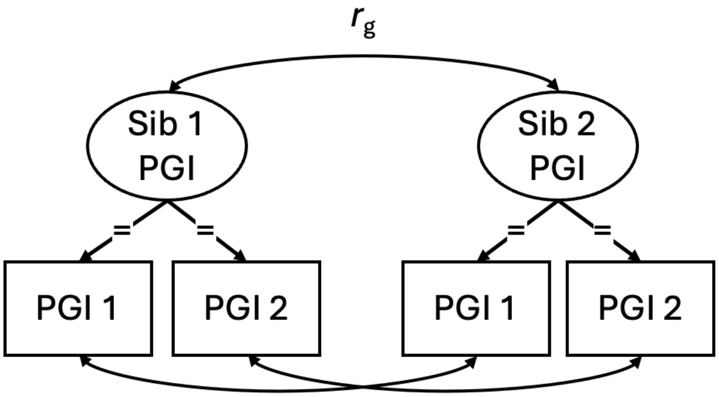


**Supplementary Figure S36. Latent Assortative Mating Model**. In the path diagram of the latent PGI model, the two PGIs estimated from split-half discovery GWAS data are depicted as PGI 1 and PGI 2. The factor loadings are constrained, denoted by an = sign, while the within-PGI cross-sibling PGI residual associations are allowed to correlate. Factor variances are constrained to 1 to identify the model.

The expected genetic correlation ($r_g$) for siblings is 0.5, and for cousins, it is 0.125. Under assortative mating equilibrium (i.e., when assortative mating has been consistent over multiple generations, resulting in stable genetic correlations), these parameters increase as a function of the genetic correlation between parents ($r_{g\ parent}$)where $r_{g\ parent}$ is the genetic correlation between parents for a given trait due to assortative mating. The functional relationships are as follows:

$$r_{g\ sib} = \frac{(1 + r_{g\ parent})}{2} \qquad \text{(Eq. 11a)}$$

and

$$r_{g\ cousins} = \left(\frac{(1 + r_{g\ parent})}{2}\right)^3. \qquad \text{(Eq. 11b)}$$

And we can express the inverse relation as:

$$r_{g\,parent} = 2 * (r_{g\,sib}) - 1 \qquad \text{(Eq. 12a)}$$

and

$$r_{g\,parent} = 2 * \left(r_{g\,cousins}^{\frac{1}{3}}\right) - 1. \qquad \text{(Eq. 12b)}$$

Based on cousin, sibling and spousal relations we generate 3 estimates of $r_{g\,parent}$. Estimating $r_{g\,parent}$ based on sibs, cousins and spouses gives us three independent estimates which guards against bias in either of the independent estimates.

To test for historical AM in personality, we used data from participants of European-like ancestry in the UK Biobank and deCODE cohorts. In the UK Biobank, we identified 67,944 spousal pairs, 21,361 sibling pairs and 66,745 cousin pairs based on kinship information. Given the UK Biobank age range at recruitment (39-70 years), we are confident in assuming that third-degree relatives based on kinship are cousins, as opposed to grandparent-grandchild relationships. We used IBD0 to distinguish between sibling and parent-offspring pairs. In deCODE, we identified 59,286 spousal pairs, 31,190 cousin pairs, and 42,882 sibling pairs.

We computed two split-half PGIs for each of the Big Five traits. To do this, we split the GWAS discovery samples into two non-overlapping groups matched as closely as possible on demographics to ensure equal representation in groups like country, questionnaire, and sample size (we note that our comparisons in section 3.3 above confirm general similarity in genetic architecture even when groups differ on these variables). Full information on which cohorts were included in which split-halves is provided in **Supplementary Table S34.** Then, we re-estimated meta-analytic genome-wide associations in each split half (as in section 1). Bivariate LDSC comparisons for each Big Five trait between each of the halves confirmed a high degree of similarity across halves (mean $r_g = .94$) and a low degree of sample overlap between them (mean cross-trait intercept = .0095). We estimated PGI weights from each split half and applied them to UK Biobank participants (following the method in section 4).

Finally, we applied the model described in **Supplementary Figure S36** to these data. By estimating two PGIs for each participant and then estimating a latent variable from their common variance, we obtain reliability-disattenuated estimates of genetic variance (Tucker-Drob, 2017). By scaling these estimates to 1 and then estimating their covariance across pairs of relatives, we estimate the genetic correlation between pairs of relatives for each Big Five trait. For spouses, the obtained quantities are direct estimates of $r_{g\,parent}$. For siblings and cousins, we compare these estimates to their expected genetic correlation based on their relatedness using Equation 12 to obtain estimates of $r_{g\,parent}$. We present genetic correlations for each combination of trait, relationship, and cohort in **Supplementary Table S35.** To obtain a single pooled estimate from these data, we estimated an inverse-variance weighted meta-analysis across all relationships and both cohort for each trait, using formulas 5a and 5b, above.

##### 5.5 Mendelian Randomization Analyses

In classical applications of *Mendelian Randomization* (MR), genetic variants with strong effects on a focal exposure are used as instrumental variables to estimate the causal effect of that exposure on a hypothesized outcome (Sanderson et al., 2022). Each individual randomly inherits their autosomal genetic material in equal halves from each parent. Therefore, conditional on appropriate corrections for ancestry, exposure to the genotype approximates random assignment in a natural experiment. An illustrative example of an application of MR is with respect to variants in the nicotinic acetylcholine receptor *CHRNA5–CHRNA3–CHRNB4* gene cluster (as represented by the SNP in **Supplementary Figure S37**), which are mechanistically associated with biological response to nicotine and therefore with variation in cigarettes smoked per day (Exposure) among smokers (Thorgeirsson et al., 2008). MR using these variants have been used to estimate the effect of smoking quantity during pregnancy on offspring birthweight among smokers ($\hat{\gamma}$), deconfounded from unmeasured third variables (e.g. socioeconomic variables) that may affect both smoking quantity and BMI (Lassi et al., 2016). The key identifying assumption of this approach is that the variants are only associated with offspring birthweight via their effects on smoking (and not via direct effects on offspring birthweight or associations with third variable confounds that affect both smoking and offspring birth weight). Supporting these assumptions, the variants are unrelated to offspring birthweight among women who do not smoke during pregnancy but are nevertheless associated with offspring birthweight (Outcome) among women who do smoke during pregnancy (Lassi et al., 2016).

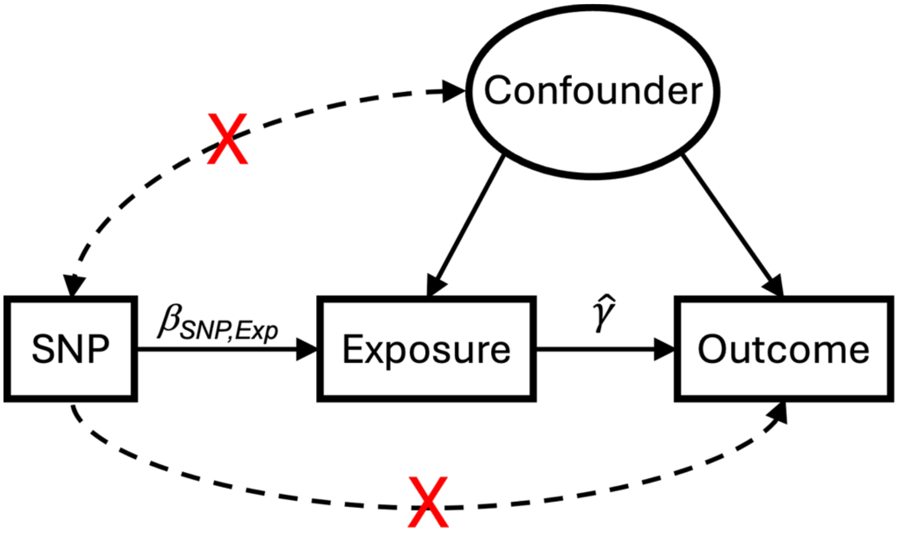


**Supplementary Figure S37. Causal diagram assumed by traditional Mendelian Randomization (MR) approaches.** The dashed lines with the red X represent paths assumed to be absent by traditional MR. The three approaches to MR that were implemented here use different approaches to estimating causal effects of the exposure on the outcome using multiple independent SNPs, such that they are robust to violations of these standard assumptions by a subset of SNPs.

Put generally, valid application of MR requires that three core assumptions be satisfied (Burgess et al., 2023): First, the SNP that serves as the instrumental variable must be relevant to the exposure (i.e., $\beta_{\text{SNP,Exposure}}$ in **Supplementary Figure S37** must be nonzero). Second the variant must be independent of horizontal pleiotropy indicative of confounding effects (the path linking the confounder and the SNP must be absent, meaning there is no correlated pleiotropy). Third, the instrument must be indicative of horizontal pleiotropy that is uncorrelated with confounding effects, meaning it must only affect the outcome through the exposure (the path from SNP to the Outcome must be absent). Given these assumptions, the following equality holds:

$$\beta_{\text{SNP,Outcome}} = \hat{\gamma} \times \beta_{\text{SNP,Exposure}} \text{ , (Eq. 13a)}$$

where $\beta_{\text{SNP,Exposure}}$ is the effect of the SNP of the exposure and $\beta_{\text{SNP,Outcome}}$ is the total effect of the SNP on the outcome (not depicted in the figure). Rearranging this equation provides a simple and intuitive estimator of the causal effect:

$$\hat{\gamma} = \frac{\beta_{\text{SNP,Outcome}}}{\beta_{\text{SNP,Exposure}}} . \quad \text{(Eq. 13b)}$$

Ideally, MR tests are conducted using SNPs with strong effects on an outcome that operate through well-characterized biological mechanisms. For complex and highly polygenic traits like personality, where individual SNP effects are small (i.e., assumption 1 only moderately holds) and biological mechanisms are not well-explicated (i.e., assumptions 2 and 3 are untested), researchers have advocated using advanced robust estimators to test MR (Burgess et al., 2023). Such tests, three of which we implement here and describe below, aggregate inference by drawing on multiple independent SNPs.

5.5.1 MR Method

We conducted tests of two-sample bidirectional mendelian randomization (MR) to estimate the potential causal effects of Big Five personality traits on a set of biobehavioral phenotypes and of those biobehavioral phenotypes on Big Five traits. Based on results of genetic correlation tests (**Supplementary Table S32**), and causal reasoning advanced in the broader personality literature, we selected 13 outcomes to bring forward for MR tests (**Supplementary Table S31**): Age at first sexual intercourse (Andrae et al., 2024), Body Mass Index (BMI) (Arumäe et al., 2021), COVID infection (Peters et al., 2023), drinks per week (Koning et al., 2011), educational attainment (Lüdtke et al., 2011), the multivariate healthy aging/longevity factor (Willroth et al., 2025), smoking initiation (Munafo et al., 2007), sports club participation and frequency of sports participation (Caille et al., 2024), spells in hospital (Atherton et al., 2024), and three measures of research study participation in UK Biobank. These 13 outcomes that we take forward for MR evaluation are well suited for performing tests of presence of causation and direction of causation. Moreover, by applying MR tests to a subset of theoretically-relevant outcomes, rather than all potential outcomes, we substantially reduce the burden of multiple testing.

We excluded psychiatric conditions from our MR testing, as the genetic correlations between personality and psychopathology were in some cases so high that MR methods would not be able to distinguish the direction of causation (c.f., Gupta et al., 2024). For example, the very high genetic correlation between neuroticism and internalizing disorders ($r_g$ = .73) could reflect a confounding structure where patterns of horizontal pleiotropy are present to a degree that the entire causal structure becomes difficult to disentangle. Indeed, associations between mental health diagnoses and personality traits may reflect lifelong co-developmental processes

(Durbin & Hicks, 2014; Ormel et al., 2013; Hopwood et al., 2022) with a wide-ranging set of intermediate confounding mechanisms, obviating simple directional examination of causality. To test this confounding structure, one could consider further multivariable MR analysis or multivariable Steiger filtering; this psychopathology-specific follow up analysis is beyond the scope of the primary GWAS, as its credible execution requires in depth case-by-case consideration.

We also selected three outcomes as negative controls: birthweight, number of sisters, and number of brothers. These outcomes cannot be caused by an individual's personality, as they develop before birth or very early in life and there are no plausible mechanisms by which personality causes them. A (false) positive causal effect on any of these traits would serve as an (imperfect) indicator of the presence and potential magnitude of biases. For example, the number of brothers/sisters one has could be influenced by parental personality, and so any false positive signal could reflect indirect genetic effects, intergenerational confounding, or population stratification. While our within-family GWAS and parent-offspring PGI analyses (see **Main Text** and **Figure 4**) suggest minimal aggregate indirect genetic effects on personality traits, individual SNPs might nonetheless operate through indirect pathways that bias MR analyses.

We used the TwoSampleMR and ieugwasr R packages (Hemani et al., 2018) to align and harmonize EUR population-level GWAS summary statistics for exposures and outcomes, following the standard procedure provided by package authors. To pre-process each set of summary statistics, we filtered out rare variants by excluding SNPs with EAF < .01, calculated the effective N for each SNP in summary statistics of case-control phenotypes, and selected genetic instruments for exposure by subsetting SNPs from each GWAS that were genome-wide significant ($p < 5\times10^{-8}$) and independent ($R^2 < .01$ within a 1Mb window).

To implement MR tests, we selected three estimators chosen for their robustness to violations of the three assumptions above, their robustness to heritable confounding (Sanderson et al., 2024), and their complementarity, in that each estimator makes different non-overlapping assumptions about the data (Burgess et al., 2023). The first is the weighted mode estimator, which estimates a single modal $\hat{\gamma}$ among genetic instruments under the assumption that their modal horizontal pleiotropic effect is zero (Hartwig et al., 2017). The weighted mode estimator is robust to effects of outlier instruments but is conservative and inefficient. The second, the weighted median estimator (Bowden et al., 2016), estimates a single median $\hat{\gamma}$ among genetic instruments under the assumption that fewer than half have horizontal pleiotropic effects. The weighted median estimator is more efficient than the weighted mode but also more sensitive to false positives introduced by outlier instruments. The third is MR Causal Analysis Using Summary Effect estimates (MR-CAUSE; Morrison et al., 2020). MR-CAUSE relies on a statistical methodology orthogonal to the other two estimators. In MR-CAUSE, a joint bayesian model is fit to all SNPs, where a proportion have a causal effect on the exposure and others are associated with the exposure via correlated pleiotropy. A value of $\hat{\gamma}$ is approximated using its median value in models with causal effects. All analyses were run with and without Steiger filtering, which removes SNPs where the variance explained in the outcome exceeds that in the exposure.

When reporting the results of these tests, we scaled effect size estimates to reflect the SD change in the exposure or outcome per effect allele. For binary outcomes, we estimated associations in terms of log odds ratios. Results of weighted median and weighted mode tests (**Supplementary Table S36**) can be interpreted straightforwardly as estimates of the $\hat{\gamma}$ parameter in **Supplementary Figure 37**. Results of MR-CAUSE analyses include the fits of models with and without causal effects, as well as posterior distributions of the estimated $\hat{\gamma}$ parameter (Morrison et al., 2020) (**Supplementary Table S37**). We considered an effect to be significant if the associated estimates were directionally consistent across all three MR methods (weighted median with Steiger filtering, weighted mode with Steiger filtering, and MR-CAUSE) and with 95% confidence/credibility intervals that excluded 0 across at least two methods. This is a stringent, conservative test that guards against the spurious identification of causal effects.

##### 5.5.2 Results and Causal Inference

Using these significance/credibility thresholds, we found effects for 38 exposure-outcome pairings (**Supplementary Tables S36** and **S37**). Twenty-four of these pairings indicated effects of personality traits on biobehavioral outcomes, and the remaining 14 indicated effects of biobehavioral phenotype exposures on personality traits. However, MR alone is not sufficient for causal inference. Causal inference must also be guided by considerations of the nature of the exposure and its temporal occurrence. Therefore, although we report MR tests of biobehavioral outcome exposure on personality traits (in addition to tests of personality on biobehavioral outcomes), in some cases these tests should be interpreted diagnostically, rather than reflective of causal effects. For example, longevity cannot cause personality, for very obvious reasons to do with the temporal ordering of events (personality formation precedes death). Had results indicated that longevity causes personality, this would likely mean that genetic variants that are heritable confounders of longevity and personality, such as smoking initiation, influenced the longevity GWAS (Rossoff et al., 2023) to such an extent that they overwhelmed the ability of the three MR estimators to guard against this confounding. Similarly, we deemed any detected effects of study follow-up participation on personality traits implausible, given that they constitute a lightweight finite set of behaviors unlikely to have lasting effects on typical patterns of thinking, feeling, and behaving. For the other biobehavioral outcomes tested, we deemed it plausible for these outcomes to have causal effects on personality. Research has demonstrated that personality traits develop across the lifespan (Bleidorn et al., 2022; Roberts & Yoon, 2022) and in response to events and experiences (Schwaba et al., 2023; Bühler et al., 2024) – so, exposures such as smoking initiation and educational attainment may indeed have causal effects on personality traits through lasting social and/or biological changes to person's affective, behavioral, and cognitive patterns.

In total, we identified 33 of 36 significant MR results as plausibly causal, of which 24 were effects of personality on outcomes and nine were effects of outcomes on personality. All five significant associations deemed implausible were effects of survey participation on personality traits. Our interpretation of MR findings linking study participation exposure to personality traits are that participation may be a marker of another unidentified exposure that does cause personality, or that survey participation is itself a behavioral index of personality. This also suggests that the (more plausible) effects detected of personality on study participation should be considered tentative. We present the results of these 33 exposure-outcome associations for which we do infer causal effects in **Supplementary Table S38** and describe these findings in the **Main Text.**

#### 6. Evaluating Confounding Caused by Gene-Environment Correlation

##### 6.1 Within-family PGI Rationale and Model

Population-level PGIs can be biased by confounds including population stratification (e.g., a causal environmental effect is correlated with a spurious genetic effect) and indirect genetic effects (e.g., a parent's non-transmitted genotype influences their offspring's phenotype). Within-family PGI estimation strongly reduces this confounding by predicting trait scores from one's genotype while controlling for the genotypes of their family members. Mendelian segregation during meiosis results in the random inheritance of half of the genetic material from each parent, independent of family environment. Sibling differences in genotype therefore serve as a stringent control for environmental confounding, such as uncontrolled population stratification. Moreover, because genetic information is inherited independently across chromosomes, sibling differences in genotype additionally serve as strong control for long range correlations across variants that arise from assortative mating (Tan et al., 2024; Veller & Coop, 2024). To test the extent of confounding in population-level PGI prediction, we compared population-level PGI prediction to within-family PGI prediction using dizygotic twins ascertained in the NTR and TEDS cohorts.

For both cohorts, we estimated within-family PGI associations among dizygotic twin pairs in the following fixed-effects regression model, which is closely related to the population-level PGI equation described in equation 10:

$$y = b_0 + b_1 PGI + \sum b_c x_c + \alpha_f + e \,, \qquad \text{(Eq. 14)}$$

where $\alpha_f$ are family-level fixed effects that are included to account for unmeasured between-family effects on the outcome, $b_1$ represents the *within-family effect* of the PGI (i.e. the association of the PGI with the personality trait score controlling for all sources of between-family variation), and the remaining terms are defined as in Eq. 10. Note that in this model, controls for ancestral principal components are unnecessary, as population stratification is controlled for by the family-level fixed effects. Results from these tests are described in the **Main Text**.

##### 6.2 Parent-Offspring PGI Rationale and Model

With genotype information from parents and offspring, associations between PGIs and measured traits among offspring can be decomposed into direct effects, which are those attributable to alleles transmitted from parent to offspring, and indirect effects, which are those attributable to alleles not transmitted from parent to offspring (Kong et al., 2018). Indirect genetic effects proxy factors such as the offspring's rearing environment and multi-generational social stratification processes and have been found to account for a substantial portion of PGI predictivity for traits like educational attainment (Nivard et al., 2024).

To quantify direct and indirect effects, we applied PGI weights as estimated above to families in the deCODE cohort (N = 34,506 probands with at least one genotyped parent). Phased genotype data was used to determine transmitted and non-transmitted alleles, as well as the parent-of-origin. Then, polygenic indices were computed for each of these, and used to predict a personality trait in the offspring using the following expansion of equation 10 above:

$$y = b_0 + b_1 PGI_{TP} + b_2 PGI_{TM} + b_3 PGI_{NTP} + b_4 PGI_{NTM} + \sum b_c x_c + e \,, \qquad \text{(Eq. 15)}$$

where $PGI_{TP}$ and $PGI_{TM}$ are PGIs estimated from the transmitted paternal and maternal alleles, respectively, and $PGI_{NTP}$ and $PGI_{NTM}$ are PGIs estimated from the non-transmitted paternal and maternal alleles, respectively (Kong et al., 2018).

As a matched comparison, we also estimated direct and indirect genetic effects for educational attainment based on summary data from Okbay and colleagues (2022) excluding participants from the deCODE and 23&Me cohorts (N = 718,525). We followed the same procedure for constructing PGI weights in SbayesR that we used for the Big Five personality traits to maximize comparability. Results from these comparisons are described in the Main Text and **Supplementary Table S39** and **S40.**

6.3 Within-Family GWAS Cohorts and Methods

For each Big Five trait, we conducted additional GWAS analyses on a subset of EUR cohorts (k = 12) that ascertained genomic and personality data among family members. These analyses allowed us to further probe the extent of confounding in personality genomics. Complete data on contributing cohorts for these analyses can be found in **Supplementary Table S41.**

Analysts followed one of two approaches to estimate these models. For k = 2 cohorts (NTR and EBB), analysts used snipar (Young et al., 2022) to pool genetic information across parents and siblings (for the Netherlands Twin Register cohort) or to impute synthetic parents from sibling genetic data and then pool this genetic information (for the Estonian Biobank cohort). Complete details of this imputation procedure can be found in Young and colleagues, 2022 and Tan and colleagues, 2024. When genetic data are provided on a participant and their parents, and when trait means are standardized to a mean of 0, the GWAS equation in equation 1, above, estimates the genotype-phenotype association in the population, conditional on covariates. We refer to this the *population-level effect*. Then, incorporating parent genotype data, this model can be elaborated for each SNP:

$$y = b_0 + \delta SNP_i + \alpha_m SNP_m + \alpha_f SNP_f + \sum b_c x_c + e \,, \qquad \text{(Eq. 16)}$$

where $SNP_i$ is the effect allele count for a given SNP for the individual offspring, $SNP_m$ is the allele count for the same SNP for that individual's mother, and $SNP_f$ is the allele count for that SNP for the same SNP for that individual's father. In this equation, $\alpha_m$ and $\alpha_f$ estimate the associations between one's parents' genotypes and trait scores, and $\delta$ thus estimates the genotype-phenotype association for an individual conditional on covariates and one's parents' genotypes. We focus on the $\delta$ term, which we refer to as the *within-family effect*.

For k = 10 cohorts, analysts implemented the pipeline developed in Howe and colleagues (2022), which uses a different, comparable method for decomposing the genotype-phenotype association into population-level and within-family effects. Rather than using information from one's parents, it estimates effects among siblings. Specifically, the population-level estimate is obtained in the same way as equation 1, but with standard errors clustered by sibship to account for non-independent genotypes between siblings. The within-family estimate is obtained using the following equation:

$$y = b_0 + \delta SNP_i + \alpha SNP_F + \sum b_c x_c + e \,, \qquad \text{(Eq. 17)}$$

where $SNP_F$ is the mean effect allele count for all siblings in the family, and $SNP_i$ is the allele count for that SNP for the individual sibling, centered around $SNP_F$ via subtraction. Standard errors of effects are adjusted for sibling cluster. Thus $\delta$ estimates the genotype-phenotype association for an individual, conditional on covariates and mean sibling genotypes.

Within family genetic effects ($\delta$) estimated from the snipar and Howe and colleagues models are comparable but not equivalent. The major conceptual difference between the two models is that the snipar model estimates within-family effects by controlling for parental genotype(s), whereas the Howe and colleagues model estimates within-family effects by controlling for sibling genotype(s). Evaluating these differences empirically is a worthwhile topic for future research.

Notably, Howe and colleagues (2022) previously estimated population-level and within-family effects for neuroticism. We incorporated these data, considered as one cohort, into the present meta-analysis of neuroticism. To remove overlapping participants with the other cohorts that contributed data to this analysis, analysts from Howe and colleagues provided us with re-estimated meta-analytic neuroticism summary statistics, excluding data from the NTR cohort (for which we had a larger sample that also estimated effects using snipar). Then, when synthesizing data, we excluded the remaining k = 7 cohorts that provided neuroticism summary statistics that overlap with the Howe results. These meta-analytically provided Howe summary statistics did not include imputation quality scores, which are necessary for stringent cohort-level QC. To estimate imputation quality for each SNP, we conducted a sample-size-weighted meta-analysis of INFO scores across the cohorts that contributed both to this study and the Howe and colleagues study (see Equation 6 above). As the Danish Twin Registry and HUNT studies contributed to the Howe and colleagues study but not to this study, we estimated INFO for SNPs in these studies using the STAGE and MoBa cohorts, respectively, as these matched in nationality and genotyping chip and thus provided the closest available estimate to what imputation quality would be in these missing datasets (**Supplementary Table S1**).

##### 6.4 Within-Family GWAS Quality Control

We then conducted stringent cohort-level QC on both population-level and within-family results. The QC procedure mirrored the full population-level analyses presented in section 1, with the following modifications.

First, we ensured that personality trait scores were standardized in one step, rather than separately for population-level and within-family models, so that their variances remained on the same scale. Second, we restricted imputed SNPs in each cohort to those with extremely high-quality (INFO scores ≥ .99); lower quality imputation, which introduces measurement error, has been shown to substantially deflate sibling correlations below their expected value (Tan et al., 2024). Third, as within-family associations are necessarily estimated with less statistical power than population-level associations, we pruned the small number of SNPs (in most cohorts, 0) for which the within-family standard error was smaller than the population-level standard error. Fourth, to check allele alignment across population-level and within-family data, we correlated the betas between these two methods and ensured that they were positive.

We identified during cohort-level quality control that, especially in smaller cohorts estimated using the Howe and colleagues (2022) pipeline, there appeared to be downward bias in $h^2$SNP estimates. This may in part be caused by small-sample deviations from the expected Z-distribution among SNPs with low MAF. For these SNPs, it may be more appropriate to estimate effects using a t-distribution. To implement this correction, we re-scaled Z-statistics of each SNP into t-scores with the degrees of freedom approximated from MAF-estimated variance and sample size, as follows:

$$\widehat{df} = 2MAF \times (1 - MAF) \times N. \quad \text{(Eq. 18)}$$

As shown in **Supplementary Figure S38**, this correction had minimal effect on SNPs estimated with larger effective sample sizes, while adjusting Z-statistics of less common SNPs. LDSC results of each cohort analyzed with the Howe and colleagues pipeline with this correction implemented are presented in **Supplementary Table 41**. As this correction only applies to smaller cohorts, and we excluded SNPs with estimated sample size < 5,000 from meta-analyses, this correction was not necessary for meta-analytically synthesized data. Consistent with near-complete sample overlap between population-level and within-family data for each cohort, cross-trait intercepts were high (mean CTI = .52). Most LDSC estimated genetic covariances between population-level and within-family associations in the same cohort were negative in smaller cohorts (**Supplementary Table 41**).

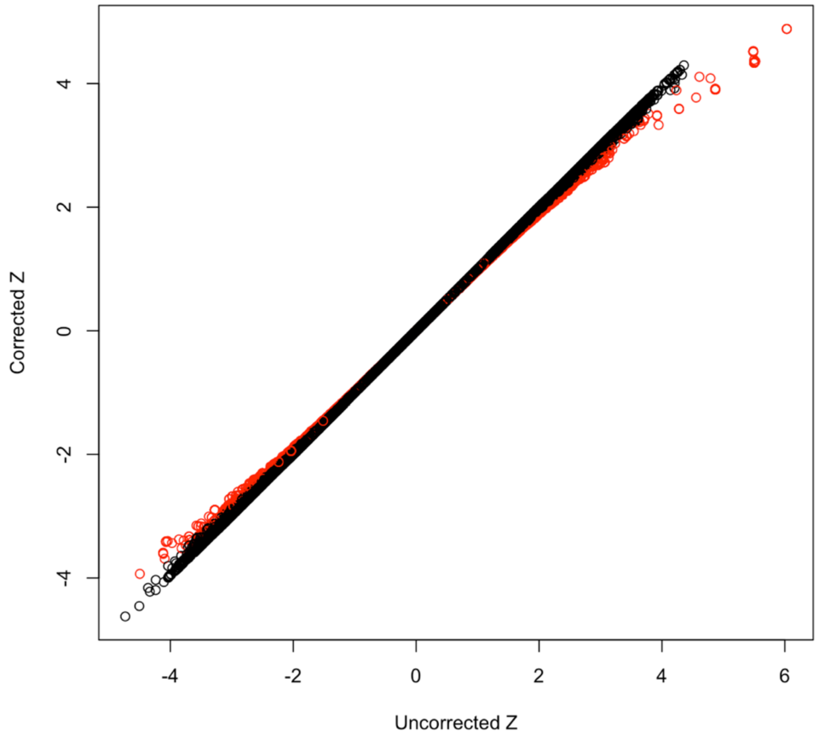


**Supplementary Figure S38. Deviations from the expected within-family Z-statistics among low-MAF SNPs in the Minnesota Center for Twin and Family Research Cohort (N = 942).** Note: SNPs with effective degrees of freedom < 50 are highlighted in red, whereas SNPs with greater degrees of freedom are depicted with black.

#### 6.5 Inverse Variance-Weighted Meta-Analysis

After, cohort-level QC, we conducted inverse variance-weighted meta-analysis for each trait on matched cohorts of population-level and within-family data, using Equation 5, above.

As with the population-level GWAS in the full set of cohorts, we re-estimated sample size for each SNP using the meta-analytic beta, standard error, and minor allele frequency (see Equation 7 above). To ensure comparability between population-level and within-family results, we approximated sample size using the MAF corresponding to the 1000 genomes 3v5 EUR panel, rather than the meta-analytic MAF estimated from contributing cohorts. Population-level estimated sample sizes were capped at the summed observed sample size across contributing cohorts, and within-family sample sizes were capped at the 75th percentile of the summed observed sample size across contributing cohorts. This threshold was chosen because, within each of the k = 12 contributing cohorts to these analyses, the maximum observed N fell between the 60th and 75th percentile of the estimated Nhat.

6.6 Within-Family GWAS Results

Across each of the Big Five traits, we did not identify any genome-wide significant lead SNPs from the meta-analytic within-family GWAS results (minimum $p = 2.26 \times 10^{-7}$). The mean chi-squared value of associations ranged from 1.04 (for agreeableness) to 1.09 (for neuroticism). We present Manhattan plots for each of these analyses below, in **Supplementary Figures S39-S43.**

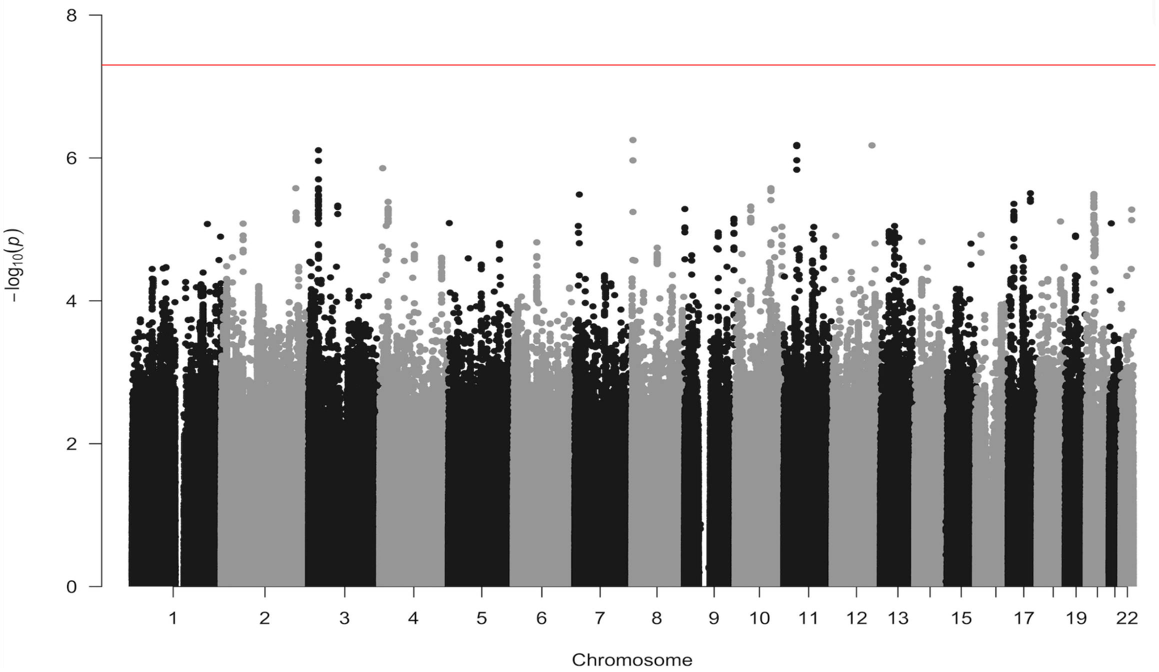


**Supplementary Figure S39. Manhattan plot of within-family GWAS of extraversion in participants of EUR-like ancestry (N = 47,630)**. Note: red line denotes genome-wide significance threshold at $p = 5 \times 10^{-8}$.

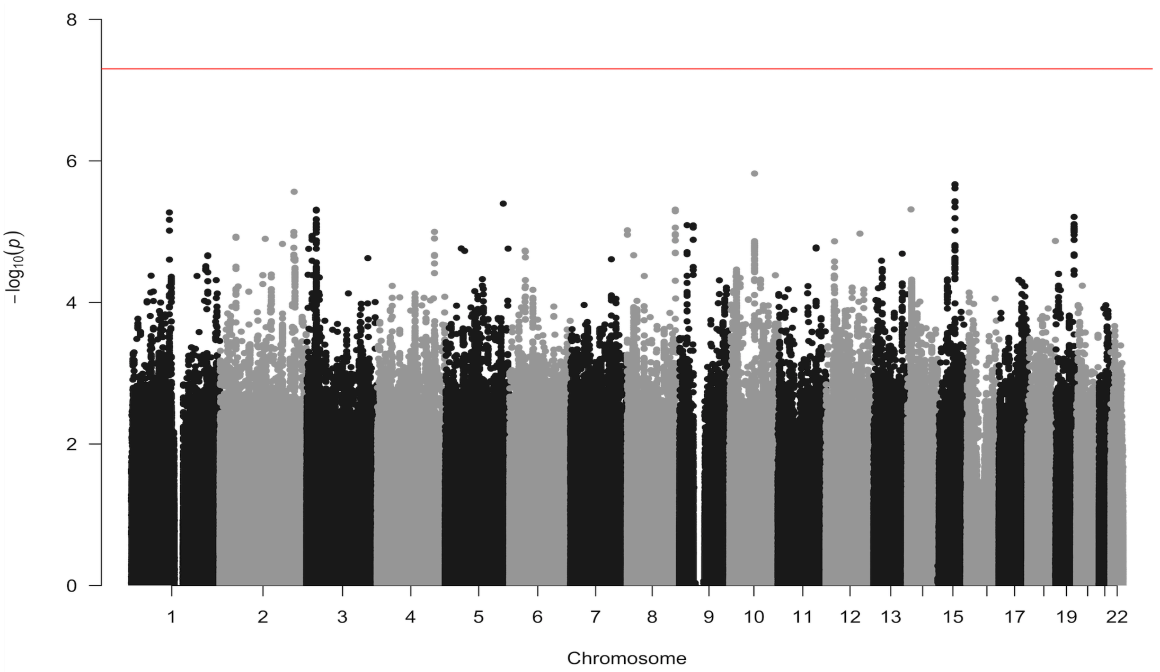


**Supplementary Figure S40. Manhattan plot of within-family GWAS of agreeableness in participants of EUR-like ancestry (N = 31,607)**. Note: red line denotes genome-wide significance threshold at $p = 5\text{x}10^{-8}$.

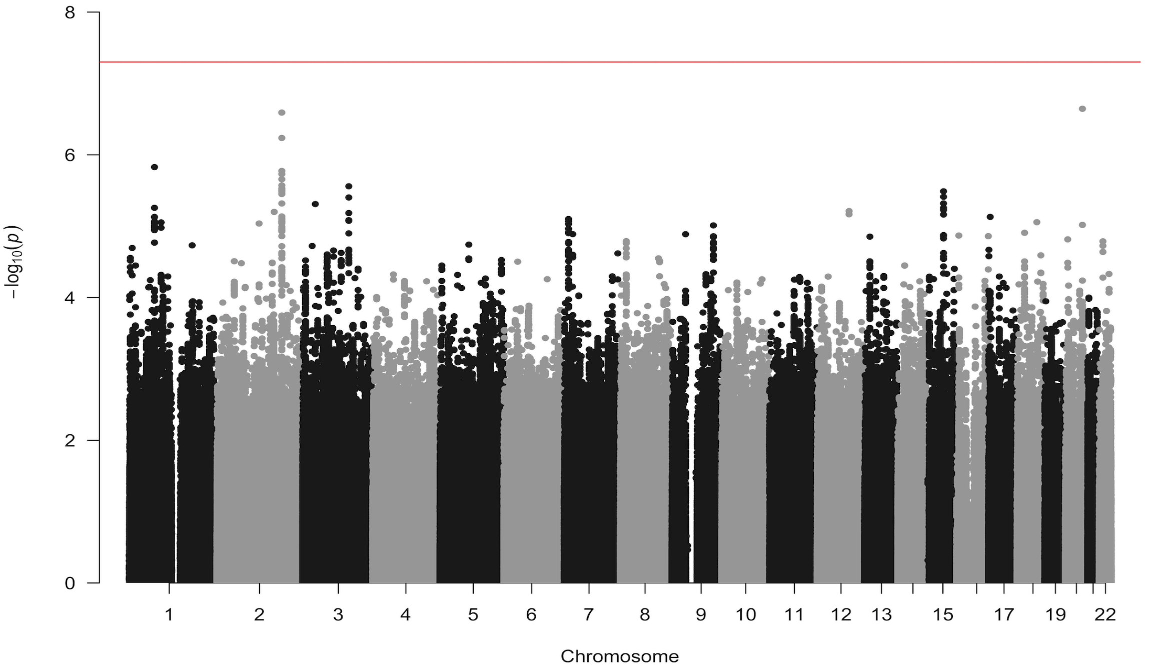


**Supplementary Figure S41. Manhattan plot of within-family GWAS of conscientiousness in participants of EUR-like ancestry (N = 31,544)**. Note: red line denotes genome-wide significance threshold at $p = 5\text{x}10^{-8}$.

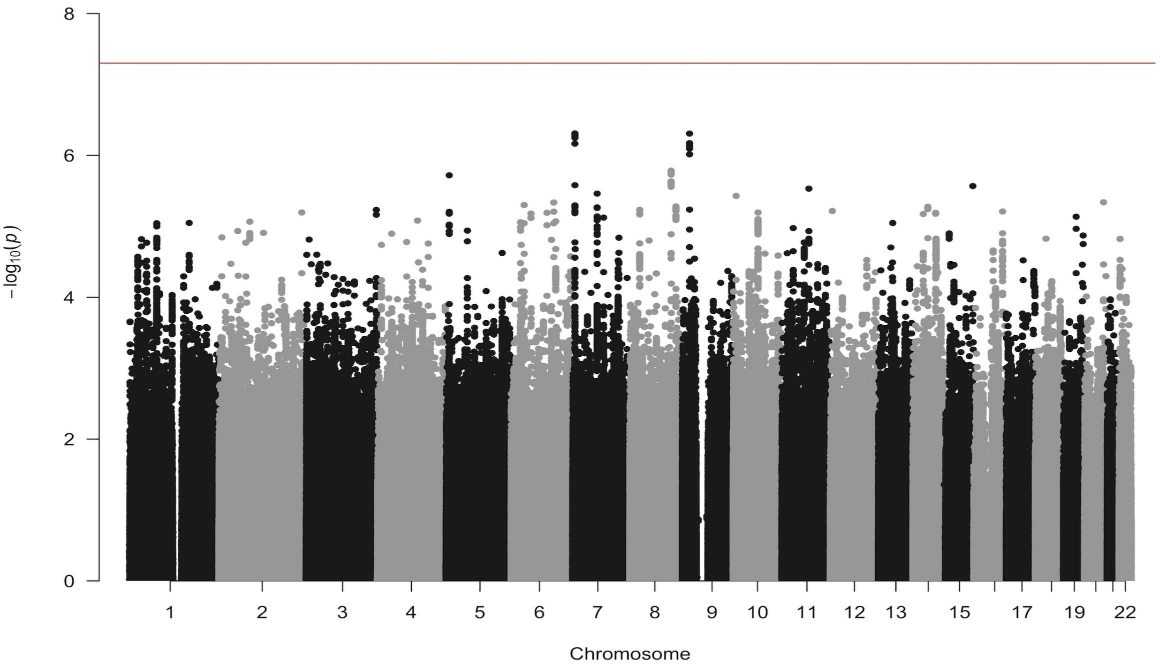


**Supplementary Figure S42. Manhattan plot of within-family GWAS of neuroticism in participants of EUR-like ancestry (N = 55,637)**. Note: red line denotes genome-wide significance threshold at $p = 5 \times 10^{-8}$.

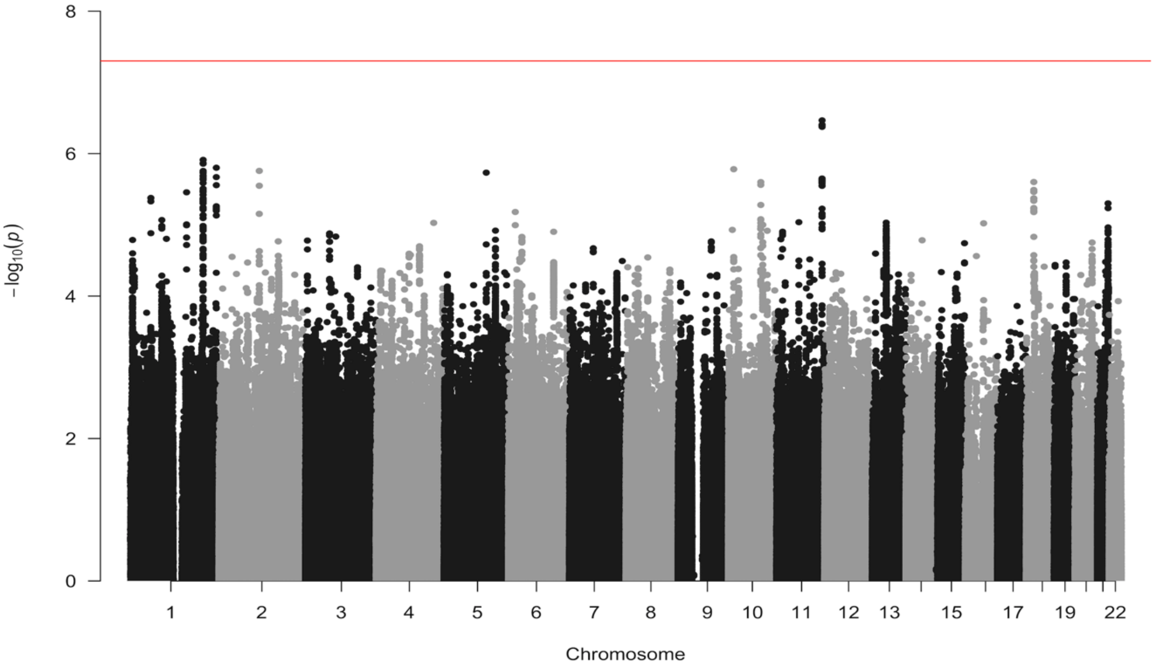


**Supplementary Figure S43. Manhattan plot of within-family GWAS of openness to experience in participants of EUR-like ancestry (N = 32,216)**. Note: red line denotes genome-wide significance threshold at $p = 5 \times 10^{-8}$.

6.7 Comparison of Population-Level and Within-Family Genetic Architectures

When summary statistics from GWAS restricted to cohorts contributing within-family data were submitted to LDSC, we observed nontrivial intercept inflation for both population-level data (mean intercept = 1.06, SE = 0.01) and within-family data (mean intercept = 1.07, SE = 0.01; **Table S42**), consistent with the inflation identified in Howe and colleagues (2022). Note that there was not appreciable intercept inflation in the full population-level GWAS (**Table 1**). For each trait, population-level and within-family $h^2$SNP was significant and substantial. We estimated genetic correlations between population-level and within-family associations, appropriately rescaling for highly-overlapping cohorts, using the rgmodel() function in the GenomicSEM R package (Ennis et al., 2025).

We then tested equivalence of population-level and within-family genetic architecture for each trait using Genomic SEM. To test equivalence of heritability, we compared the fit of two bivariate models to the data: a fully saturated model with 0 degrees of freedom, in which population-level and within-family heritability are freely estimated and their covariance is freely estimated, with a heritability-constrained model in which population-level and within-family heritability are constrained to be equal and their covariance is freely estimated. The chi-square nested model comparison between these two models, which differ in 1 degree of freedom, tests the hypothesis that heritability is equivalent across the two methods. For openness to experience, but not the other Big Five traits, heritability significantly differed at $p < .05$ ($\chi^2(1) = 5.80$, $p = .016$).

To test whether genetic correlations were significantly less than 1, we compared the fully saturated model described above to a correlation-constrained model, in which population-level and within-family heritability were freely estimated and their correlation is constrained to unity. Chi-square model tests indicated that genetic correlations across within-family and population-level results were significantly less than 1 for agreeableness ($r = .79$) ($\chi^2(1) = 4.53$, $p = .033$) and neuroticism ($r = .75$) ($\chi^2(1) = 11.53$, $p = .0006$). We present full results of these analysis in the **Main Text** and **Supplementary Table S42**.

We then compared the average SNP effect sizes between population-level and within-family data. A regression model comparison of raw SNP-trait associations between the two sets of data is uninformative: this regression is downwardly biased by error in both the population-level and within-family effect estimates as these GWAS are relatively small and there is dependency in these errors due to sample overlap. To account for this and enable appropriate comparison, we took two steps.

First, we restricted effect size comparisons to the set of SNPs for each trait that have maximal statistical signal. We identified the set SNPs in the full EUR GWAS meta-analysis with $p < 1x10^{-5}$ and sequentially pruned SNPs within +/- 250kb of each other (from most strongly associated to least strongly associated).

Second, among these SNPs, we compared effect sizes in terms of a ratio. In the numerator was the regression slope of within-family SNP-trait effect sizes (from the within-family GWAS) on the population-level GWAS SNP-trait effect sizes (among EUR cohorts that did not contribute to the within-family GWAS). In the denominator was the regression slope of population-level SNP-trait effect sizes (from the within-family GWAS) on the full EUR population-level GWAS SNP-trait effect sizes (among cohorts that did not contribute to the within-family GWAS). By regressing within-family GWAS estimates on non-overlapping population-level GWAS estimates, we minimize overlap between cohorts and reduce correlated error.

This ratio can be interpreted as the relative strength of replication of estimates across within- and population- levels of analysis: after holding out overlapping cohorts, how strongly do within-family and population-level effects from the within-family GWAS replicate population-level effects in the full EUR GWAS? The slope of each regression in the numerator and denominator is expected to be 1 in the absence of biasing factors with differential effects, such as effect size shrinkage due to winners' curse, measurement differences, heritability differences, and peculiarities of individual cohorts. As shown in **Supplementary Table S42**, the average strength of replication of within-family data is .90, and the average strength of replication of population-level data is .94. However, these biasing factors are expected to affect the two regression betas equally, meaning the ratio of the two regression slopes is expected to be below 1 only in the presence of gene environment correlation introduced by assortment, dynastic or genetic nurture effects. Indeed, the average ratio between the two replication effect sizes is .95 (ranging from .86 for conscientiousness to 1.06 for agreeableness), indicating very little confounding due to these factors.

Finally, we compared genetic correlations across the Big Five traits between EUR population-level data, EUR within-family data, and AFR population-level data, using bivariate LDSC. These associations revealed similarity in personality's cross-trait pleiotropy across these sources (**Supplementary Figure S44**). Specifically, correlations were sign-concordant across the three methods and were highly similar between population-level and within-family EUR data. Associations among the traits in AFR population-level data were stronger in magnitude than in EUR data, but with correspondingly larger standard errors.

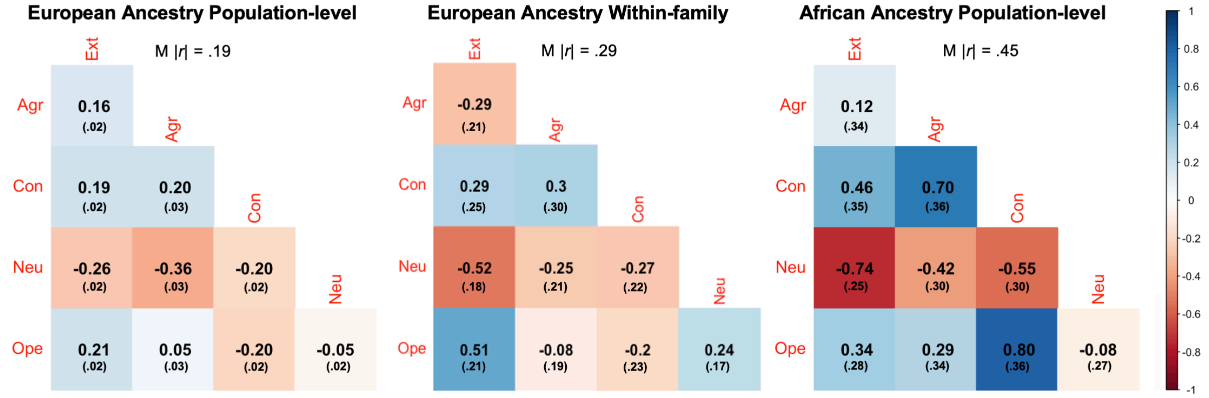


**Supplementary Figure S44. Correlations among the Big Five in EUR population-level, EUR within-family, and AFR population participant data.** Note: Ext = Extraversion. Agr = Agreeableness. Con = Conscientiousness. Neu = Neuroticism. Ope = Openness to experience. Numbers in parentheses depict standard errors. Genetic correlations estimated using LDSC.

### Cohort Information

#### 23andMe, Inc. (23andMe)

The 23andMe cohort consists of 59,225 participants who completed the Five Dimensions of Personality survey module provided at the company's website, which measured personality traits using the 44-item BFI (John et al., 1991). Participants were those with >97% European ancestry filtered to unrelated individuals who shared less than 700 cM IBD. For complete details regarding this cohort, see Lo and colleagues (2017). We would like to thank the research participants and employees of 23andMe, Inc. for making this work possible.

John, O. P., Donahue, E. M., & Kentle, R. (1991). The Big Five Inventory. Technical report, University of California, Berkeley

Lo, M.T., Hinds, D.A., Tung, J.Y., Franz,C., Fan, C.C., Wang, Y., Smeland, O.B., Schork, A., Holland, D., Kauppi, K., Sanyal, N., Escott-Price, V., Smith, D.J., O'Donovan, M.C., Stefansson, H., Bjornsdottir, G., Thorgeirsson, T.E., Stefansson, K., McEvoy, L.K., Dale, A.M., Andreassen, O.A. & Chen, C.H. (2017). Genome-wide analyses for personality traits identify six genomic loci and show correlations with psychiatric disorders. *Nature Genetics*, *49*(1), 152-156.

#### Australian Genetics of Depression Study (AGDS)

The recruitment and sample characteristics of the Australian Genetics of Depression Study (AGDS) have been described in detail elsewhere (Byrne et al 2020). Briefly, more than 21,000 participants were recruited (15,792 currently genotyped) through a dual recruitment approach whereby cases were recruited either through Australian government prescription records or through a media campaign. Once enrolled in the study, participants completed online questionnaires comprising an obligatory core module that assessed MDD diagnosis as well as a range of satellite modules assessing a variety of phenotypes. The personality phenotypes of Extraversion and Neuroticism were measured using the Eysenck Personality Questionnaire (EPQ). AGDS was primarily funded by National Health and Medical Research Council (NHMRC) of Australia grant (1086683).

Byrne, E.M., Kirk, K.M., Medland, S.E., McGrath, J.J., Colodro-Conde, L., Parker, R., Cross, S., Sullivan, L., Statham, D.J., Levinson, D.F., Licinio, J., Wray, N.R., Hickie, I.B., & Martin, N.G. (2020). Cohort profile: the Australian genetics of depression study. *BMJ Open*, *10*(5), e032580.

#### Avon Longitudinal Study of Parents and Children (ALSPAC)

The Avon Longitudinal Study of Parents and Children (ALSPAC) is a prospective population-based study that initially recruited pregnant women in the Avon district of England in 1990-1992 (Northstone et al., 2019). When children were 13.5 years old, parents reported on their adolescent child's personality traits using the 50-item International Personality Item Pool inventory (IPIP-50), which measures the Big Five traits with ten questions each (Goldberg, 1999). In this study, we restricted genetic analyses to the N = 4,284 adolescents (74% females) with personality trait data who were unrelated (<10% of alleles shared IBD) and of European ancestry (clustered within the CEU HapMap2 population using multidimensional scaling of genome-wide IBS pairwise distances).

Goldberg, L.R. (1999). A broad-bandwidth, public domain, personality inventory measuring the lower-level facets of several five-factor models. In I. Mervielde, I.J. Deary, F. De Fruyt, & F. Ostendorf (Eds.). *Personality Psychology in Europe*, Vol. 7, Tilburg University Press, Tilburg.

Northstone, K., Lewcock, M., Groom, A., Boyd, A., Macleod, J., Timpson, N., & Wells, N. (2019). The Avon Longitudinal Study of Parents and Children (ALSPAC): An update on the enrolled sample of index children in 2019. *Wellcome Open Research*, *4*.

**BiDirect Study**

The BiDirect Study examines the association between depression and subclinical arteriosclerosis and includes three distinct cohorts recruited in the district of Münster, Germany: 1) patients hospitalized for an acute episode of depression in one of five psychiatric hospitals, 2) patients recruited 2–4 months after an acute coronary event in cardiac rehabilitation facilities, and 3) a random sample of the inhabitants of the city of Münster. The joint ethics committee of the Westphalian Chamber of Physicians and the University of Münster approved the study (2009-391-f-s). All participants in the three cohorts were aged 35 to 65 years at baseline and examined in parallel with identical methods. Overall, 2,258 individuals were included at baseline (among them 946 in cohort 3) and 1,816 (824 in cohort 3) of them participated in the second examination wave. (Effective sample: 1,467 (unrelated) individuals (692 females, 775 males; mean age: 56 +/- 7.8 years.)

The examination program in each wave included a medical interview on socioeconomic factors, comorbidities and medication, a depression classification including subtypes, a vascular diagnostic work-up, anthropometric measurements, a neuro-psychological test battery and a magnetic resonance imaging (MRI) of the brain. A battery of interview-based or self-administered questions and scales (e.g., depressive and anxiety symptoms from the M.I.N.I. vs5.0.0, childhood trauma, personality, resilience, and quality of life) was also assessed. In addition, blood samples (serum, plasma, EDTA) were collected in each wave and DNA and RNA once. Between 2010 and 2021 four in-person examination waves were performed, with mean follow-up intervals of 2.5 years. The Big-5 personality traits were collected in examination wave 2 and the 15-item questionnaire BFI-S used as part of the self-report questionnaires. BiDirect is funded by German Ministry of Research and Education (BMBF, No. 01ER0816 + 01ER1506).

Linnemann P., Friedrich N., Nauck M., Teismann H., & Berger K. (2022). The relationship between cortisol awakening response and trait resilience in two patient cohorts and one population-based cohort. *World Journal of Biological Psychiatry, 14*, 1-10.

Schneider, G., Köhnke, C., Teismann, H., & Berger, K. (2021). Childhood trauma and personality explain more variance in depression scores than sociodemographic and lifestyle factors - Results from the BiDirect Study. *Journal of Psychosomatic Research, 147,* 110513.

Teismann, H., Wersching, H., Nagel, M., Arolt, V., Heindel, W., Baune, B.T., Wellmann, J., Hense, H.W., & Berger, K. (2014). Establishing the bidirectional relationship between depression and subclinical arteriosclerosis – rationale, design, and characteristics of the BiDirect Study. *BMC Psychiatry*, *14*, 174.

**Baltimore Longitudinal Study of Aging (BLSA)**

The National Institute on Aging's Baltimore Longitudinal Study of Aging (BLSA) is an ongoing longitudinal study of community-dwelling volunteers living primarily in the Baltimore-Washington area (USA), who have been continually recruited since 1958 (Ferrucci, 2008). Participants are healthy at study entry and undergo extensive evaluations including medical evaluations and physiological, functional, and neuropsychological testing at each clinic visit. Longitudinal personality assessments (NEO-PI-R) were aggregated to provide more reliable estimates. There are 841 individuals of European ancestry with genotyping and personality data available. 455 are male and 386 female, and year of birth ranged from 1902 to 1979. The BLSA study is supported in part by the Intramural research program of the National Institute on Aging, NIH – Baltimore MD USA

Ferrucci,L. (2008). The Baltimore Longitudinal Study of Aging (BLSA): a 50-year-long journey and plans for the future. *Journal of Gerontology, Series A, Biological Sciences and Medical Sciences, 63*(12), 1416-1419.

Kang, H.M., Sul, J.H., Service, S.K., Zaitlen, N.A., Kong, S.Y., Freimer, N.B., Sabbati, C., & Eskin, E. (2010). Variance component model to account for sample structure in genome-wide association studies. *Nature Genetics*, *42*(4), 348-354.

**Cilento Study**

The Cilento study is a population-based study that includes 2,331 individuals from three isolated populations of South Italy. Cilento participants rated themselves on a 60-item version of the NEO Personality Inventory (NEO-PI-R), the NEO-FFI. Personality scores for the five factors are based each on based on 12 items. Factors from the NEO-PI-R questionnaire were available for 849 genotyped subjects. Of this sample, 63% were women. The mean age of all participants was 54.34 (SD=19.25), of the men 54.54 (SD=18.9) and of the women 54.23 (SD= 19.45). The Cilento study was supported by the "National Biodiversity Future Center" on 'Biodiversity', financed under the National Recovery and Resilience Plan (NRRP), Mission 4, Component 2, Investment 1.4 "Strengthening of research structures and creation of R&D 'national champions' on some Key Enabling Technologies"; the CNR projects FOE-2021- DBA.AD005.225 and FOE 2022 - DSB.AD006.371.

Nutile, T., Ruggiero, D., Herzig, A. F., Tirozzi, A., Nappo, S., Sorice, R., Marangio, F., Bellenguez, C., Leutenegger, A.L., & Ciullo, M. (2019). Whole-exome sequencing in the isolated populations of Cilento from South Italy. Scientific Reports, 9(1), 4059.

Costa, P.T., & McCrae, R.R. (1992). Professional manual for the NEO-PI-R and NEO-FFI. Odessa, FL: Psychological Assessment Resources.

#### The Collaborative Study on the Genetics of Alcoholism (COGA)

The Collaborative Study on the Genetics of Alcoholism (COGA) is a large-scale family study designed to identify genes that affect the risk for alcoholism (i.e., alcohol dependence) and alcohol-related characteristics and behaviors. This collaborative project is funded by the National Institute on Alcohol Abuse and Alcoholism. Data collection, analysis, and/or storage for this study takes place at eleven sites across the United States. COGA has gathered detailed, standardized data on study participants, including diagnostic and neurophysiological assessments. The age range of the sample used in these analyses is between 11 and 34, with a mean of 20 years (SD=3.5). Among participants with genome-wide data, 1,751 (823 males, 891 females) individuals contributed to European ancestries, and 859 (414 males, 445 females) individuals contributed to African ancestries.

COGA used the NEO Five-Factor Inventory (NEO-FFI), a shortened version of the revised NEO Personality Inventory (NEO PI-R). NEO-FFI is a 60-item self-report questionnaire designed to give quick, reliable, and valid measures of the five domains of adult personality. The NEO-FFI was administered to subjects aged 13 and older in Phases II-III of COGA. It is administered in Phase IV to subjects aged 12 and older at baseline if not completed previously. It is re-administered once to subjects who age up to at least 17 at follow-up.

Costa P.T., & McCrae R.R. (1986). Cross-sectional studies of personality in a national sample: 1. Development and validation of survey measures. *Psychology and Aging*, *1*(2), 140-143.

Costa, P.T., & McCrae, R.R. (1992). Professional manual: revised NEO personality inventory (NEO-PI-R) and NEO five-factor inventory (NEO-FFI). Odessa, FL: Psychological Assessment Resources, 61.

#### deCODE Study on the Genetics of Personality

Icelandic data were collected through deCODE studies approved by the National Bioethics Commmittee (approval no. VSN-09-098, VSN-18-113 and VSN-19-196) following review by the Icelandic Data Protection Authority. The 60-item NEO-Five Factor Inventory (NEO-FFI; Bjornsdottir et al., 2014; Costa & McCrae, 1992) was administered to a sample of approximately 39,700 participants, evaluating neuroticism, extraversion, openness, agreeableness and conscientiousness. The dataset consists of a combination of two cohorts; one from a study on the genetics of addiction (N~6,700; Bjornsdottir et al., 2014) and another from an online population study on personality (N~33,000; Jonsdottir et al., 2021). For individuals that participated in both studies, scores from the first participation were included in the analysis. As the Icelandic population is a homogeneous one with extensive genealogy information, instead of using principal components to adjust for population structure, participants that did not have at least three grandparents recorded in the genealogy database were excluded from the analysis and the data subsequently adjusted for county of origin. The scores were also adjusted for sex, age and age squared, and then subjected to rank-based inverse normal transformation. Participants' age ranged from 18 to 90 (M=46.5, SD=13.4) and 66% of them were females.

Bjornsdottir, G., Jonsson, F. H., Hansdottir, I., Almarsdottir, A. B., Heimisdottir, M.,

Tyrfingsson, T., ... & Thorgeirsson, T. E. (2014). Psychometric properties of the Icelandic NEO-FFI in a general population sample compared to a sample recruited for a study on the genetics of addiction. *Personality and individual differences*, 58, 71-75.

Costa Jr, P. T. (1992). Revised NEO personality inventory and NEO five-factor inventory. Professional manual.

Jonsdottir, G. A., Einarsson, G., Thorleifsson, G., Magnusson, S. H., Gunnarsson, A. F., Frigge, M. L., ... & Stefansson, K. (2021). Genetic propensities for verbal and spatial ability have opposite effects on body mass index and risk of schizophrenia. *Intelligence,* 88, 101565.

**Estonian BioBank (EBB)**

Estonian Biobank (EBB) was started in 1999, and the first samples were collected in 2002 (Milani et al., 2024). EBB incorporates over 210,000 participants, 20% of adults in Estonia. During the first decade, recruitment occurred through random sampling of visitors of general practitioners across the country. From 2017, the recruitment was volunteer-based. The participants have completed an extensive phenotype questionnaire about their demographic, genealogical, educational, and occupational backgrounds, lifestyle, health status, and medical history. Between November 2021 and April 2022, 73,986 participants included in this study completed the 100 Nuances of Personality (Henry & Mõttus, 2022; 52,216 female and 21,767 male, mean age 47.6, sd=14.6, range 18-102), along with 21,986 informant reports. Big Five domains were estimated with a PCA, with 12 best-loading items per domain. PCA resulted in very low correlations between domains (Anni et al., 2024). More details can be seen in the behavioral cohort paper (Vaht et al., 2024).

The activities of the EBB are regulated by the Human Genes Research Act, which was adopted in 2000 specifically for the operations of the EBB. Individual level data analysis in the EBB was carried out under ethical approvals of 1.1-12/1409 and 1.1-12/2161 from the Estonian Committee on Bioethics and Human Research (Estonian Ministry of Social Affairs), using data according to release application 6-7/GI/31997 from the Estonian Biobank. This study was funded by European Union through the European Regional Development Fund Project No. 2014-2020.4.01.15-0012 GENTRANSMED. Data analysis was carried out in part in the High-Performance Computing Center of University of Tartu. This work was also funded by Estonian Research Council's personal research funding start-up grants PSG656 and PSG759, and Estonian Research Council's team grants PRG2190 and PRG1291.

Anni, K., Vainik, U., & Mõttus, R. (2024). Personality profiles of 263 occupations. *Journal of Applied Psychology*.

Milani, L., Alver, M., Laur, S., Reisberg, S., Haller, T., Aasmets, O., ... & Metspalu, A. (2024). From Biobanking to Personalized Medicine: the journey of the Estonian Biobank. *medRxiv,* 2024-09.

Vaht, M., Arumäe, K., Realo, A., Ausmees, L., Allik, J., Henry, S., ... & Vainik, U. (2024). Cohort profile: personality measurements at the Estonian Biobank of the Estonian Genome Center, University of Tartu.

**FinnTwin**

The Finnish Twin Cohort (FTC) consists of three cohort studies, the older cohort of twins born before 1958 with both co-twins alive in 1974, twins born 1974-1979 (the FinnTwin16 study) and twins born 1983-1987 (the FinnTwin12 study).

FinnTwin12:

The FinnTwin12 (FT12) study includes all Finnish twins born in 1983–1987 ($N = 6{,}272$). From the Finnish population registry (which covers the entire population), the twins were identified as those born on the same day to the same mother. The baseline questionnaires were sent to the twins and parents in the year the twins turned 11, with the next two waves being at ages 14 and 17. Embedded in the study was a more detailed phenotyping targeting 1,035 families, from which 1,852 twins were interviewed at age 14. These twins were invited again as young adults (wave 4), and 1,347 participated. This current study is based on this subset of the fourth wave of the FinnTwin12 study. Personality was measured with the 55-item version of the NEO FFI (Costa & McCrae, 1992). The Finnish version of the NEO FFI is based on a longer 180-item personality inventory, which is an authorized adaptation of the NEO Personality Inventory (Pulver et al., 1995). Seven extra items for a sensation-seeking facet of the extraversion scale were also used in our study. There were total of 1,253 genotyped samples available (573 men, 680 women), with ages ranging 19-26 (SD=0.72)

Older Finnish Twins study:

The older part of the Finnish Twin Cohort consists of all Finnish twin pairs of the same gender born before 1958 with both co-twins alive in 1974. These twin pairs were selected from the Central Population Registry of Finland in 1974. Four surveys of the entire cohort have been carried out. The first questionnaire was mailed to all pairs in August-October 1975. Three follow-up questionnaire studies have been carried out in 1981, 1990, and 2011. Personality measures for extraversion and neuroticism were derived using a short form of the Eysenck Personality Inventory (Eysenck & Eysenck, 1968). There were a total of 7,544 samples (3,350 men and 4,194 women) with genotype data available for an age range of 18-70 (SD=9.6).

Nicotine Addiction Genetics Finland Study (NAG-FIN):

The study sample was drawn from the population-based Finnish Twin Cohort Study, which consists altogether of 35,834 adult twins born in 1938–1957. Twin pairs concordant for ever smoking were identified and recruited along with their family members (mainly siblings) for the Nicotine Addiction Genetics (NAG-FIN) Finland study. Priority was given to heavier smokers. The data collection took place in 2001–2005. Participants were assessed by DNA sample collection and a structured diagnostic psychiatric interview resulting in detailed phenotypic information on multiple smoking behavior traits. Personality measures were the same as with FT12 study. There were total of 2013 genotyped samples available (962 men and 1,051 women), age ranging from 32-92 (SD=8.5).

Costa, P.T. & McCrae, R.R. (1992). Professional manual for the NEO-PI-R and NEO-FFI. Odessa, FL: Psychological Assessment Resources.

Eysenck, H. J., & Eysenck, S. B. (1968). Eysenck personality inventory. Journal of Clinical Psychology.

Pulver A., Allik J., Pulkkinen L., & Hämäläinen M. The Big Five Personality Inventory in two non-Indo-European languages. *European Journal of Personality*, *9*(2), 109–124.

#### Genes for Good

The Genes for Good study uses social media to engage a large, diverse participant pool in genetics research and education (Brieger et al., 2019). Health history and daily tracking surveys are administered through a Facebook application, and participants who complete a minimum number of surveys are mailed a saliva sample kit to collect DNA for genotyping. As of 2022, a total of 20,700 participants are both genotyped and have completed surveys. Participants completed the 44-item Big Five inventory (BFI), which measures extraversion, agreeableness, conscientiousness, neuroticism, and openness to experience. Descriptive statistics for the BFI in the Genes for Good sample have been reported previously (Liu et al., 2020). After classifying the ancestry of participants and removing individuals with excessive homozygosity or heterozygosity, we perform GWAS on the self-reported personality traits from the BFI in a total of 19,199 individuals (EUR=16,553, AFR=684, LAT=1,444, and EAS=518). Participants are aged between 20-90 (M=35.75, SD=12.88) and 72% of them are females.

Brieger, K., Zajac, G.J., Pandit, A., Foerster, J.R., Li, K.W., Annis, A. C., Schmidt, E.M., Clark, C.P., McMorrow, K., Zhou, W., Yang, J., Kwong, A.M., Boughton, A.P., Wu, J., Scheller, C. Parikh, T., de la Vega, A., Brazel, D.M., Frieser, M., ... Abecasis, G.R. (2019). Genes for good: engaging the public in genetics research via social media. *The American Journal of Human Genetics*, *105*(1), 65-77.

Liu, MZ, Rea-Sandin G, Foerster J, Fritsche L, Brieger K, Clark C, Li K, Pandit A, Zajac G, Abecasis GR, Vrieze S (2020). Validating online measures of cognitive ability in Genes for Good, a genetic study of health and behavior. *Assessment, 27*, 136-148.

#### Generation Scotland: Scottish Family Health Study (Generation Scotland)

Generation Scotland: Scottish Family Health Study (GS:SFHS) is a family-based population cohort dedicated to exploring the genetic and environmental factors that contribute to health and well-being. A total of 23,960 participants from ~7,000 nuclear or extended families were recruited between February 2006 to March 2011 by a cross-disciplinary collaboration of Scottish medical schools and the National Health Service (NHS) in Aberdeen, Dundee, Edinburgh, and Glasgow (Smith et al., 2012). After quality control, there were 555,752 SNPs and 20,027 participants retained. They were aged from 18-99, with a mean age of 46.83 and SD of 15.08. 11,799 (58.92%) of them were female.

We performed GWAS on self-reported personality using two tests. The first was the EPQ-R (Eysenck, 1994) which was used to assess extraversion (N = 19,510) and neuroticism (N = 19,518). A subset of the Generation Scotland cohort (Habota et al., 2021) also took the IPIP (Goldberg, 1992) which measured extraversion (N = 1,101), agreeableness (N = 1,110), conscientiousness (N = 1,104), emotional stability (N = 1,099), and intellect (N = 1,089).

A total of 21,309 participants had data on the EPQ-R and 1,186 on the IPIP, with 1,141 participants having taken both the IPIP and the EPQ-R. Within 1,106 participants who had taken the IPIP and the EPQ-R the correlation between the two extraversion scales was r = 0.782. The correlation between neuroticism from the EPQ-R and emotional stability on the IPIP was r = -0.758 in 1,109 participants.

Amador, C., Huffman, J., Trochet, H., Campbell, A., Porteous, D., Generation Scotland,

Wilson, J.F., Hastie, N., Vitart, V., Hayward, C., Navarro, P., & Haley, C.S. (2015). Recent genomic heritage in Scotland. *BMC Genomics, 16,* 437.

Eysenck H.J., Eysenck S.B.G. (1994). *Manual of the Eysenck Personality Questionnaire: (EPQ-R adult)*. EdITS/Educational and Industrial Testing Service.

Goldberg L.R. (1992). The development of markers for the Big-Five factor structure. *Psychological Assessment*, *4*(1), 26–42.

Habota, T., Sandu, A., Waiter, G.D., McNeil, C.J., Steele, J.D., Macfarlane, J.A., Whalley, H.C., Valentine, R., Younie, D., Crouch, N., Hawkins, E.L., Hirose, Y., Romaniuk, L., Milburn, K., Buchan, G., Coupar, T., Stirling, M., Jagpal, B., MacLennan, B., … McIntosh, A.M. (2019). Cohort profile for the STratifying Resilience and Depression Longitudinally (STRADL) study: A depression-focused investigation of Generation Scotland, using detailed clinical, cognitive, and neuroimaging assessments. *Wellcome Open* Research, *4*, 185.

Nagy, R., Boutin, T.S., Marten, J. Huffman, J.E., Kerr, S.M., Campbell, A., Evenden, L., Gibson, J., Amador, C., Howard, D.M., Navarro, P., Morris, A., Deary, I.J., Hocking, L.J., Padmanabhan, S., Smith, B.H., Joshi, P., Wilson, J.F., Hastie, N.D., … Hayward, C. (2017). Exploration of haplotype research consortium imputation for genome-wide association studies in 20,032 Generation Scotland participants. *Genome Medicine, 9*, 23.

Smith, B.H., Campbell, A., Linksted, P., Fitzpatrick, B., Jackson, C., Kerr, S.M., Deary, I.J., MacIntyre, D.J., Campbell, H., McGilchrist, M., Hocking, L.J., Wisely, L., Ford, I., Lindsay, R.S., Morton, R, Palmer, C.N.A., Dominiczak, A.F., Porteous, D.J, & Morris, A.D. (2012). Cohort Profile: Generation Scotland: Scottish Family Health Study (GS: SFHS). The study, its participants and their potential for genetic research on health and illness. *International Journal of Epidemiology*, *42*(3), 689–700.

#### Helsinki Birth Cohort Study (HBCS)

The Helsinki Birth Cohort Study (HBCS) is composed of 8 760 individuals born between the years 1934-44 in one of the two main maternity hospitals in Helsinki, Finland. Between 2001 and 2003, a randomly selected sample of 928 males and 1075 females participated in a clinical follow-up study with a focus on cardiovascular, metabolic and reproductive health, cognitive function and depressive symptoms. In 2004, various psychological phenotypes were assessed, including the Big Five personality dimensions with the NEO-PI personality inventory. The research plan of the HBCS was approved by the Institutional Review Board of the National Public Health Institute and all participants have signed an informed consent. Detailed information on the selection of the HBCS participants and on the study design can be found elsewhere (Barker et al., 2005; Eriksson et al., 2006; Räikkönen et al., 2008).

HBCS Acknowledgements: We thank all study participants as well as everybody involved in the Helsinki Birth Cohort Study. Helsinki Birth Cohort Study has been supported by grants from the Academy of Finland, the Finnish Diabetes Research Society, Folkhälsan Research Foundation, Novo Nordisk Foundation, Finska Läkaresällskapet, Juho Vainio Foundation, Signe and Ane Gyllenberg Foundation, University of Helsinki, Ministry of Education, Jalmari ja Rauha Ahokas foundation, Emil Aaltonen Foundation, and Yrjö Jahnsson foundation.

Barker, D. J., Osmond, C., Forsén, T. J., Kajantie, E., & Eriksson, J. G. (2005). Trajectories of growth among children who have coronary events as adults. *New England Journal of Medicine*, *353*(17), 1802-1809.
Eriksson, J. G., Osmond, C., Kajantie, E., Forsen, T. J., & Barker, D. J. (2006). Patterns of growth among children who later develop type 2 diabetes or its risk factors. *Diabetologia*, *49*, 2853-2858.
Raikkönen, K., Pesonen, A. K., Heinonen, K., Lahti, J., Kajantie, E., Forsén, T., ... & Eriksson, J. G. (2008). Infant growth and hostility in adult life. *Biopsychosocial Science and Medicine*, *70*(3), 306-313.

**Health and Retirement Study (HRS)**

The Health and Retirement Study (HRS) is a representative national longitudinal study of Americans 50 years old and older and their spouses (Sonnega et al., 2014). Every two years since 2006, HRS participants have completed the 26-adjective Midlife Development Inventory (MDI), which measures the Big Five personality traits with four to seven questions each (Lachman & Weaver, 1997). In 2010, five additional questions were added to the MDI to bolster the measurement of conscientiousness, which we excluded because they prevented aggregation of scores across years and did not markedly improve internal consistency (see Smith et al., 2017). In this study, we restricted genetic analyses to participants who provided personality trait data that were unrelated and of African (N = 1,997) or European (N = 10,079) ancestry (for complete details, see Faul et al., 2014).

Faul, J., Smith, J., & Zhao, W. (2014). Health and retirement study: candidate gene and SNP data description. *Health Retire. Study, Univ. Mich., Ann Arbor, MI.*
Lachman, M.E., & Weaver, S.L. (1997). The Midlife Development Inventory (MIDI) personality scales: Scale construction and scoring. *Waltham, MA: Brandeis University*, *7*, 1-9.
Smith, J., Ryan, L., Fisher, G.G., Sonnega, A., & Weir, D. (2017). HRS psychosocial and lifestyle questionnaire 2006–2016: Documentation report core section LB. *Health and Retirement Study. Retrieved Mar 14*(2022), 202006-2016.
Sonnega, A., Faul, J. D., Ofstedal, M. B., Langa, K. M., Phillips, J. W., & Weir, D. R. (2014). Cohort profile: the health and retirement study (HRS). *International Journal of Epidemiology*, *43*(2), 576-585.

**KFO 256 (KFO)**

Recruitment for this study was done by the KFO 256 project, which is a clinical research unit that is dedicated to investigating the mechanisms of disturbed emotion processing in BPD and is funded by the German Research Foundation (DFG) (Schmahl et al., 2014). Only control subjects were included in the present analyses. Trained clinical psychologists diagnosed BPD according to DSM-IV using the International Personality Disorder Examination (IPDE; Loranger, 1999), and all patients met at least five of the nine DSM-IV criteria for BPD. General exclusion criteria included a history of psychotic or bipolar I disorder, current substance addiction, current pregnancy, organic brain disease, skull or brain damage, severe neurological illness, or use of psychotropic medication at the time of testing. Healthy controls were also excluded if they had any lifetime or current psychiatric diagnoses. The study was conducted in accordance with the Declaration of Helsinki and was approved by the Research Ethics Board of the University of Heidelberg. Prior to study participation, subjects provided written informed consent.

Personality traits were assessed by using an online version of the HEXACO-60 questionnaire (Moshagen et al., 2014). For the present analyses, the scales Openness, Extraversion and Conscientiousness were used. Scales were calculated for subjects with at least 9 out of the 10 respective items. For the present analysis, data were available from 313 central European subjects (69 males, 244 females), with a mean age of 28.12 years (SD = 6.93, range 18-50).

Moshagen, M., Hilbig, B.E., & Zettler, I. (2014). Faktorenstruktur, psychometrische Eigenschaften und Messinvarianz der deutschsprachigen Version des 60-item HEXACO Persönlichkeitsinventars. *Diagnostica*.

Schmahl, C., Herpertz, S.C., Bertsch, K., Ende, G., Flor, H., Kirsch, P., Lis, S., Meyer-Lindenberg, A., Rietschel, M., Schneider, M., Spanagel, R., Treede, R., & Bohus, M. (2014). Mechanisms of disturbed emotion processing and social interaction in borderline personality disorder: state of knowledge and research agenda of the German Clinical Research Unit. *Borderline Personality Disorder and Emotion Dysregulation*, *1*(1), 1-17.

**Lothian Birth Cohorts 1921 and 1936 (LBC1921 & LBC1936)**

The Lothian Birth Cohorts include surviving participants from the Scottish Mental Surveys of 1932 or 1947 (SMS1932 and SMS1947), having been born, respectively in 1921 (LBC1921) and 1936 (LBC1936) (Deary et al, 2004, Deary et al. 2007, Deary et al., 2012, Taylor et al. 2018). The LBC1921 cohort consists of 550 relatively healthy individuals, 316 females and 234 males, assessed on cognitive and medical traits at about 79 years of age. When tested, the sample had a mean age of 79.1 years (SD = 0.6). The LBC1936 consists of 1,091 relatively healthy individuals assessed on cognitive and medical traits at about 70 years of age. At baseline the sample of 548 men and 543 women had a mean age 69.6 years (SD = 0.8). They were all Caucasian and almost all lived independently in the Lothian region (Edinburgh city and surrounding area) of Scotland. In LBC1936 personality was assessed using the 60-item NEO-FFI (Costa & McCrae, 1992), and in LBC1921 personality was assessed using the 50-item IPIP (Goldberg, 1999). Quality control measures were applied; 517 and 1005 participants remained for LBC1921 and LBC1936 respectively. Among participants with genome-wide data, 425 (LBC1921) and 880 (LBC1936) individuals were available for the present analysis.

Costa, P.T., & McCrae, R.R. (1992). Professional manual for the NEO-PI-R and NEO-FFI. Odessa, FL: Psychological Assessment Resources.

Deary I.J., Whiteman M.C., Starr J.M., Whalley L.J., Fox H.C. (2004). The impact of childhood intelligence on later life: following up the Scottish Mental Surveys of 1932 and 1947. *Journal of Personality and Social Psychology*, *86*, 130-147.

Deary I.J., Gow A.J., Taylor M.D., Corley J., Brett C., Wilson V., Campbell, H., Whalley, L.J., Visscher, P.M., Porteous, D.J., & Starr, J.M. (2007). The Lothian Birth Cohort 1936: a study to examine influences on cognitive ageing from age 11 to age 70 and beyond. *BMC Geriatrics, 7*, 28.

Deary I.J., Gow A.J., Pattie A., Starr J.M. (2012). Cohort profile: The Lothian Birth Cohorts of 1921 and 1936. I*nternational Journal of Epidemiology*, *41*(6), 1576-1584.
Goldberg, L.R. (1999). A broad-bandwidth, public domain, personality inventory measuring the lower-level facets of several five-factor models. In I. Mervielde, I.J. Deary, F. De Fruyt, & F. Ostendorf (Eds.). *Personality Psychology in Europe*, Vol. 7, Tilburg University Press, Tilburg.
Taylor, A.M., Pattie, A., Deary, I.J. (2018). Cohort profile update: the Lothian Birth Cohorts of 1921 and 1936. *International journal of epidemiology*, *47*(4), 1042-1042r.

#### Lifelines

Lifelines is a multidisciplinary prospective population-based cohort study examining in a unique three-generation design the health and health-related behaviors of 167,729 persons living in the North of the Netherlands (Scholtens et al., 2015). It employs a broad range of investigative procedures in assessing the biomedical, socio-demographic, behavioral, physical and psychological factors which contribute to the health and disease of the general population, with a special focus on multi-morbidity and complex genetics. Recruitment and data collection have been described elsewhere (Scholtens et al., 2015). In short, participants were recruited through their general practitioners or family members. Additionally, self-registration online was also possible. For the present analyses, we used data on personality from the baseline measurements carried out between 2007 and 2013. Personality was assessed using two abbreviated versions of the revised Neuroticism-Extroversion-Openness (NEO) Personality Inventory (NEO-PI-R, Costa et al., 1992; Hoekstra et al., 2007) that cover three domains of personality: neuroticism, extraversion, and conscientiousness. Therefore, we performed GWAS on these self-reports as our phenotypes.

Costa, P.T., & McCrae, R.R. (1992). Professional manual for the NEO-PI-R and NEO-FFI. Odessa, FL: Psychological Assessment Resources.
Hoekstra, H., Ormel, J., & De Fruyt, F. (2007). *NEO-PI-R Handleiding*. Hogrefe Uitgevers BV.
Scholtens, S., Smidt, N., Swertz, M.A., Bakker, S.J.L., Dotinga, A., Vonk, J.M., van Dijk, F., van Zon, S.K.R., Wijmenga, C., Wolffenbuttel, B.H.R., & Stolk, R.P. (2015). Cohort Profile: LifeLines, a three-generation cohort study and biobank. *International Journal of Epidemiology*, *44*(4), 1172–1180.

#### Mannheim Blood Donor Study

The Mannheim Blood Donor Study is a sample collected at blood donation events in Baden-Württemberg, Germany. The study was approved by the Ethics Committee of the Medical Faculty of the University of Heidelberg and written informed consent was obtained from all subjects. This cohort was a sample of the general population for which blood donation exclusion criteria applied: bodyweight < 50 kg, current general diseases, and high risk of infection. Subjects were given a questionnaire to assess demographic information and information about their mental and physical health at the blood donation event, and these questionnaires were then returned to the study center by mail. The 12-item Neuroticism scale of the NEO-Five-Factor-Inventory (NEO-FFI) was applied. For the present analysis, data were available from 1,444 central European subjects (662 males and 782 females), with a mean age of 44.74 years (SD = 13.03, range 18-72).

Nenadić, I., Meller, T., Schmitt, S., Stein, F., Brosch, K., Mosebach, J., Ettinger, U., Grant, P., Meinert, S., Opel, N., Lemke, H., Fingas, S., Förster, K., Hahn, T., Jansen, A., Andlauer, T.F.M., Forstner, A.J., Heilmann-Heimbach, S., Hall, A.S.M., … Kircher, T. (2022). Polygenic risk for schizophrenia and schizotypal traits in non-clinical subjects. *Psychological Medicine*, *52*(6), 1069-1079.

Streit, F., Witt, S.H., Awasthi, S., Foo, J.C., Jungkunz, M., Frank, J., Colodro-Conde, L., Hindley, G., Smeland, O., Maslahati, T., Schwarze, C., Schott, B., Hartmann, A., Giegling, I., Zillich, L., Sirignano, L., Poisel, E., Chen, C., Nöthen, M., ... Andreassen, O.A. (2022). Borderline personality disorder and the big five: molecular genetic analyses indicate shared genetic architecture with neuroticism and openness. *Translational Psychiatry*, *12*(1), 1-8.

Witt, S.H., Streit, F., Jungkunz, M., Frank, J., Awasthi, S., Reinbold, C.S., Treutlein, J., Degenhardt, F., Forstner, A.J., Heilmann-Heimbach, S., Dietl, L., Schwarze, C.E., Schende, D.I., Strohmaier, J., Bethell, A., Craddock, N., Di Florio, A., Forty, E., Fraser, C., ... Lieb, K. (2017). Genome-wide association study of borderline personality disorder reveals genetic overlap with bipolar disorder, major depression and schizophrenia. *Translational Psychiatry*, *7*(6), e1155-e1155.

#### Minnesota Center for Twin & Family Research (MCTFR)

The MCTFR GWAS sample represents participants from three longitudinal studies: The Minnesota Twin Family Study (MTFS), the Sibling Interaction and Behavior Study (SIBS), and the Enrichment Study (ES). These studies share similar assessment protocols and a common sampling unit, a four-member family consisting of sibling pairs and their rearing parents. Offspring in all three samples were initially assessed in adolescence and followed into at least early adulthood.

Genome-wide genotyping was carried out using the Illumina Human660W-Quad array (Miller et al., 2012). Untyped genotypes were imputed to the Haplotype Reference Consortium through the Michigan Imputation Server, using Minimac3.

Phenotypes studied were neuroticism and extraversion created by a harmonization of items in the MPQ personality test (van den Berg et al., 2014). There were 3824 individuals with phenotypes in the main analysis and 942 siblings with the phenotypes for the within-sibling analysis. Age of the main sample ranged from 12-64 (M=40, SD=11; 51% female).

MCTFR recruitment, assessment and genotyping was supported in part by USPHS Grants from the National Institute on Alcohol Abuse and Alcoholism (AA09367 and AA11886), the National Institute on Drug Abuse (DA05147, DA13240, and DA024417), and the National Institute on Mental Health (MH066140).

Miller, M. B., Basu, S., Cunningham, J., Eskin, E., Malone, S. M., Oetting, W. S., Schork, N., Sul, J. H., Iacono, W. G., & McGue, M. (2012). The Minnesota Center for Twin and Family Research genome-wide association study. *Twin research and human genetics*, 15(6), 767–774.

van den Berg, S.M. et al. (2014) “Harmonization of neuroticism and extraversion phenotypes across inventories and cohorts in the genetics of Personality Consortium: An application of item response theory," *Behavior genetics*, 44(4), pp. 295–313.

#### Midlife in the United States (MIDUS)

Midlife in the United States (MIDUS) is a national longitudinal study of health and well-being (http://midus.wisc.edu/). It was conceived by a multidisciplinary team of scholars interested in understanding aging as an integrated bio-psycho-social process, and as such it includes data collected in a wide array of research protocols using a variety of survey and non-survey instruments (Radler, 2014). MIDUS participants rated themselves on the Midlife Development Inventory (MIDI; Lachman & Weaver, 1997) to index the Big Five. There were 2,117 participants with genome-wide data aged 25-87 (M = 53.8, SD = 12.57; 54% female) that were available for the present analyses. N = 1,298 aged 25-83 (M = 54.2, SD = 12.42; 51% female) contributing to European ancestry analyses and N = 316 aged 25-87 (M = 51.1, SD = 11.89; 68% female) contributing to African ancestry analyses. Since 1995 the MIDUS study has been funded by the following: John D. and Catherine T. MacArthur Foundation Research Network, National Institute on Aging (P01-AG020166), National institute on Aging (U19-AG051426). Biomarker data collection was further supported by the NIH National Center for Advancing Translational Sciences (NCATS) Clinical and Translational Science Award (CTSA) program as follows: UL1TR001409 (Georgetown), UL1TR001881 (UCLA), 1UL1RR025011 (UW).

Lachman, M.E., & Weaver, S.L. (1997). The Midlife Development Inventory (MIDI) personality scales: Scale construction and scoring. *Waltham, MA: Brandeis University*, *7*, 1-9.

Radler, B.T. (2014). The Midlife in the United States (MIDUS) Series: A National Longitudinal Study of Health and Well-being. *Open Health Data*, *2*(1).

#### The Norwegian Mother, Father, and Child Cohort Study (MoBa)

MoBa is a population-based pregnancy cohort study conducted by the Norwegian Institute of Public Health (Magnus et al., 2016). Participants were recruited from all over Norway from 1999-2008. The women consented to participation in 41% of the pregnancies. The cohort now includes 114,500 children, 95,200 mothers, and 75,200 fathers. The current study is based on version 12 of the quality-assured data files. The Big-5 personality traits were assessed among fathers at recruitment in the 15th week of pregnancy and among mothers when the child was 5 years old by using The International Personality Item Pool (IPIP) Big-Five Factor Markers (Goldberg, 1999). The IPIP is self-reported and consists of 10 items for each of the Big-5 personality traits. We allowed up to two of the ten items to be missing among participants and calculated an individual's score on a trait as the average of the valid items. Among participants with genome-wide data, 42,436 individuals aged 30-85 (M = 47.32, SD = 5.07; 47% female) were available for the present analyses (Corfield et al., 2022).

We are grateful to all the participating families in Norway who take part in this on-going cohort study. We thank the Norwegian Institute of Public Health (NIPH) for generating high-quality genomic data. The MoBa analyses was performed on the Tjeneste for Sensitive Data (TSD) facilities, owned by the University of Oslo, operated and developed by the TSD service group at the University of Oslo, IT-Department (USIT), using resources provided by Sigma2—the National Infrastructure for High Performance Computing and Data Storage in Norway (UNINETT).

Corfield, E. C., Shadrin, A. A., Frei, O., Rahman, Z., Lin, A., Athanasiu, L., ... & Havdahl, A. (2022). The Norwegian Mother, Father, and Child cohort study (MoBa) genotyping data resource: MoBaPsychGen pipeline v. 1. *BioRxiv*, 2022-06.

Goldberg, L.R. (1999). A broad-bandwidth, public domain, personality inventory measuring the lower-level facets of several five-factor models. In I. Mervielde, I.J. Deary, F. De Fruyt, & F. Ostendorf (Eds.). *Personality Psychology in Europe*, Vol. 7, Tilburg University Press, Tilburg.

Magnus P., Birke, C., Vejrup, K., Haugan, A., Alasker, E., Daltveit, A.K., Handal, M., Haugen, M., Høiseth, G., Knudsen, G.P., Paltiel, L., Schreuder, P., Tambs, K, Vold, L., & Stoltenberg, C. (2016). Cohort Profile Update: The Norwegian Mother and Child Cohort Study (MoBa). I*nternational Journal of Epidemiology*, *45*(2), 382-388.

#### Million Veterans Program (MVP)

The VA's Million Veteran Program (MVP) is a national research program looking at how genes, lifestyle, military experiences, and exposures affect health and wellness in veterans. MVP participants completed the BFI-10 as part of a self-report lifestyle survey, with two items for each of the Big Five personality traits. Initial MVP analyses of these data are described in full in Gupta and colleagues (2024), and genotyping and imputation in this sample is described in full in Gaziano and colleagues (2016) and Levey and colleagues (2020). For this study, the BFI-10 GWAS was re-estimated using controls for age, age-squared, sex, 10 ancestral principal components, and birth cohort, to ensure consistency with other contributed cohort GWAS. The award number that funded the VA analysis was 5IK2BX005058

Gaziano, J. M., Concato, J., Brophy, M., Fiore, L., Pyarajan, S., Breeling, J., ... & O'Leary, T. J. (2016). Million Veteran Program: A mega-biobank to study genetic influences on health and disease. *Journal of clinical epidemiology*, *70*, 214-223.

Gupta, P., Galimberti, M., Liu, Y., Beck, S., Wingo, A., Wingo, T., ... & Levey, D. F. (2024). A genome-wide investigation into the underlying genetic architecture of personality traits and overlap with psychopathology. *Nature Human Behaviour*, 1-15.

Levey, D. F., Gelernter, J., Polimanti, R., Zhou, H., Cheng, Z., Aslan, M., ... & Stein, M. B. (2020). Reproducible genetic risk loci for anxiety: results from∼ 200,000 participants in the Million Veteran Program. *American Journal of Psychiatry*, *177*(3), 223-232.

#### Neuronale Korrelate von Wissen, Intelligenz, Persönlichkeit, Emotionaler Kompetenz und Motivation (NKWIPEM)

The Neural correlates of knowledge, intelligence, personality, emotional competence, and motivation (NKWIPEM) project was designed to investigate the neural correlates of various psychological variables, including general knowledge, intelligence, and Big Five personality traits (Genç et al., 2018; Schlüter et al., 2022). Data acquisition started in 2014 and ended in 2019. Subjects were recruited via advertisements posted online or flyers placed on campus at

Ruhr University Bochum. The final sample includes 520 (265 male) healthy individuals (no history of psychological or neurological disorders) aged 18-75 (M = 27.15, SD = 9.10). On their first day of participation, subjects underwent MRI scanning at Bergmannsheil hospital (Bochum, Germany). On their second to fourth day of participation, subjects completed several pen-and-paper tests and questionnaires at Ruhr University Bochum. Personality traits were assessed via the German version of the Revised NEO-Personality-Inventory (NEO-PI-R), which comprises a total of 240 self-report items to measure the domains extraversion, agreeableness, conscientiousness, neuroticism, and openness to experience as well as six more specific facets of each domain (Ostendorf & Angleitner, 2004). We perform GWAS on the standardized domain scores as our primary phenotypes.

Genç, E., Fraenz, C., Schlüter, C., Friedrich, P., Hossiep, R., Voelkle, M.C., Ling, J.M., Güntürkün, O., & Jung, R.E. (2018). Diffusion markers of dendritic density and arborization in gray matter predict differences in intelligence. *Nature Communications*, *9*(1), 1905.

Ostendorf, F. & Angleitner, A. (2004). *NEO-Persönlichkeitsinventar nach Costa und McCrae, Revidierte Fassung (NEO-PI-R)*: Hogrefe Göttingen.

Schlüter, C., Fraenz, C., Friedrich, P., Güntürkün, O., & Genç, E. (2022). Neurite density imaging in amygdala nuclei reveals interindividual differences in neuroticism. *Human Brain Mapping*, *43*(6), 2051-2063.

#### Netherlands Twin Register (NTR)

The Netherlands Twin Register (NTR) is an ongoing research initiative collecting data in twins and their extended family members. Since 1987, the Netherlands Twin Register (NTR) database has been accumulating information on twins and triplets. This registry is initiated when the parents of newborn twins voluntarily register them. For twins under the age of 14, data are collected from their parents and teachers. At 14 years old, the twins themselves contribute data through self-reports. Ethical clearance for this study has been granted by the Central Ethics Committee on Research Involving Human Subjects at the VU University Medical Centre in Amsterdam, which is an Institutional Review Board certified by the U.S. Office of Human Research Protections. The approval carries the IRB number IRB-2991 under Federal-wide Assurance-3703 and includes specific institute codes (94/105, 96/205, 99/068, 2003/182, 2010/359). All participants have given their informed consent to be part of this registry. Detailed information on data collection in the NTR can be found elsewhere (Ligthart et al., 2019).

Here, we used survey waves 7, 9 and 10 and the DHBQ16 that all employed the NEO-FFI-3 version of the NEO Five-Factor Inventory (Costa & McCrae, 1992; Hoekstra, Ormel, & De Fruyt, 1996), which contains 12 questions for each personality trait. Item scores were recoded such that a higher score reflected a higher trait-score. Only surveys completed when participants were between 18 and 65 years (inclusive) old were included. If multiple surveys were available in any given participant, the survey with the lowest amount of missing values for the five personality traits was selected. If multiples surveys meeting these criteria remained, the survey where the age of the participant at baseline was closest to 18 was selected.

Hoekstra, H. A., Ormel, J., & De Fruyt, F. (1996). Manual of the Dutch version of the NEO-PI-R/NEO-FFI.

Ligthart, L., van Beijsterveldt, C. E., Kevenaar, S. T., de Zeeuw, E., van Bergen, E., Bruins, S., ... & Boomsma, D. I. (2019). The Netherlands twin register: Longitudinal research based on twin and twin-family designs. *Twin research and human genetics*, 22(6), 623-636.

#### Pre-, Peri-, and Postnatal Stress: Epigenetic Impact on Depression (POSEIDON)

The Pre-, Peri-, and Postnatal Stress: Epigenetic impact on Depression (POSEIDON) study is a birth cohort collected in the Rhein-Neckar region of Germany. The study was approved by the Ethics Committee of the Medical Faculty of the University of Heidelberg and written informed consent was obtained from all subjects. In POSEIDON, data was collected during the third trimester of pregnancy (T1), in the days after childbirth (T2), and six (T3) and 45 months (T4) postpartum. To replace dropouts, new parents were included in the study at T4. In 2020, a reassessment with online questionnaires was conducted (T5). Inclusion criteria of the mothers: age 16-45 years, German speaking, main caregiver of the child. Exclusion criteria of the newborn children: born at 30 weeks of pregnancy or earlier, birthweight < 1500 grams.

Big Five were assessed at different time points for subsets of the cohort. The mothers of the POSEIDON cohort completed the NEO-FFI-30 at T1, the fathers at T3; the mothers and fathers of the newly recruited subsample at T4. For the present analysis, data were available from 693 central European subjects (320 males, 373 females), with a mean age of 33.75 years (SD = 5.82, range 18-55).

Send, T.S., Gilles, M., Codd, V., Wolf, I., Bardtke, S., Streit, F., Strohmaier, J., Frank, J., Schendel, D., Sütterlin, M.W., Denniff, M., Laucht, M., Samarni, N.J., Deuschle, M, Rietschel, M., & Witt, S. H. (2017). Telomere length in newborns is related to maternal stress during pregnancy. *Neuropsychopharmacology*, *42*(12), 2407-2413.

Send, T.S., Bardtke, S., Gilles, M., Wolf, I.A.C., Sütterlin, M.W., Kirschbaum, C., Laucht, M., Witt, S.H., Rietschel, M., Streit, F., & Deuschle, M. (2019). Stress reactivity in preschool-aged children: Evaluation of a social stress paradigm and investigation of the impact of prenatal maternal stress. *Psychoneuroendocrinology*, *101*, 223-231.

#### QIMR (QIMR_1-QIMR_4)

The QIMR cohort is made up of the following datasets, containing overlapping participants:

The Prospective Imaging Study of Aging: Genes, Brain and Behaviour (PISA): Participants of the online component of the PISA study (Lupton 2021) were invited to complete an online survey providing a global assessment of lifestyle, cognitive, and behavioural function. Data for personality traits were collected using the NEO Five-Factor Inventory-3 (NEO-FFI-3). PISA was funded by a National Health and Medical Research Council (NHMRC) Boosting Dementia Research Initiative Team Grant (APP1095227).

The Brisbane Adolescent Twin Study: Adolescent twins and siblings previously participated in studies identifying genetic polymorphisms associated with moliness and cognitive function, as well as several other projects. 950 families were invited to participate in a mail and phone study assessing reading ability, taste and smell sensitivity, and health and wellbeing (Wright, 2004). This included an assessment of personality using the Revised NEO-Personality Inventory (NEO-PI-R).

The Borderline Personality Disorder Study: Twins aged 18 years and older were recruited from the Australian Twin Registry (ATR) and previous studies conducted at QIMR. All participants were invited to complete an online or paper-based questionnaire collecting data related to Borderline Personality Disorder (BPD) and the personality dimensions that underlie the symptoms of this disorder. This included the revised NEO-Personality Inventory (NEO-PI-R).

Memory, Attention and Problem Solving in Adolescent Twins (MAPS): Twin study measuring key components of cognition in order to identify genes influencing mental functioning. Participants aged 16 and older were recruited primarily through South East Queensland primary and secondary schools (Wright, 2001). Personality was assessed using the Neo Five-Factor Inventory (NEO-FFI and NEO-FFI-3).

Twenty-Five and Up (25Up) Study: A large cohort of Australian Twins that have been studied longitudinally from the age of 12. Data were collected between 2016 and 2018 from 2,540 twins and their non-twin siblings on mental health disorders, general demographic information, IQ and environment (Mitchell, 2019). Personality traits were measured using the NEO Ten-Item Personality Inventory (NEO-TIPI).

In this study, analyses were stratified by measurement instrument, such that participants who completed the NEO-FFI were considered "QIMR_1," participants who completed the NEO-FFI 3 were considered "QIMR_2," participants who completed the NEO-PI-R were considered "QIMR_3," and participants who completed the TIPI were considered "QIMR_4." Overlapping responses were pruned so that participants solely contributed to one questionnaire's GWAS (See **Table S1** for complete details).


Lupton, M.K., Robinson, G.A., Adam, R.J., Rose, S., Byrne, G.J., Salvado, O., Pachana, N.A., Almeida, O.P., McAloney, K., Gordon, S.D., Raniga, P., Fazlollahi, A., Xia, Y., Ceslis, A., Sonkusare, S., Zhang, Q., Kholghi, M., Karunanithi, M., Mosley, P.E., ... Breakspear, M.A. (2021). A prospective cohort study of prodromal Alzheimer's disease: Prospective Imaging Study of Ageing: Genes, Brain and Behaviour (PISA). *Neuroimage: Clinical*, *29*, 102527.

Wright, M.J., & Martin, N.G. (2004). Brisbane Adolescent Twin Study: Outline of study methods and research projects. *Australian Journal of Psychology*, *56*,65-78.

Wright, M., De Geus, E., Ando, J., Luciano, M., Posthuma, D., Ono, Y., Hansell, N., Van Baal, C., Hiraishi, K., Hasegawa, T., Smith, G., Geffen, G., Geffen, L., Kanba, S., Miyake, A., Martin, N., & Boomsma, D. (2001). Genetics of cognition: outline of a collaborative twin study. *Twin Research and Human Genetics*, *4*(1), 48-56.

Mitchell, B.L., Campos, A.I., Rentería, M.E., Parker, R., Sullivan, L., McAloney, K., Couvy-Duchesne, B., Medland, S.E., Gillespie, N.A., Scott, J., Zietsch, B.P., Lind, P.A., Martin, N.G., & Hickie, I.B. (2019). Twenty-Five and Up (25Up) Study: A New Wave of the Brisbane Longitudinal Twin Study. *Twin Research and Human Genetics*, *22*(3), 154-163.


#### Spit for Science (S4S)

Spit for Science (S4S) VCU Student Survey is an effort led by researchers at VCU to create a university-wide research opportunity for VCU students. The scientific focus of the project is to understand why some people are more likely than others to develop problems associated with the use of alcohol and other substances, and difficulties with emotional health. The project aims to understand how individual predispositions come together with environmental factors to contribute to these outcomes. 21.3% of the sample is of African ancestry, 12.5% is American, 9.7% is East Asian, 47.9% is European and 8.6% is South Asian. Participants rated themselves using shortened 15-item Big Five inventory, which measures extraversion, agreeableness, conscientiousness, neuroticism, and openness to experience. Among participants with genome-wide data, 7,676 individuals aged 17.8-32.1 (M = 18.6, SD = 0.6; 62% female) were available for the present analyses, with N = 1637 and 3849 participants contributing to African and European ancestry analyses, correspondently.

#### SardiNIA

The SardiNIA longitudinal project is an ongoing study of about 8,000 general population participants enrolled in four small towns of the central east coast of Sardinia, Italy (Orrù et al. 2013, Steri et al. 2017). The study captures 5 phases of sample and data collection and investigates the genetics and epidemiology of complex traits/phenotypes, including immune, cardiometabolic risk factors, a variety of biometrical parameters and personality and cognitive features. Among participants with genome-wide data, 5,697 individuals from 1,102 families aged 14-94 (M = 42.83, SD = 17.1, male 2407; female 3290; 57.75% female) were available for the present analyses, with all participants with European ancestry. All SardiNIA participants completed once the Italian version (Terracciano, 2003) of the Revised NEO Personality Inventory (NEO-PI R)(Costa & McCrae). The questionnaire consists of 240 items answered on a 5-point Likert scale, from "strongly disagree" to "strongly agree". Data across languages and cultures support the reliability and validity of the NEO-PI R ([McCrae et al., 2005). Indeed, in the SardiNIA cohort, the factor structure showed high congruence with the normative structure (Tucker phi ≥ 0.91) and high internal consistency (Cronbach alpha ≥ .80) for the five factors (Costa et al., 2007).

Costa, P.T., & McCrae, R.R. (1992). Professional manual for the NEO-PI-R and NEO-FFI. Odessa, FL: Psychological Assessment Resources.

Costa, P.T., Terracciano, A., Uda, M., Vacca, L., Mameli, C., Pilia, G., Zonderman, A.B., Lakatta, E., Schlessinger, D., & McCrae, R.R. (2007). Personality traits in Sardinia: Testing founder population effects on trait means and variances. *Behavior Genetics*, *37*, 376-387

McCrae, R.R., & Terracciano, A. (2005). Universal features of personality traits from the observer's perspective: Data from 50 cultures. *Journal of Personality and Social Psychology*, *88*, 547-561.

Orrù, V., Steri, M., Sole, G., Sidore, C., Virdis, F., Dei, M., Lai, S., Zoledziewska, M., Busonero, F., Mulas, A., Floris, M., Mentzen, W.I., Urru, S.A.M., Olla, S., Marongiu, M., Piras, M.G., Lobina, M., Maschio, A., Pitzalis, M., … Cucca, F. (2013). Genetic variants regulating immune cell levels in health and disease. *Cell*, *155*, 242–256.

Steri, M., Orrù, V., Idda, M.L., Pitzalis, M., Pala, M., Zara, I., Sidore, C., Faà, V., Floris, M., Deiana, M., Asunis, I., Porcu, E., Mulas, A., Piras, M.G., Lobina, M., Lai, S., Marongiu, M., Serra, V., Marongiu, M., … Cucca, F. (2017). Overexpression of the cytokine BAFF and autoimmunity risk. *New England Journal of Medicine*, *376*, 1615–1626.

Terracciano, A. (2003). The Italian version of the NEO PI-R: conceptual and empirical support for the use of targeted rotation. *Personality and Individual Differences*, *35*, 1859-1872.

#### The STAGE Cohort of the Swedish Twin Registry

Personality data in the Swedish STAGE cohort, which includes all adult twins born between 1959 and 1985 registered with the Swedish Twin Registry (Lichtenstein et al., 2022; Lichtenstein et al., 2006; Magnusson et al., 2013), were collected online in 2012 and 2013 as part of a larger study on music-related traits. In total around 9,800 twins filled in a Swedish translation (Zakrisson, 2010) of the 44-item Big Five Inventory (BFI; John et al., 2008) assessing extraversion, agreeableness, conscientiousness, neuroticism, and openness to experience. Responses were given on a five-point Likert scale, ranging from 'do not agree at all' to 'agree completely'. Reliability and validity of the Swedish version has been found to be similar to previously reported estimates for personality (Zakrisson, 2010). Among participants with BFI data, N = 5,213 aged between 27 and 54 (M = 40.9, SD = 7.80, 61.7% female) from 4,220 families were genotyped and available for the present analyses. The study was approved by the Regional Ethics Review Board in Stockholm (Dnrs 2011/570-31/5, 2011/1425-31, 2012/1107/32 and 2019-05879).

The Swedish Twin Registry is managed by Karolinska Institutet and receives funding through the Swedish Research Council under the grant no 2017-00641. The computations and data handling was enabled by resources provided by the National Academic Infrastructure for Supercomputing in Sweden (NAISS) and the Swedish National Infrastructure for Computing (SNIC) at Uppsala Multidisciplinary Center for Advanced Computational Science (UPPMAX) partially funded by the Swedish Research Council through grant agreements no. 2022-06725 and no. 2018-05973.v

John, O. P., Naumann, L. P., & Soto, C. J. (2008). Paradigm shift to the integrative big five trait taxonomy. *Handbook of Personality: Theory and Research*, *3*(2), 114-158.

Lichtenstein, P., De Faire, U., Floderus, B., Svartengren, M., Svedberg, P., & Pedersen, N. L. (2002). The Swedish Twin Registry: a unique resource for clinical, epidemiological and genetic studies. *Journal of Internal Medicine*, *252*(3), 184-205.

Lichtenstein, P., Sullivan, P. F., Cnattingius, S., Gatz, M., Johansson, S., Carlström, E., Björk, C., Svartengren, M., Wolk, A., Klareskog, L., de Faire, Ul, Schalling, M., Palmgren, J., & Pedersen, N. L. (2006). The Swedish Twin Registry in the third millennium: an update. *Twin Research and Human Genetics*, *9*(6), 875-882.

Magnusson, P. K., Almqvist, C., Rahman, I., Ganna, A., Viktorin, A., Walum, H., Halldner, L., Lundström, S., Ullén, F., Långström, N., Larsson, H., Nyman, A., Hellner Gumpert, C., Råstam, M., Anckarsäter, H., Cnattingius, S. Johannesson, M., Ingelsson, E., Klareskog., ... Lichtenstein, P. (2013). The Swedish Twin Registry: establishment of a biobank and other recent developments. *Twin Research and Human Genetics*, *16*(1), 317-329.

Zakrisson, I. (2010). Big Five Inventory (BFI): Utprövning för svenska förhållanden. Mid Sweden University.

#### Twins Early Development Study (TEDS)

The Twins Early Development Study (TEDS), one of the largest twin cohorts in the world, investigates how genetic and environmental factors shape individual differences in cognitive abilities, educational achievement, behavior, and emotions in the context of typical development. All twins born in England and Wales between 1994 and 1996 were invited to take part in the study, with over 13,000 families participating in the first wave of data collection when twins were around 18 months old. The sample remains fairly representative of the population in England and Wales in terms of ethnicity and family socioeconomic factors. 92.9% of the full sample is white, comparable with the national equivalents of 93% for cohorts of parents with young children born in late 1990s – early 2000s. More details about the representativeness of the TEDS sample in terms of family socioeconomic factors is available in Rimfeld et al. (2019).

TEDS twins rated themselves on the 30-item Big Five personality test (Mullins-Sweatt et al., 2006), which has 6 items for each trait, at ages 16 and 21. We performed GWAS on the aggregated scores across the two measurement occasions. Overall, 41.1% of the available sample were male. For PGI analyses, 6,362 participants were included in population-level PGI analyses, and 4,751 participants were included in within-family PGI analyses.

Mullins-Sweatt, S.N., Jamerson, J.E., Samuel, D.B., Olson, D.R., & Widiger, T.A. (2006). Psychometric properties of an abbreviated instrument of the five-factor model. *Assessment*, *13*(2), 119-137.

Rimfeld, K., Malanchini, M., Spargo, T., Spickernell, G., Selzam, S., McMillan, A., Dale, P.S., Eley, T.C., & Plomin, R. (2019). Twins early development study: A genetically sensitive investigation into behavioral and cognitive development from infancy to emerging adulthood. *Twin Research and Human Genetics*, *22*(6), 508-513.

##### Tracking Adolescents' Individual Lives Survey Parent Sample (TRAILS)

TRAILS (TRacking Adolescents' Individual Lives Survey; https://www.trails.nl/en/home) is an ongoing prospective cohort study of Dutch adolescents and young adults, with bi- or triennial measurements from age 11 onwards, which started in 2001 (Huisman et al., 2008; Oldehinkel et al., 2015). TRAILS consists of a general population and a clinical cohort (TRAILS-CC), both from the North of the Netherlands. The study was approved by the Dutch Central Committee on Research Involving Human Subjects. At the third study wave, both the adolescents and their parents provided information about their personality. For the present purposes, we only used parental data, because the data of parents and children are not independent. Personality data was available for approximately 870 fathers (mean age 47.7, SD 5.0, range 33-74 years) and 1,090 mothers (mean age 44.9, SD 4.7, range 32-60 years), all of European ancestry. Based on several facets of the NEO-PI-R (Costa & McCrae, 1992), we constructed scores for neuroticism, extraversion, and conscientiousness. More specifically, the Neuroticism score was calculated by averaging the facet scores on Angry Hostility, Self-consciousness, Impulsivity, and Vulnerability; the Extraversion score by averaging the scores on Assertiveness, Gregariousness, Activity, and Excitement seeking; and the Conscientiousness score by averaging the scores on Competence, Achievement Striving, Self-discipline, and Deliberation.

TRAILS is a collaborative project involving various departments of the University Medical Center and University of Groningen, the University of Utrecht, and the Parnassia Psychiatric Institute, all in the Netherlands. TRAILS has been financially supported by grants from the Netherlands Organization for Scientific Research NWO (Medical Research Council program grant GB-MW 940-38-011; ZonMW Brainpower grant 100-001-004; ZonMw Risk Behavior and Dependence grant 60-60600-97-118; ZonMw Culture and Health grant 261-98-710; Social Sciences Council medium-sized investment grants GB-MaGW 480-01-006 and GB-MaGW 480-07-001; Social Sciences Council project grants GB-MaGW 452-04-314 and GB-MaGW 452-06-004; ZonMw Longitudinal Cohort Research on Early Detection and Treatment in Mental Health Care grant 636340002, NWO large-sized investment grant 175.010.2003.005; NWO Longitudinal Survey and Panel Funding 481-08-013 and 481-11-001; NWO Vici 016.130.002, 453-16-007/2735, and Vi.C 191.021; NOW Gravitation 024.001.003), the Dutch Ministry of Justice (WODC), the European Science Foundation (EuroSTRESS project FP-006), the European Research Council (ERC-2017-STG-757364 en ERC-CoG-2015-681466), Biobanking and Biomolecular Resources Research Infrastructure BBMRI-NL (CP 32), The Gratema foundation, the Jan Dekker foundation, the participating universities, and Accare Centre for Child and Adolescent Psychiatry. Statistical analyses were carried out on the Genetic Cluster Computer (http://www.geneticcluster.org), which is financially supported by the Netherlands Scientific Organization (NWO 480-05-003) along with a supplement from the Dutch Brain Foundation.

Costa, P.T., & McCrae, R.R. (1992). Professional manual for the NEO-PI-R and NEO-FFI. Odessa, FL: Psychological Assessment Resources.

Huisman, M., Oldehinkel, A.J., De Winter, A.F., Minderaa, R.B., De Bildt, A., Huizink, A.C., Verhulst, F.C., & Ormel, J. (2008). Cohort profile: The Dutch "TRacking Adolescents' Individual Lives' Survey"; TRAILS. *International Journal of Epidemiology*, *37*(6), 1227-35.

Oldehinkel, A.J., Rosmalen, J.G.M., Buitelaar, J.K., Hoek, H.W., Ormel, J., Raven, D., Reijneveld, S.A., Veenstra, R., Verhulst, F.C., Vollebergh, W.A.M., & Hartman, C.A. (2015). Cohort Profile update. The TRacking Adolescents' Individual Lives Survey (TRAILS). *International Journal of Epidemiology*, *44*(1), 76-76n.

#### Texas Twin Project (TTP)

The Texas Twin Project (TTP) is a study of school-age twins ($3^{rd}$ through $12^{th}$ grade) enrolled in public schools in the Austin, Texas and Houston, Texas metropolitan areas (Harden et al., 2013), designed to maximize representation of low socioeconomic status families and racial/ethnic minorities. School rosters were used to identify twin families from a target population with sizable populations of African-American (18%), Hispanic / Latino (48%), and non-Hispanic White (27%) children and adolescents, over half of whom meet U.S. guidelines for classification as economically disadvantaged. TTP participants rated themselves on up to four occasions using a child version of the 44-item Big Five Inventory, which measures Extraversion, Agreeableness, Conscientiousness, Neuroticism, and Openness to Experience (John, Naumann, & Soto, 2008), as described in further detail in Mõttus et al. (2019). We performed GWAS on the 662 individuals from 355 families (324 males and 338 females, *M*age = 13.18, age range 7-20) who provided personality data, were of European ancestry, and had been genotyped.

Harden, K.P., Tucker-Drob, E.M., & Tackett, J.L. (2013). The Texas twin project. *Twin Research and Human Genetics*, *16*(1), 385-390.

John, O.P., Naumann, L.P., & Soto, C.J. (2008). Paradigm shift to the integrative big five-trait taxonomy: History, measurement, and conceptual issues. In O. P. John, R. W. Robins, & L. A. Pervin (Eds.), *Handbook of Personality: Theory and Research* (3rd ed., pp.114–158). New York: Guilford.

Mõttus, R., Briley, D.A., Mann, F.D., Tackett, J.L., Harden, K.P., & Tucker-Drob,E. M. (2019). Kids becoming less alike: A behavioral genetic decomposition of developmental increases in personality variance from childhood to adolescence. *Journal of Personality and Social Psychology, 117,* 635-638.

**UK Biobank (UKB)**

The UK Biobank is a large-scale biomedical database and research resource, containing in-depth, de-identified genetic and health information from half a million UK participants. Since 2006, UK Biobank has collected an unprecedented amount of biological and medical data on participants aged between 40 and 69 years old and living in the UK (Bycroft et al., 2018). For this study, we conducted novel analyses of the 12-item Eysenck Personality Questionnaire-revised neuroticism scale (Eysenck et al., 1984) among EUR participants. We restricted analyses to participants who completed nine or more neuroticism items, leading to a final EUR GWAS sample of 441,067 participants ($M_{age}$ = 56.75, SD = 8.02); analyses were conducted in BOLT-LMM to account for participant relatedness (Loh et al., 2015). Neuroticism scores were calculated as the average of item scores a participant responded to. Among AFR participants, we used publicly available neuroticism GWAS summary statistics from the Nealelab pan-ancestry genetic analyses of the UK Biobank (see pan.ukbb.broadinstitute.org for complete information).

Bycroft, C., Freeman, C., Petkova, D., Band, G., Elliott, L. T., Sharp, K., ... & Marchini, J. (2018). The UK Biobank resource with deep phenotyping and genomic data. *Nature*, *562*(7726), 203-209.

Loh, P. R., Tucker, G., Bulik-Sullivan, B. K., Vilhjálmsson, B. J., Finucane, H. K., Salem, R. M., ... & Price, A. L. (2015). Efficient Bayesian mixed-model analysis increases association power in large cohorts. *Nature genetics*, *47*(3), 284-290.

**Dortmund Vital Study (VITAL)**

The aim of the Dortmund Vital Study is to evaluate the effect of several endogenous and exogenous parameters and their complex interaction on cognition and the underlying neuronal activity across the adult life span (Gajewski et al., 2022). This will be achieved by implementing both cross-sectional and longitudinal designs by repeated measurements every 5 years over a long period of time. Cognitive performance in humans is a complex phenomenon that is influenced by numerous variables such as age, infections, inflammatory processes, metabolic parameters, personality, lifestyle factors, type of work, nutrition and stress as well as numerous genetic variants that can be analyzed by polygenic scores. Risk factors for possible cognitive impairment associated with depression, occupational burn-out or age-related diseases, such as mild cognitive impairment or dementia, but also non-pathological cognitive decline shall be established. The total number of participants completed the baseline measure was 610. The final sample with GWAS scores includes 515 healthy individuals (no history of psychological or neurological disorders) aged 20-70 (M = 45.0, SD = 14.2). The male subsample includes 199 individuals (38.6%) aged 20-70 (M = 47.2, SD = 14.5) and the female subsample includes 316 individuals (61.4%) aged 20-70 (M = 43.6, SD = 13.8). On their first day of participation, subjects completed several questionnaires, and underwent a neuropsychological, cardiovascular and electrophysiological testing using PC-based cognitive tasks. On their second day of participation, subjects completed further pen-and-paper tests and completed a second session of cognitive PC-based tests with EEG-recording and MRI scanning. Both sessions took place at Leibniz Research Centre for Working Environment and Human Factors at TU Dortmund, Germany (IfADo). The data were collected between 2016 and 2020. Personality traits were assessed via the German version of the Revised NEO-Personality-Inventory (NEO-FFI), which comprises a total of 60 self-report items to measure the domains neuroticism, extraversion, openness to experience, agreeableness, and conscientiousness.

Gajewski, P.D., Getzmann, S., Bröde, P., Burke, M., Cadenas, C., Capellino, S., Claus, M., Genç, E., Golka, K., Hengstler, J.G., Kleinsorge, T., Marchan, R., Nitsche, M., Reinders, J., van Thriel, C., Watzl, C., & Wascher, E. (2022). Impact of biological and lifestyle factors on cognitive aging and work ability in the Dortmund Vital Study: Protocol of an interdisciplinary, cross-sectional and longitudinal study. *JMIR Research Protocols*, *11*(3), e32352.

#### Wisconsin Longitudinal Study (WLS)

The Wisconsin Longitudinal Study (WLS) is a cohort-based study of people who graduated from Wisconsin high schools in 1957 and one of their randomly selected siblings (Herd et al., 2014). In 1992-1993, 2004-2005, and 2011, participants reported on their personality traits using a 29-item version of the Big Five Inventory (John, 1995), which measures the Big Five traits with 6 questions each (and Neuroticism with 5 questions). In this study, we restricted genetic analyses to the N = 5,867 participants (52% female, 48% male, Mage = 62.85, SDage = 5.21) with personality trait data who were unrelated (IBD < .044) and of European Ancestry (within an ellipse of greatest density of self-reported "white" participants based on the first two PC eigenvectors). The WLS contains 15 sets of identical twins; we randomly selected one twin per pair for genome-wide analyses.

Herd, P., Carr, D., & Roan, C. (2014). Cohort profile: Wisconsin longitudinal study (WLS). *International journal of epidemiology*, *43*(1), 34-41.
John, O. (1995). Big Five Inventory. Berkeley: University of California, Institute of Personality and Social Research.

#### Yale-Penn Study

The Yale-Penn (YP) studies include participants recruited in the eastern United States, predominantly in Connecticut and Pennsylvania. YP participants were administered the 240 NEO Personality Inventory-Revised, which measures each of the Big Five personality traits with 48 items. Each study received IRB approval from all participating institutions and written informed consent was obtained from all study participants. Additional information is available in the previous GWAS publications (Gelernter et al., 2014a; Gelernter et al., 2014b, Gelernter et al., 2014c, Gelernter et al., 2015, Sherva et al., 2016). In this study, the three Yale-Penn contributing samples are referred to as Yale-Penn_1 through Yale-Penn_3 (See **Supplementary Tables S1** and **S2** for complete information).

Gelernter, J., Kranzler, H. R., Sherva, R., Koesterer, R., Almasy, L., Zhao, H., & Farrer, L. A. (2014). Genome-wide association study of opioid dependence: multiple associations mapped to calcium and potassium pathways. *Biological Psychiatry*, *76*(1), 66-74.
Gelernter, J., Sherva, R., Koesterer, R., Almasy, L., Zhao, H., Kranzler, H. R., & Farrer, L. (2014). Genome-wide association study of cocaine dependence and related traits: FAM53B identified as a risk gene. *Molecular Psychiatry*, *19*(6), 717-723.
Gelernter, J., Kranzler, H. R., Sherva, R., Almasy, L., Koesterer, R., Smith, A. H., Anton, R., Preuss, U.W., Ridinger, M., Rujescu, D., Wodarz, N., Zill, P., Zhao, H., & Farrer, L. A. (2014). Genome-wide association study of alcohol dependence: significant findings in African-and European-Americans including novel risk loci. *Molecular Psychiatry*, *19*(1), 41-49.
Gelernter, J., Kranzler, H. R., Sherva, R., Almasy, L., Herman, A. I., Koesterer, R., Zhao, H., & Farrer, L. A. (2015). Genome-wide association study of nicotine dependence in American populations: identification of novel risk loci in both African-Americans and European-Americans. *Biological Psychiatry*, *77*(5), 493-503.
Sherva, R., Wang, Q., Kranzler, H., Zhao, H., Koesterer, R., Herman, A., Farrer, L.A., & Gelernter, J. (2016). Genome-wide association study of cannabis dependence severity, novel risk variants, and shared genetic risks. *JAMA Psychiatry*, *73*(5), 472-480.

#### Young Finns Study (YFS)

The Young Finns Study (YFS) is a prospective multi-centre follow-up study assessing cardiovascular risk factors from childhood to adulthood (Raitakari et al., 2008). The study was initiated in 1980 with 3,596 children and adolescents aged 3–18 years. The participants were randomly selected from the areas of five university hospitals in Finland (Turku, Tampere, Helsinki, Kuopio, and Oulu) and have been followed up for over 40 years. The study was approved by the ethical committee of the Hospital District of Southwest Finland on June 20, 2017 (ETMK:68/1801/2017), and all participants have given an informed written consent. This study involved 1,898 individuals with European ancestry aged 30-45 (M = 37.68, SD = 5.04; 57% female).

We used the NEO-FFI, developed by Rantanen et al. (2007). The Finnish version used in this study contains a total of 60 questions (12 items per trait) that are based on the original NEO-FFI (Costa & McCrae, 1992) and questions from the Finnish version of the NEO-PI (Personality Inventory). Some of the questions in the Finnish Personality Inventory (PI) version were modified to better correspond to non-Indo-European languages. Each personality trait was measured with 12 items (i.e., a total of 60 items in the questionnaire). The NEO-FFI was used twice, in 2007 and in 2012. We first calculated a mean score for year 2007 and for year 2012. Then, we calculated a mean score of the 2007 and 2012 mean scores (for all the participants who had data available in at least one measurement year, i.e., in 2007 and/or 2012). Finally, the mean score of each personality trait was standardized (mean = 0, SD = 1). We allowed max 10% of missing values per trait (i.e., one missing item per trait) in each measurement year. High values of the mean scores indicated high neuroticism, high extraversion, high openness, high conscientiousness, and high agreeableness.

The Young Finns Study has been financially supported by the Academy of Finland: grants 356405, 322098, 286284, 134309 (Eye), 126925, 121584, 124282, 129378 (Salve), 117797 (Gendi), 141071 (Skidi), 349708, 330809, and 338395; the Social Insurance Institution of Finland; Competitive State Research Financing of the Expert Responsibility area of Kuopio, Tampere and Turku University Hospitals (grant X51001); Juho Vainio Foundation; Paavo Nurmi Foundation; Finnish Foundation for Cardiovascular Research ; Finnish Cultural Foundation; The Sigrid Juselius Foundation; Tampere Tuberculosis Foundation; Emil Aaltonen Foundation; Yrjö Jahnsson Foundation; Signe and Ane Gyllenberg Foundation; Diabetes Research Foundation of Finnish Diabetes Association; EU Horizon 2020 (grant 755320 for TAXINOMISIS and grant 848146 for To Aition); European Research Council (grant 742927 for MULTIEPIGEN project); Tampere University Hospital Supporting Foundation, Finnish Society of Clinical Chemistry, the Cancer Foundation Finland; pBETTER4U_EU (Preventing obesity through Biologically and bEhaviorally Tailored inTERventions for you, project number: 101080117); CVDLink (EU grant nro. 101137278), and the Jane and Aatos Erkko Foundation.

Costa, P.T., & McCrae, R.R. (1992). Professional manual for the NEO-PI-R and NEO-FFI. Odessa, FL: Psychological Assessment Resources.

Raitakari, O.T., Juonala, M., Rönnemaa, T., Keltikangas-Järvinen, L., Räsänen, L., Pietikäinen, M., Hutri-Kähönen, N., Taittonen, L., Jokinen, E., Marniemi, J., Jula, A., Telama, R., Kähönen, M., Lehtimäki, T., Åkerblom, H.K., & Viikari, J.S.A. (2008). Cohort profile: The Cardiovascular Risk in Young Finns study. *International Journal of Epidemiology*, *37*(6), 1220–1226.

Rantanen, J., Metsäpelto, R., Feldt, T., Pulkkinen, L., & Kokko, K. (2007). Long-term stability in the big five personality traits in adulthood. *Scandinavian Journal of Psychology*, *48*, 511–518.

#### References not cited in main text


Akingbuwa, W. A., Hammerschlag, A. R., Bartels, M., Nivard, M. G., & Middeldorp, C. M. (2022). Ultra-rare and common genetic variant analysis converge to implicate negative selection and neuronal processes in the aetiology of schizophrenia. *Molecular psychiatry*, *27*(9), 3699-3707.

Aitken, A. C. (1936). IV.—On Least Squares and Linear Combination of Observations. Proceedings of the Royal Society of Edinburgh, 55, 42–48. doi:10.1017/S0370164600014346

Andrae, R., Krämer, M. D., Hopwood, C. J., Denissen, J., Scholz, U., Bocklet, V. V., & Bleidorn, W. (2024) Transactions between Personality Traits and First Sexual Experiences in Adolescence and Emerging Adulthood. *PsyArxiv.*

Arumäe, K., Briley, D., Colodro-Conde, L., Mortensen, E. L., Jang, K., Ando, J., ... & Vainik, U. (2021). Two genetic analyses to elucidate causality between body mass index and personality. *International journal of obesity*, *45*(10), 2244-2251.

Atherton, O. E., Willroth, E. C., Weston, S. J., Mroczek, D. K., & Graham, E. K. (2024). Longitudinal associations among the Big Five personality traits and healthcare utilization in the US. *Social science & medicine*, *340*, 116494.

Bates, D. M. (2010, February). *lme4: Mixed-effects modeling with R*.

Benjamini, Y., & Hochberg, Y. (1995). Controlling the false discovery rate: a practical and powerful approach to multiple testing. *Journal of the Royal statistical society: series B (methodological)*, *57*(1), 289-300.

Borenstein, M., Hedges, L., & Rothstein, D. (2007). Meta-analysis: Fixed effect vs. random effects. Retrieved from http://www.metaanalysis.com/downloads/Meta-analysis_fixed_effect_vs_random_ effects_sv.pdf

Bowden, J., Davey Smith, G., Haycock, P. C., & Burgess, S. (2016). Consistent estimation in Mendelian randomization with some invalid instruments using a weighted median estimator. *Genetic epidemiology*, *40*(4), 304-314.

Bryois, J., Skene, N. G., Hansen, T. F., Kogelman, L. J., Watson, H. J., Liu, Z., ... & Sullivan, P. F. (2020). Genetic identification of cell types underlying brain complex traits yields insights into the etiology of Parkinson's disease. *Nature genetics*, *52*(5), 482-493.

Burgess, S., Smith, G. D., Davies, N. M., Dudbridge, F., Gill, D., Glymour, M. M., ... & Theodoratou, E. (2023). Guidelines for performing Mendelian randomization investigations: update for summer 2023. *Wellcome open research*, *4*, 186.

Caille, P., Stephan, Y., Sutin, A. R., Luchetti, M., Canada, B., Heraud, N., & Terracciano, A. (2024). Personality and change in physical activity across 3–10 years. *Psychology & health*, *39*(5), 670-690.

Chang, C. C., Chow, C. C., Tellier, L. C., Vattikuti, S., Purcell, S. M., & Lee, J. J. (2015). Second-generation PLINK: rising to the challenge of larger and richer datasets. *Gigascience*, *4*(1), s13742-015.

Clapp Sullivan, M. L., Schwaba, T., Harden, K. P., Grotzinger, A. D., Nivard, M. G., & Tucker-Drob, E. M. (2024). Beyond the factor indeterminacy problem using genome-wide association data. *Nature human behaviour*, *8*(2), 205-218.

Costa, P. T., & McCrae, R. R. (2008). The revised neo personality inventory (neo-pi-r). *The SAGE handbook of personality theory and assessment*, *2*(2), 179-198.

De Moor, M. H., Costa, P. T., Terracciano, A., Krueger, R. F., De Geus, E. J., Toshiko, T., ... & Boomsma, D. I. (2012). Meta-analysis of genome-wide association studies for personality. *Molecular psychiatry*, *17*(3), 337-349.

De Moor, M. H., Van Den Berg, S. M., Verweij, K. J., Krueger, R. F., Luciano, M., Vasquez, A. A., ... & Genetics of Personality Consortium. (2015). Meta-analysis of genome-wide association studies for neuroticism, and the polygenic association with major depressive disorder. *JAMA psychiatry*, *72*(7), 642-650.

DeYoung, C. G. (2015). Cybernetic big five theory. *Journal of research in personality*, *56*, 33-58.

Digman, J. M. (1997). Higher-order factors of the Big Five. *Journal of personality and social psychology*, *73*(6), 1246.

Dowle, M., Srinivasan, A., Gorecki, J., Chirico, M., Stetsenko, P., Short, T., ... & Tan, X. (2019). Package 'data. table'.

Durbin, C. E., & Hicks, B. M. (2014). Personality and psychopathology: A stagnant field in need of development. *European journal of personality*, *28*(4), 362-386.

Eysenck, H. J., & Eysenck, S. B. G. (1984). Eysenck personality questionnaire-revised.

Ennis, G., Williams, C., Mallard, T. T., Schwaba, T., de la Fuente, J., & Tucker-Drob, E. M. (2025). Genomic taxometric analysis of negative emotionality and major depressive disorder highlights a gradient of genetic differentiation across the severity spectrum. *MedRxiv*

Grosz, M. P., Rohrer, J. M., & Thoemmes, F. (2020). The taboo against explicit causal inference in nonexperimental psychology. *Perspectives on psychological science*, *15*(5), 1243-1255.

Grotzinger, A. D., de la Fuente, J., Privé, F., Nivard, M. G., & Tucker-Drob, E. M. (2023). Pervasive downward bias in estimates of liability-scale heritability in genome-wide association study meta-analysis: a simple solution. *Biological psychiatry*, *93*(1), 29-36.

Grotzinger, A. D., Rhemtulla, M., de Vlaming, R., Ritchie, S. J., Mallard, T. T., Hill, W. D., ... & Tucker-Drob, E. M. (2019). Genomic structural equation modelling provides insights into the multivariate genetic architecture of complex traits. *Nature human behaviour*, *3*(5), 513-525.

Hahn, E., Gottschling, J., & Spinath, F. M. (2012). Short measurements of personality–Validity and reliability of the GSOEP Big Five Inventory (BFI-S). *Journal of research in personality*, *46*(3), 355-359.

Hartwig, F. P., Davey Smith, G., & Bowden, J. (2017). Robust inference in summary data Mendelian randomization via the zero modal pleiotropy assumption. *International journal of epidemiology*, *46*(6), 1985-1998.

Hemani, G., Zheng, J., Elsworth, B., Wade, K. H., Haberland, V., Baird, D., ... & Haycock, P. C. (2018). The MR-Base platform supports systematic causal inference across the human phenome. *elife*, *7*, e34408.

Hopwood, C. J., Schwaba, T., Wright, A. G., Bleidorn, W., & Zanarini, M. C. (2022). Longitudinal associations between borderline personality disorder and five-factor model traits over 24 years. *European journal of personality*, *36*(1), 72-90.

Jang, K. L., McCrae, R. R., Angleitner, A., Riemann, R., & Livesley, W. J. (1998). Heritability of facet-level traits in a cross-cultural twin sample: support for a hierarchical model of personality. *Journal of personality and social psychology*, *74*(6), 1556-1565.

Jiang, L., Zheng, Z., Qi, T., Kemper, K. E., Wray, N. R., Visscher, P. M., & Yang, J. (2019). A resource-efficient tool for mixed model association analysis of large-scale data. *Nature genetics*, *51*(12), 1749-1755.

John, O. P., Donahue, E. M., & Kentle, R. (1991). The Big Five Inventory. Technical report, University of California, Berkeley

John, O. P., Naumann, L. P., & Soto, C. J. (2008). Paradigm shift to the integrative big five trait taxonomy. *Handbook of personality: Theory and research*, *3*(2), 114-158.

Jokela, M., Alvergne, A., Pollet, T. V., & Lummaa, V. (2011). Reproductive behavior and personality traits of the Five Factor Model. *European journal of personality*, *25*(6), 487-500.

Kenny, D. A., & West, T. V. (2010). Similarity and agreement in self-and other perception: A meta-analysis. *Personality and social psychology review*, *14*(2), 196-213.

Koellinger, P. D., Okbay, A., Kweon, H., Schweinert, A., Karlsson Linnér, R., Goebel, J., ... & Hertwig, R. (2021). Cohort Profile: Genetic data in the German Socio-Economic Panel Innovation Sample (Gene-SOEP). *bioRxiv*, 2021-11.

Koning, I. M., Van den Eijnden, R. J., Engels, R. C., Verdurmen, J. E., & Vollebergh, W. A. (2011). Why target early adolescents and parents in alcohol prevention? The mediating effects of self-control, rules and attitudes about alcohol use. *Addiction*, *106*(3), 538-546.

Lassi, G., Taylor, A. E., Timpson, N. J., Kenny, P. J., Mather, R. J., Eisen, T., & Munafò, M. R. (2016). The CHRNA5–A3–B4 gene cluster and smoking: from discovery to therapeutics. *Trends in neurosciences,* 39(12), 851-861.

Lee, K., & Ashton, M. C. (2004). Psychometric properties of the HEXACO personality inventory. *Multivariate behavioral research*, *39*(2), 329-358.

Lucas, R. E. (2023). Why the cross-lagged panel model is almost never the right choice. *Advances in Methods and practices in psychological science*, *6*(1), 25152459231158378.

Luciano, M., Hagenaars, S. P., Davies, G., Hill, W. D., Clarke, T. K., Shirali, M., ... & Deary, I. J. (2018). Association analysis in over 329,000 individuals identifies 116 independent variants influencing neuroticism. *Nature genetics*, *50*(1), 6-11.

Lüdtke, O., Roberts, B. W., Trautwein, U., & Nagy, G. (2011). A random walk down university avenue: life paths, life events, and personality trait change at the transition to university life. *Journal of personality and social psychology*, *101*(3), 620-637.

Machiela, M. J., & Chanock, S. J. (2015). LDlink: a web-based application for exploring population-specific haplotype structure and linking correlated alleles of possible functional variants. *Bioinformatics*, *31*(21), 3555-3557.

McCrae, R. R., Terracciano, A., & 78 Members of the Personality Profiles of Cultures Project. (2005). Universal features of personality traits From the observer's perspective: Data from 50 cultures. *Journal of personality and social psychology, 88*(3), 547–561.

Morrison, J., Knoblauch, N., Marcus, J. H., Stephens, M., & He, X. (2020). Mendelian randomization accounting for correlated and uncorrelated pleiotropic effects using genome-wide summary statistics. *Nature genetics*, *52*(7), 740-747.

Munafo, M. R., Zetteler, J. I., & Clark, T. G. (2007). Personality and smoking status: A meta-analysis. *Nicotine & tobacco research*, *9*(3), 405-413.

Nagel, M., Jansen, P. R., Stringer, S., Watanabe, K., De Leeuw, C. A., Bryois, J., ... & Posthuma, D. (2018). Meta-analysis of genome-wide association studies for neuroticism in 449,484 individuals identifies novel genetic loci and pathways. *Nature genetics*, *50*(7), 920-927.

Novick, M. R., Jackson, P. H., & Thayer, D. T. (1971). Bayesian inference and the classical test theory model: Reliability and true scores. *Psychometrika*, *36*(3), 261-288.

Ormel, J., Jeronimus, B. F., Kotov, R., Riese, H., Bos, E. H., Hankin, B., ... & Oldehinkel, A. J. (2013). Neuroticism and common mental disorders: Meaning and utility of a complex relationship. *Clinical psychology review*, *33*(5), 686-697.

Peters, H., Götz, F. M., Ebert, T., Müller, S. R., Rentfrow, P. J., Gosling, S. D., ... & Matz, S. C. (2023). Regional personality differences predict variation in early COVID-19 infections and mobility patterns indicative of social distancing. *Journal of personality and social psychology*, *124*(4), 848-872.

Purcell, S., Neale, B., Todd-Brown, K., Thomas, L., Ferreira, M. A., Bender, D., ... & Sham, P. C. (2007). PLINK: a tool set for whole-genome association and population-based linkage analyses. *The American journal of human genetics*, *81*(3), 559-575.

R Core Team (2023). _R: A Language and Environment for Statistical Computing_. R Foundation for Statistical Computing, Vienna, Austria.

Revelle, W. R. (2017). psych: Procedures for personality and psychological research.

Sanderson, E., Rosoff, D., Palmer, T., Tilling, K., Smith, G. D., & Hemani, G. (2024). Bias from heritable confounding in Mendelian randomization studies. *medRxiv*, 2024-09.

Saucier, G., Thalmayer, A. G., Payne, D. L., Carlson, R., Sanogo, L., Ole-Kotikash, L., ... & Zhou, X. (2014). A basic bivariate structure of personality attributes evident across nine languages. *Journal of Personality*, *82*(1), 1-14.

Spearman, C. (1910). Correlation calculated from faulty data. *British journal of psychology*, *3*(3), 271.

Streit, F., Awasthi, S., Hall, A. S., Niarchou, M., Marouli, E., Babajide, O., ... & Witt, S. H. (2024). Genome-wide association study of borderline personality disorder identifies six loci and highlights shared risk with mental and somatic disorders. *medRxiv*, 2024-11.

Thalmayer, A. G., Saucier, G., & Eigenhuis, A. (2011). Comparative validity of brief to medium-length Big Five and Big Six Personality Questionnaires. *Psychological assessment*, *23*(4), 995-1009.

The 1000 Genomes Project Consortium. (2015). A global reference for human genetic variation. *Nature,* 526, 68–74.

Thorgeirsson, T. E., Geller, F., Sulem, P., Rafnar, T., Wiste, A., Magnusson, K. P., ... & Stefansson, K. (2008). A variant associated with nicotine dependence, lung cancer and peripheral arterial disease. *Nature*, *452*(7187), 638-642.

Tucker-Drob, E. M., & Briley, D. A. (2014). Continuity of genetic and environmental influences on cognition across the life span: a meta-analysis of longitudinal twin and adoption studies. *Psychological bulletin*, *140*(4), 949.

Turkheimer, E., Pettersson, E., & Horn, E. E. (2014). A phenotypic null hypothesis for the genetics of personality. *Annual review of psychology*, *65*(1), 515-540.

Turner, S. D. (2014). qqman: an R package for visualizing GWAS results using QQ and manhattan plots. *Biorxiv*, 005165.

Turner, S., Armstrong, L. L., Bradford, Y., Carlson, C. S., Crawford, D. C., Crenshaw, A. T., ... & Ritchie, M. D. (2011). Quality control procedures for genome-wide association studies. *Current protocols in human genetics*, *68*(1), 1-19.

Van den Berg, S. M., de Moor, M. H., Verweij, K. J., Krueger, R. F., Luciano, M., Arias Vasquez, A., ... & Boomsma, D. I. (2016). Meta-analysis of genome-wide association studies for extraversion: findings from the genetics of personality consortium. *Behavior genetics*, *46*, 170-182.

Vazire, S. (2010). Who knows what about a person? The self–other knowledge asymmetry (SOKA) model. *Journal of personality and social psychology*, *98*(2), 281-300.

Watanabe, K., Taskesen, E., Van Bochoven, A., & Posthuma, D. (2017). Functional mapping and annotation of genetic associations with FUMA. *Nature communications*, *8*(1), 1826.

Wei, T., Simko, V., Levy, M., Xie, Y., Jin, Y., & Zemla, J. (2017). Package 'corrplot'. *Statistician*, *56*(316), e24.

Wickham, H., Averick, M., Bryan, J., Chang, W., McGowan, L. D. A., François, R., ... & Yutani, H. (2019). Welcome to the Tidyverse. *Journal of open source software*, *4*(43), 1686.

Willroth, E. C., Beck, E., Yoneda, T. B., Beam, C. R., Deary, I. J., Drewelies, J., ... & Graham, E. K. (2025). Associations of personality trait level and change with mortality risk in 11 longitudinal studies. *Journal of personality and social psychology*.

Winkler, T. W., Day, F. R., Croteau-Chonka, D. C., Wood, A. R., Locke, A. E., Mägi, R., ... & Genetic Investigation of Anthropometric Traits (GIANT) Consortium. (2014). Quality control and conduct of genome-wide association meta-analyses. *Nature protocols*, *9*(5), 1192-1212.

Wu, X. R., Li, Z. Y., Yang, L., Liu, Y., Fei, C. J., Deng, Y. T., ... & Yu, J. T. (2024). Large-scale exome sequencing identified 18 novel genes for neuroticism in 394,005 UK-based individuals. *Nature human behaviour*, 1-14.

Yang, J., Lee, S. H., Goddard, M. E., & Visscher, P. M. (2011). GCTA: a tool for genome-wide complex trait analysis. *The american journal of human genetics*, *88*(1), 76-82.

Young, A. I., Nehzati, S. M., Benonisdottir, S., Okbay, A., Jayashankar, H., Lee, C., ... & Kong, A. (2022). Mendelian imputation of parental genotypes improves estimates of direct genetic effects. *Nature genetics*, *54*(6), 897-905.
